# The circadian clock gene circuit controls protein and phosphoprotein rhythms in *Arabidopsis thaliana*

**DOI:** 10.1101/760892

**Authors:** Johanna Krahmer, Matthew Hindle, Laura K Perby, Tom H Nielsen, Karen J Halliday, Gerben van Ooijen, Thierry Le Bihan, Andrew J Millar

## Abstract

24-hour, circadian rhythms control many eukaryotic mRNA levels, whereas the levels of their more stable proteins are not expected to reflect the RNA rhythms, emphasizing the need to test the circadian regulation of protein abundance and modification. Here we present circadian proteomic and phosphoproteomic time-series from *Arabidopsis thaliana* plants under constant light conditions, estimating that just 0.4% of quantified proteins but a much larger proportion of quantified phospho-sites were rhythmic. Approximately half of the rhythmic phospho-sites were most phosphorylated at subjective dawn, a pattern we term the ‘phospho-dawn’. Members of the SnRK/CDPK family of protein kinases are candidate regulators. A *CCA1*-over-expressing line that disables the clock gene circuit lacked most circadian protein phosphorylation. However, the few phospho-sites that fluctuated despite *CCA1*-over-expression still tended to peak in abundance close to subjective dawn, suggesting that the canonical clock mechanism is necessary for most but perhaps not all protein phosphorylation rhythms. To test the potential functional relevance of our datasets, we conducted phosphomimetic experiments using the bifunctional enzyme fructose-6-phosphate-2-kinase / phosphatase (F2KP), as an example. The rhythmic phosphorylation of diverse protein targets is controlled by the clock gene circuit, implicating post-translational mechanisms in the transmission of circadian timing information in plants.

## Introduction

Most circadian observations in the well-characterised plant species *Arabidopsis thaliana* can be explained by a genetic network of mostly negatively interacting transcription factors (1, 2). In addition to transcriptional interactions, this transcriptional-translational feedback loop (TTFL) system requires post-translational modification of transcription factor proteins (3). Phosphorylation of CCA1 by casein kinase (CK) 2 for example is necessary for the circadian clock function (4).

Protein phosphorylation is involved in the circadian clock mechanism not only in plants but also in fungi, animals and cyanobacteria (3, 5). While the transcription factors of TTFLs in animals, fungi and plants are not evolutionarily conserved (3), many kinases that play a role in circadian timekeeping are similar across eukaryotes. For instance CK2 is also important for circadian timing in mammals (6) and fungi (7). Protein phosphorylation is also involved in the output of the circadian clock (8). However, only one study has so far addressed the question of how pervasive circadian protein phosphorylation is in higher plants (9).

In circadian biology, transcriptional studies have long dominated research efforts, leading to the well-established TTFL models (e.g. refs (1,2,10–12)). However, it has become apparent that protein abundance and post-translational modification cannot be ignored, since these do not simply follow transcript expression patterns (e.g. refs. (13–15)). There is even evidence that circadian oscillations can be driven by non-transcriptional oscillators (NTOs) that are independent of rhythmic transcription. The cyanobacterial circadian clock is based on rhythmic autophosphorylation of the KaiC protein together with the KaiA and KaiB proteins and this mechanism does not even require a living cell to create oscillations (16). Evidence for NTOs also exists in eukaryotes; the protein peroxiredoxin (PRX) is rhythmically over-oxidized in the absence of transcription in algae and human red blood cells (17, 18). Circadian rhythms of PRX over-oxidation were also observed in organisms that have impaired circadian oscillators, in mutants of the fungus *Neurospora crassa* and in transgenic Arabidopsis plants (19). The circadian PRX over-oxidation rhythm even exists in cyanobacteria and archaea (19). In addition, circadian magnesium and potassium ion transport has been observed across eukaryotes and can occur in transcriptionally inactive *Ostreococcus tauri* and human red blood cells (20, 21). Therefore, at least some eukaroytes possess NTOs that appear to be evolutionarily ancient and conserved (3).

With mass spectrometer technology becoming more and more advanced, several circadian proteomics studies have been conducted in different species, such as proteomics analyses of protein abundance time courses (15,22–25), proteomics specifically at the day / night transition (25, 26) or circadian phosphoproteomics (9, 24). In this study, we used mass spectrometry based proteomics and phosphoproteomics on circadian time courses to address the following questions: (1) How pervasive are rhythms in protein abundance and phosphorylation as a clock output in a normally functioning circadian clock system, and what are the characteristics of such rhythms? and (2) can protein abundance or phosphorylation be rhythmic in a plant with a disabled transcriptional oscillator? To investigate (1), we used a time course of WT plants, and for addressing (2), we generated time courses from plants overexpressing the *CIRCADIAN CLOCK-ASSOCIATED 1* gene (CCA1-OX), which have an impaired TTFL. We generated global proteomics and phosphoproteomics data in parallel from the same protein extracts. Our analysis revealed that the transcriptional oscillator is required for most rhythmic protein phosphorylation, and that most rhythmic phosphopeptides peak at subjective dawn. We also found this ‘phospho-dawn’ trend among the time courses of fluctuating phosphopeptides in the CCA1-OX. Finally, we selected a phosphosite of the bifunctional enzyme fructose-6-phosphate-2-kinase / fructose-2,6-bisphosphatase (F2KP) to illustrate how our data can be used to study the mechanisms of clock output pathways that connect to central carbon metabolism.

## Experimental Procedures

### Plant material

*Arabidopsis thaliana* WT (Col-0 accession) and a *CCA1* over-expressing plants (‘CCA1-OX’, (27)) were used in this study. Seeds were germinated and grown on plates (2.15g/l Murashige & Skoog medium Basal Salt Mixture (Duchefa Biochemie), pH 5.8 (adjusted with KOH) and 12g/l agar (Sigma)) at 85µmol m^-2^ s^-1^ white fluorescent lights at 21°C in Percival incubators for 11 days in 12h light, 12h dark cycles. Seedlings were transferred to soil and grown for 11 more days at a light intensity of 110µmol m^-2^ s^-1^ in the same light-dark cycle.

### Experimental Design and Statistical Rationale

After plants had grown for a total of 22 days, from ZT 0 of day 23 lights remained switched on continuously and collection of plant material started at ZT 12 (dataset I) or ZT 24 (dataset II). In dataset I, six time points at 4h intervals were taken with five replicates for each time point, harvesting eight rosettes for the WT and 12 rosettes for the CCA1-OX. In dataset II, also at 4h intervals, at least six replicates of eight rosettes each were taken for the WT, five (16 rosettes each) of the CCA1-OX. In dataset I, the time course was sampled from ZT12 to ZT32, in dataset II from ZT24 to 48 (CCA1-OX) or ZT24 to ZT52 (WT).

Statistical analysis of time courses required assessment of not only changes but also rhythmicity. We therefore used both analysis of variance (ANOVA) and JTK_CYCLE as statistical tools (see below for details).

### Sample preparation

Rosettes without roots were harvested by flash-freezing in liquid nitrogen. Protein extraction and precipitation was carried out according to method ‘IGEPAL-TCA’ described by (28). Briefly, protein was extracted and precipitated with TCA and phase separation, then washed with methanol and acetone. 500μg re-suspended protein was digested using a standard in-solution protocol and peptides were desalted. Before drying, eluted peptides were separated into two parts: 490μg of the digest was used for phosphopeptide enrichment, 10μg was saved for global protein analysis. Phosphopeptides were enriched using the Titansphere™ spin tip kit (GL Sciences Inc.) and desalted on BondElut Omix tips (Agilent) according to the manufacturers’ instructions.

### Mass spectrometry, peptide merging and Progenesis analysis

LC-MS/MS measurement and subsequent analysis by the Progenesis software (version 4.1.4924.40586) was carried out as previously described (28): Dried peptides were dissolved in 12 μl (phosphoproteomics) or 20µl (global proteomics) 0.05% TFA and passed through Millex-LH 0.45μm (Millipore) filters. 8 μl were run on an on-line capillary-HPLC-MSMS system consisting of a micropump (1200 binary HPLC system, Agilent, UK) coupled to a hybrid LTQ-Orbitrap XL instrument (Thermo-Fisher, UK). Reverse phase buffer used for LC-MS separation was buffer A (2.5% acetonitrile, 0.1% FA in H_2_O) and buffer B (10% H_2_O, 90% acetonitrile, 0.1% formic acid, 0.025% TFA). LC peptide separation was carried out on an initial 80 min long linear gradient from 0% to 35% buffer B, then a steeper gradient up to 98% buffer B over a period of 20 min then remaining constant at 98% buffer B for 15 min until a quick drop to 0% buffer B before the end of the run at 120 min.

The tair Arabidopsis_1rep (version 20110103, 27416 protein entries) database was used for data-dependent detection, using the Mascot search engine (version 2.4), including all peptide sequences of rank smaller than 5. Search parameters were as follows: charges 2+, 3+ and 4+, trypsin as enzyme, allowing up to two missed cleavages, carbamidomethyl (C) as a fixed modification, Oxidation (M), Phospho (ST) and Phospho (Y), Acetyl(Protein N-term) as variable modifications, a peptide tolerance of 7 ppm, and MS/MS tolerance of 0.4 Da, peptide charges 2+, 3+, and 4+, on an ESI-trap instrument, with decoy search and an ion-cutoff of 20. In all but one cases, these parameters resulted in a false-discovery rate (FDR), of less than 5% with one exception (phosphoproteomics dataset I: 3.5%, phosphoproteomics dataset II: 3.2, global dataset I: 6.8%, global dataset II: 4.5%, calculated using the formula 2*d/(n+d) (29), n and d being the number of hits in the normal and decoy databases, respectively, using an ion score cutoff of 20). Peptides were quantified by their peak area by Progenesis, and proteins were quantified by using the sum of the quantitation of all unique peptides. Where peptides matched very similar proteins, multiple accession numbers are shown in exported results from Progenesis (Supplementary Data S1, S2).

In order to remove duplicates of phosphopeptides due to alternative modifications other than phosphorylation or missed cleavages, we used the qpMerge software following the Progenesis analysis (30). The data are publicly available in the pep2pro database (31) at http://fgcz-pep2pro.uzh.ch (Assembly names ‘ed.ac.uk Global I’, ‘ed.ac.uk Global II’, ‘ed.ac.uk Phospho I’, ‘ed.ac.uk Phospho II’) and have been deposited to the ProteomeXchange Consortium (http://proteomecentral.proteomexchange.org) via the PRIDE partner repository (32) with the dataset identifier PXD009230. Exported .csv files from Progenesis with all peptide and protein quantifications can be found in the online supplementary material (Data S1 and S2).

### General statistics and outlier removal

Results of statistical analyses are summarised in Data S3. Outlier analysis, statistics and Venn diagrams were done using R version 3.2.1 (33). Zero values for the quantification were exchanged for ‘NA’. For outlier analysis and parametric tests such as ANOVA, arcsinh transformed data was used to obtain a normal distribution, while untransformed data was used for plotting time courses and for the non-parametric JTK_CYCLE analysis. For phosphoproteomics analysis all replicates in which the Pearson correlation coefficient among replicates of the same time point was lower than 0.8 were regarded as outliers (supplementary methods, Supplementary Data S4). In global dataset II, the first run replicate of each time point had to be excluded as an outlier due to an apparent drift (Supplementary Data S4). To generate heat maps, the abundance of each peptide or protein was normalized by the time course mean of the peptide or protein, followed by taking the log2 to centre values around 0, and the heatmap.2 function from the pvclust R package was applied (34).

### JTK_CYCLE analysis

We used the R-based tool JTK_CYCLE (35) to determine rhythmicity, with the following modifications: (1) we ran the JTK_CYCLE algorithm for each phosphosite or protein separately rather than the entire list, to allow handling of missing quantification values for some replicates. Benjamini Hochberg (BH) (36) correction of p-values was carried out after application of the JTK_CYCLE tool. (2) Since our time course durations are close to the periods of rhythms we are searching for, some identifications were assessed as rhythmic by the original JTK_CYCLE tool that were increasing or decreasing continually over the entire time course. We excluded these from the group of rhythmic identifications (‘excl.’ in p-value column in Supplementary Data S5). For dataset-wide analyses we used p<0.05 as a cutoff for rhythmicity and then trusted results that agree between experiments. Similarly, for judging individual time courses we focus on those with p < 0.05 in both datasets.

### Kinase prediction using GPS3.0

Kinases for each site were predicted using the *Arabidopsis thaliana* specific GPS 3.0 prediction tool in its species specific mode for *Arabidopsis thaliana* (37, 38). For each phosphorylation site, an amino acid sequence was generated that contained 50 amino acids on either side of the phosphorylated residue using a python script. This resulted in 101 amino acid long sequences, unless the phosphosite was closer than 50 residues to the C or T terminus in which case the missing positions were filled by ‘X’. The phosphopeptides used as foreground were the significantly rhythmic phosphosites (JTK_CYCLE p-value < 0.05) peaking at a given time point, all other phosphosites identified in the same experiment were used as background. The high threshold setting was used to minimize false-positive predictions and searches were done for S and T residues. In order to reduce the complexity of the dataset, we used a simplification: where kinases from different families were predicted, only the one with the highest difference between score and cutoff was used. For foreground and background the numbers of predictions for each occurring kinase group were counted and the Fisher’s Exact test was used to determine predicted kinases that were significantly enriched in the foreground group (p<0.05).

### GO analysis

For GO analysis, foreground and background were chosen as in the kinase prediction analysis. With these groups, GO analysis was conducted using the topGO script (39), followed by Fisher’s exact test to determine enrichment of terms (supplementary data S6).

### Generation of F2KP point mutations and expression constructs

The F2KP coding sequence in the pDONR221 vector, lacking a stop codon, was kindly provided by Dr Sebastian Streb, ETH Zürich. The QuikChange Lightning Site-Directed Mutagenesis kit (Agilent Technologies) was used to introduce point mutations using primer pair AspF and AspR for mutation of S276 to aspartic acid or AlaF and AlaR for mutation to alanine (Table S 4). The WT or mutated F2KP coding sequences were amplified by PCR using primers F2KP-F and F2KP-R (Table S 4), introducing restriction sites for AflII (3’ end) and XbaI (5’ end) and a stop codon. Using these restriction sites, digested PCR products were ligated into the pmcnEAVNG expression vector which allows expression in *E.coli* with an N-terminal GST tag and a T7 promoter for IPTG inducibility. Plasmids were transformed into Rosetta™2(DE3)pLysS Competent expression strain (Novagen).

### Expression of GST-F2KP constructs in E. coli cells

The three constructs, WT, S276E and S276A were expressed in *E.coli* and purified using the GST tag. Two independent expression experiments were performed (experiment 1 and experiment 2), each in triplicates. 200ml *E.coli* cultures were grown at 37°C with 100μg/ml ampicillin and 25μg/ml chloramphenicol and induced with 1mM IPTG at OD_600_ values between 0.6 and 0.8. Cells were harvested by centrifugation after 2.5h of expression at 37°C. Each pellet was from 50ml bacterial culture. For purification of GST-F2KP, pellets were lysed in 2.5ml PBS with Complete protease inhibitor cocktail (Roche) with a probe sonicator. After clearing of the lysates, AP was carried out using GSH-agarose beads, with 167µl GSH-agarose bead suspension (Protino glutathione agarose 4B, Macherey-Nagel), a binding incubation of 30min at room temperature, 4 washes with 10x the volume of the bead suspension and elution in PBS with 100mM reduced glutatione (pH 8.0) 3 times 30min at room temperature.

### F2KP activity assay

F2KP activity of purified F2KP was measured as described in (40) (F-2,6-BP producing reaction) and (41) (measurement of generated F-2,6-BP by its activation of PFP and subsequent production of NADH produced from glycolytic enzymes).

### Western blot quantification of F2KP concentration in AP eluates

To test whether the differences in F2KP activity of eluates could be caused by differences in abundance in the eluate, we quantified the amount of F2KP in equal volumes of eluates by western blotting. Samples were prepared for SDS-PAGE with 25% 4xLDS (Life Technologies, NP0008) and 20mM DTT and were incubated at 70°C for 10min. Two concentration series of an equal mix of all three eluates were loaded on each gel for relative quantitation. Individual eluates were examined in duplicates (expression 1) or triplicates (expression 2). 4-12% Bis-Tris Gels (Life Technologies) were run and protein was blotted onto nitrocellulose membrane using the iBlot**^®^** system. Two primary antibodies were used in parallel: anti-F2KP from rabbit, raised against amino acids 566-651 (42) and anti-GST from mouse (Thermo Scientific), both at a dilution of 1:1,000 overnight. Secondary antibodies were goat-anti-rabbit (IRDye®800CW, LI-COR) and goat-anti-mouse (IRDye®680RD, LI-COR). Bands were quantified using the ImageStudioLite (version 2) software.

## Results

### Circadian protein phosphorylation requires the canonical transcriptional oscillator

We generated global proteomics and phosphoproteomics datasets for two independent circadian time courses and for two genotypes each – WT and the CCA1-OX line which has an impaired circadian oscillator (27). This resulted in four datasets: global protein and phosphopeptide datasets I (Zeitgeber time (ZT) 12 to ZT32) and datasets II (ZT24 to ZT48 (CCA1-OX) or ZT52 (WT)) (Figure 1A,B). We removed outliers before conducting further statistical analysis (Supplementary Data S4).

**Figure 1:**
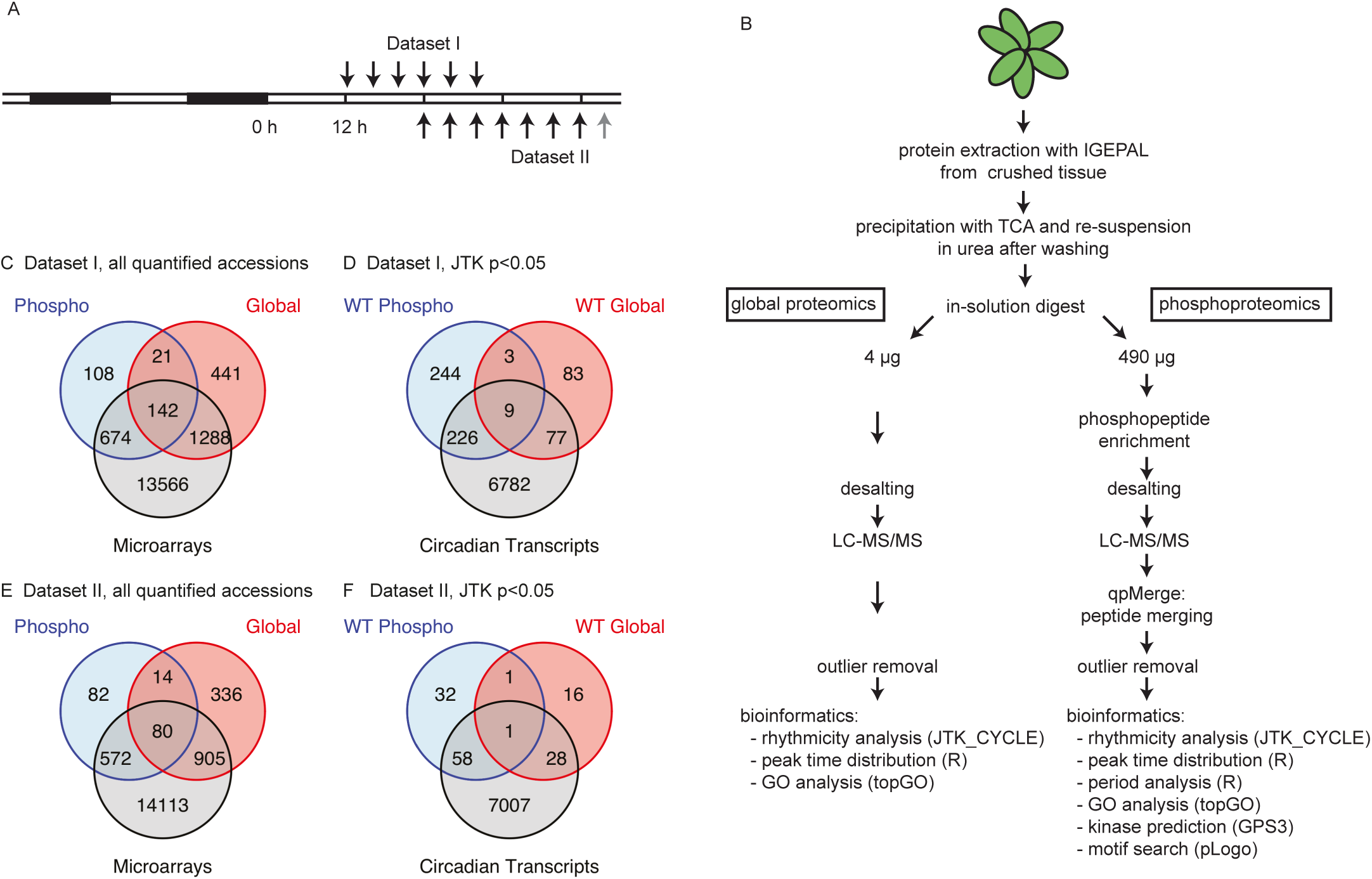
Experiment workflow and comparison of protein and phosphopeptide numbers with published transcriptome time courses. A) WT and CCA1-OX plants were grown in 12h light:12h dark cycles for 22 days and then subjected to continuous light. In dataset I, rosettes were harvested every 4h from ZT12, in dataset II from ZT24 until ZT48 (black arrows) with an additional WT time point at ZT52 (grey arrow). B) Sample processing and data analysis workflow: Plants were crushed and protein was extracted, precipitated and digested in-solution. Peptides were split into 10µg for global protein analysis (of which 4µg were injected) and 490µg for phosphopeptide enrichment on TiO_2_ spin tips. Peptides were analysed by LC-MS/MS. Outliers were removed before further bioinformatics analysis. C) Venn diagrams showing overlap of quantified (C,E) and rhythmic (WT only; D,F) transcripts (Covington et al. 2008), proteins and phosphoproteins in dataset I (C,D) and dataset II (E,F).

We identified 2287 phosphopeptides in dataset I, 1664 in dataset II, which condensed to between 1000 and 1500 in each dataset after applying qpMerge (30) to remove duplicate phosphopeptides (Table 1A). These were from several hundred proteins in each dataset and over 1000 in both datasets together (Table 1B). In the global protein analysis, we identified 1896 and 1340 proteins in dataset I and II, respectively, adding up to a total of 2501 for both datasets combined (Table 1A,B). To assess the circadian rhythmicity of each phosphopeptide or protein, we employed the non-parametric JTK_CYCLE method (35) as it can be applied to time courses of only one cycle, taking the curve shape into account. Unless otherwise stated we considered periods of 22 to 26h and excluded continuously increasing or decreasing profiles from the group of rhythmic phosphopeptides or proteins. In dataset I, 606 (40%) phosphopeptides were rhythmic in the WT, in dataset II 100 (8.8%) based on the p-value of the individual timeseries. 338 (23%) (dataset I) and 26 (2.3%) (dataset II) were rhythmic after adjusting for multiple testing (‘q-value’ < 0.05) (36). The fraction of rhythmic proteins in the global proteomics analysis was smaller than in the case of phosphopeptides: 171 (9.0%) in dataset I, 45 (3.4%) in dataset II had JTK_CYCLE p-values < 0.05. 6 proteins also had q-value < 0.05 in each dataset (0.32% in dataset I, 0.45% in dataset II). Phosphopeptides and proteins with JTK_CYCLE p<0.05 in both datasets are listed in Table 2.

**Table 1:** Identification counts for global and phosphoproteomics datasets. A) Numbers of quantified and changing identifications in each dataset, B) shared and added numbers of both datasets. In B) protein IDs of peptides are used for simplicity of comparison.

| A | Dataset I |  |  |  | Dataset II |  |  |  |
| --- | --- | --- | --- | --- | --- | --- | --- | --- |
|  | Global Proteomics |  | Phosphoproteomics |  | Global Proteomics |  | Phosphoproteomics |  |
|  | WT | CCA1-OX | WT | CCA1-OX | WT | CCA1-OX | WT | CCA1-OX |
| Quantifiable identifications before merging | 1896 proteins |  | 2287 phosphopeptides |  | 1340 proteins |  | 1664 phosphopeptides |  |
| Quantifiable identifications after merging |  |  | 1498 from 944 proteins |  |  |  | 1132 from 747 proteins |  |
| Significantly changing (ANOVA $p < 0.05$) | 245 | 147 | 406 | 296 | 68 | 46 | 88 | 83 |
| Adjusted for multiple testing (ANOVA $q < 0.05$) | 13 | 3 | 56 | 4 | 3 | 2 | 14 | 7 |
| Rhythmic by JTK_CYCLE ($p < 0.05$) | 171 | 122 | 606 from 481 proteins | 37 from 32 proteins | 45 | 57 | 100 from 91 proteins | 17 from 17 proteins |
| Adjusted for multiple testing (JTK_CYCLE $q < 0.05$) | 6 | 0 | 338 | 2 | 6 | 0 | 26 | 0 |
| JTK_CYCLE $p < 0.05$ in both genotypes | 17 | | 22 | | 9 | | 3 | |

| B | Global Proteomics |  | Phosphoproteomics |  |
| --- | --- | --- | --- | --- |
|  | WT | CCA1-OX | WT | CCA1-OX |
| JTK $p < 0.05$ detected in both datasets | 6 | 3 | 48 | 2 |
| all quantified protein IDs detected in both datasets | 137 |  | 626 |  |
| sum of all quantifiable protein IDs in both datasets | 2501 |  | 1065 |  |

**Table 2:**
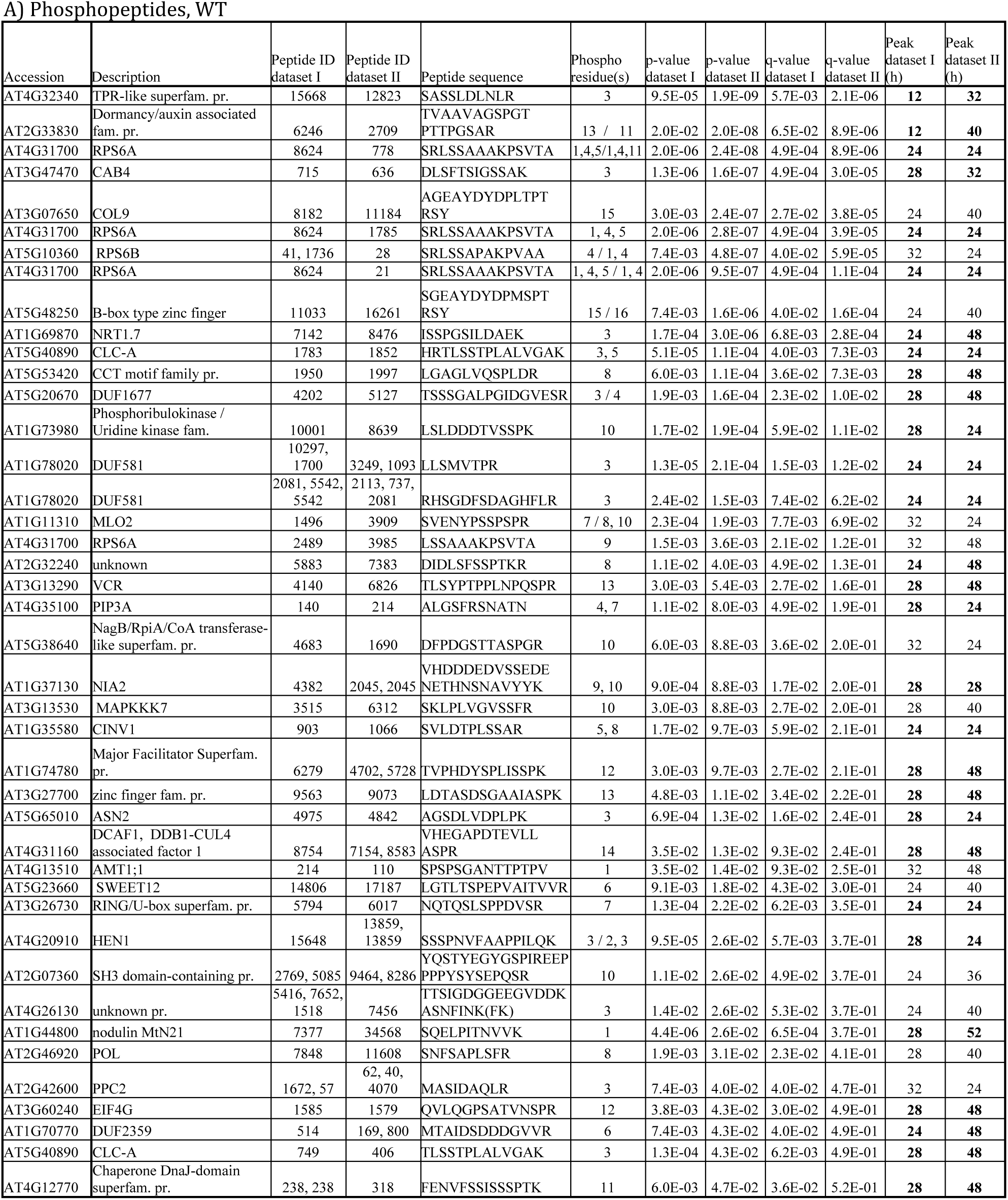
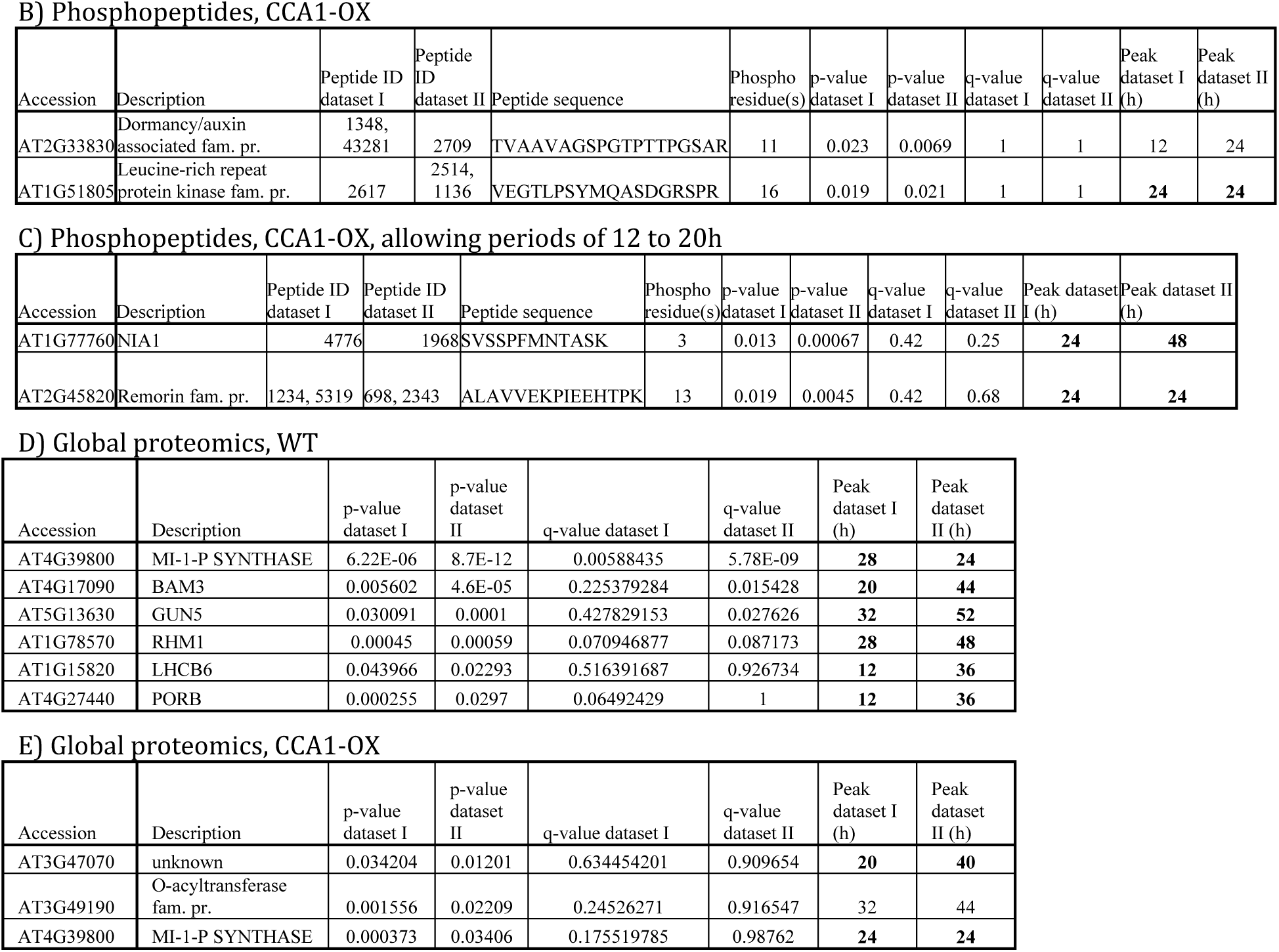
List of all proteins and phosphopeptides that are rhythmic in WT or CCA1-OX in both datasets (i.e. shared rhythmic IDs). All p-values are based on JTK_CYCLE analysis. * in some phosphopeptides, the location of phosphorylated residues differs slightly (either shifted by 1-2 residues or one of several phosphates is missing) between datasets, in which case the phosphorylated residues of both datasets are noted (dataset I / dataset II). Bold: phase difference of peak is up to 4h. fam. = family, pr. = protein

In the CCA1-OX line, 37 (2.5%) phosphopeptides had a JTK_CYCLE p-value < 0.05 in dataset I, 17 (1.5%) in dataset II; in the global datasets, 122(6.4%) and 57 (4.3%) had a JTK_CYCLE p-value < 0.05 in datasets I and II, respectively. After adjusting for multiple testing, only two significant identifications remained for the CCA1-OX phosphopeptides in dataset I and none in dataset II (Table 1A). This analysis suggests that a functional TTFL is required for most rhythmic protein phosphorylation.

For further whole-dataset analyses, we used JTK_CYCLE p-value < 0.05 as a criterion for rhythmicity and treated results as reliable if rhythmic scores were obtained from separate analysis of each dataset. This approach is also supported by comparison with existing data: Among the phosphopeptides with p < 0.05 and q>0.05 were phosphosites that were previously shown to be rhythmic with almost identical phases, such as RTT(pS)LPVDAIDS of WITH NO LYSINE (WNK) 1, and TL(pS)STPLALVGAK of CHLORIDE-CHANNEL-A (CLC-A) (Supplementary Data S3) (9).As expected, the global proteomic analysis did not quantify all the proteins identified by the phosphoproteomic enrichment (Figure 1 C,E). We found very few proteins with rhythms in abundance as well as rhythmic phosphopeptides (12 in dataset I, 2 in dataset II). About half of the rhythmic phosphopeptides or proteins had rhythmic transcripts (Figure 1D,F).

Since *CCA1* is a morning-expressed gene, the CCA1-OX line might be expected to have a ‘morning-locked’ circadian oscillator. To test whether this observation applies at the protein abundance and phosphorylation level, we calculated the absolute value of the difference between CCA1-OX and WT (**|**CCA1-OX - WT**|**) at each dawn and dusk time point (i.e. ZT12, 24, 36 and 48) for time courses of all proteins or phosphopeptides that were rhythmic in WT (p-value <0.05) and quantified in CCA1-OX. For each dawn – dusk pair, we determined whether the difference between CCA1-OX and WT was larger at dusk or at dawn and counted the number of such pairs as a coarse indication of the difference between the proteome or phosphoproteome of these two genotypes, for each dataset and time point pair (Table S 1). CCA1-OX differed more from the WT at dusk rather than dawn, in all but one of the time point pairs (the exception was one of the smallest pairs, in global dataset II). This is consistent with a partially morning-locked circadian clock at the protein (modification) level, as expected from the role of CCA1 in the TTFL. The consistency of the dataset supports our interpretation, from the very few rhythmic identifications in CCA1-OX, that the TTFL is necessary for most of the rhythms observed in WT plants.

### Circadian rhythms of proteins in the global proteomics datasets

We applied GO analysis to the global proteomics data, using rhythmic proteins at each peak time point as foreground and all other identified proteins as background (Supplementary Data S6). In both datasets, enriched GO terms in the WT at the end of the subjective day (ZT12 and ZT36) were photosynthesis related terms, ‘response to glucose’, ‘regulation of protein dephosphorylation’ and oxidoreductase activity with NAD or NADP as acceptor. The latter was also enriched in the CCA1-OX in dataset I. In addition, in the CCA1-OX, two terms related to cell wall metabolic processes were enriched in both datasets during the subjective night (ZT20 and ZT44) (Supplementary Data S6).

Among rhythmic proteins shared between the datasets, we identified six for the WT, three for the CCA1-OX (Table 2D,E). One protein, INOSITOL 3-PHOSPHATE SYNTHASE 1 (MIPS1, AT4G39800) was rhythmic with a peak at 24-28h in both datasets and both genotypes but with lower p-value and higher amplitude in the WT (Figure 2A). Two other examples of high-confidence rhythms in protein abundance are chloroplast BETA-AMYLASE 3 (BAM3) (Figure 2B) and the light-harvesting chlorophyll a/b binding protein LHCB2.2 (Figure 2C).

**Figure 2:**
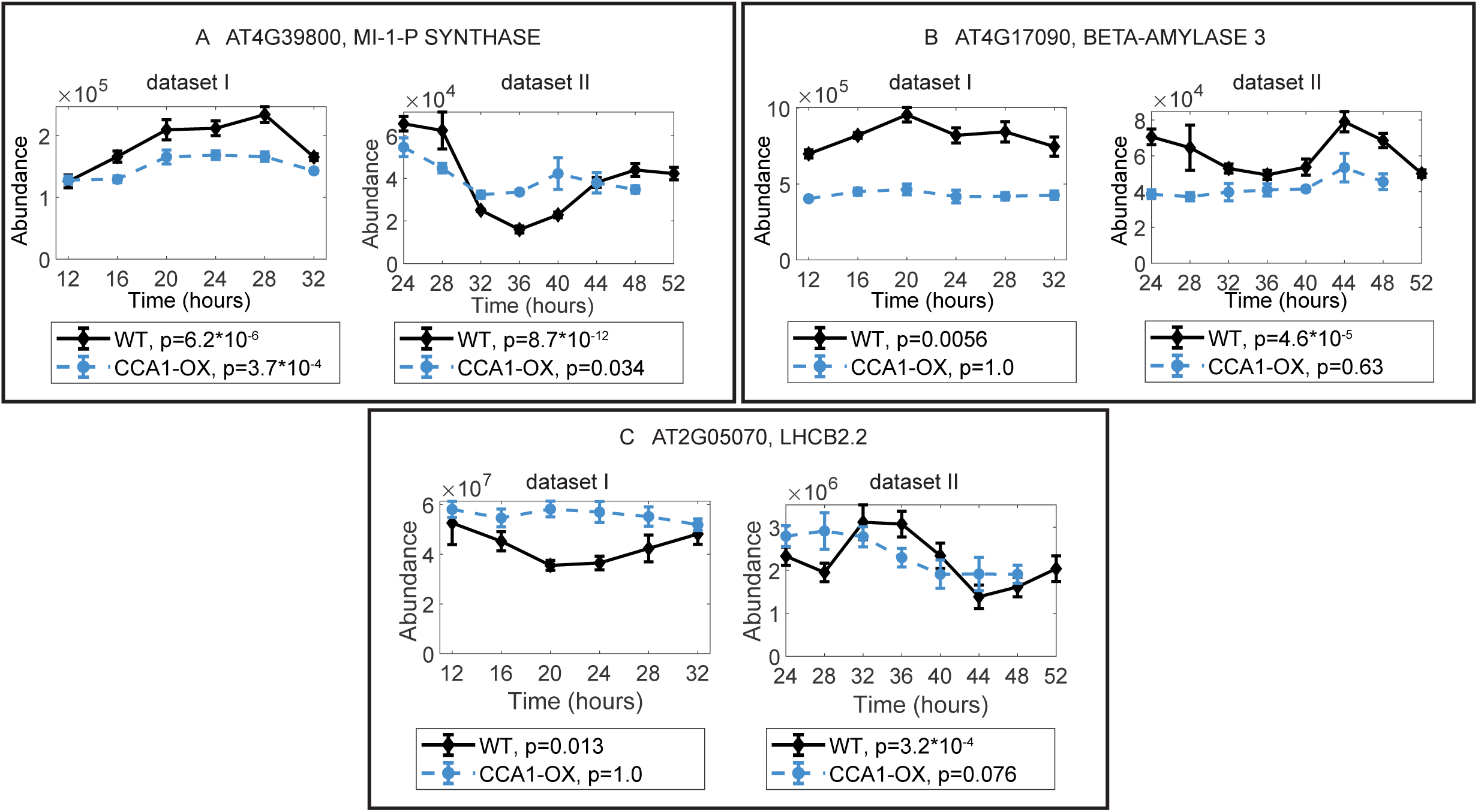
Examples of three proteins with rhythmic abundance in both datasets. (A) MIPS1, (B) BAM3, (C) LHCB2.2. JTK_CYCLE p-values are indicated under the graphs.

### WT phosphoproteomics data reveals rhythmic phosphoproteins with a variety of functions, including previously unknown phosphosite rhythms

GO term enrichment within the peak time groups of the phosphoproteomics data revealed that only one GO term (‘cotyledon development’) was shared between the two datasets and the ZTs of enrichment are 16 h apart (Supplementary Data S6). Several terms were shared between the WT and the CCA1-OX in dataset I, most of them related to energy metabolism or ion homeostasis, and all of them were enriched at ZT24 or ZT28. Apart from overlap of exact GO IDs, we found enrichment of terms related to translation in the WT at 24h in both phosphoproteomics datasets, which is consistent with rhythmic phosphorylation of RPS6 isoforms (Supplementary data S6, Table 2, Data S3) (9).

In agreement with the small number of consistently enriched GO terms and with (9), we found that the proteins for which we found rhythmic phosphosites in the WT are associated with a large variety of functions, such as translation initiation (RPS6A, RPS6B), nitrogen / amino acid metabolism or transport (NIA2, NRT1.7, CLC-A), light harvesting (CAB4) or flowering (COL-9).

In our datasets we also found previously unknown phosphosite rhythms, such as on ASPARAGINE SYNTHETASE (ASN) 2, SWEET12, PLASMA MEMBRANE INTRINSIC PROTEIN (PIP) 2;7 and VARICOSE RELATED (VCR) (Figure 3, and see discussion).

**Figure 3:**
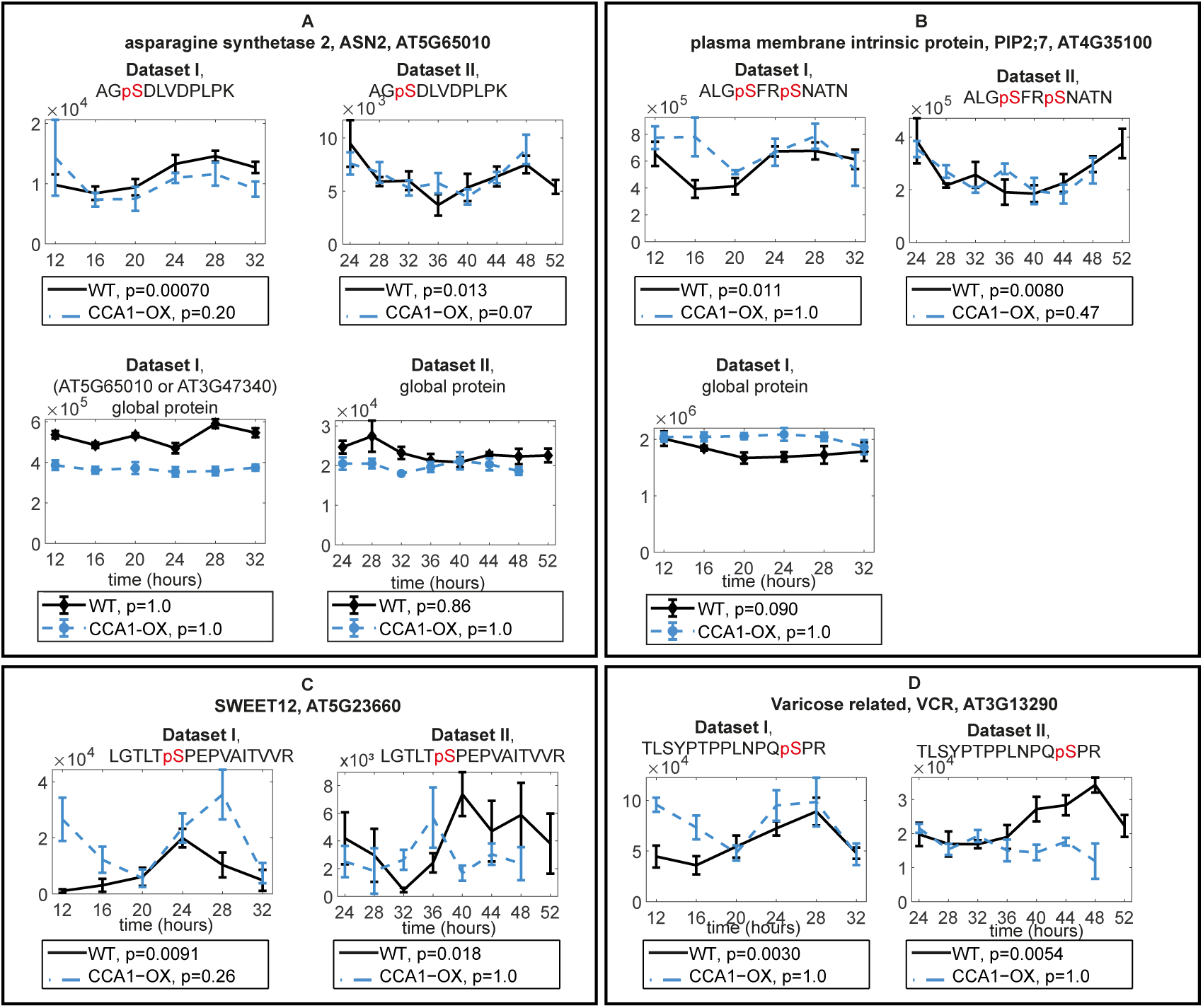
Examples of newly described phosphopeptide rhythms with JTK_CYCLE p<0.05 in both datasets. Phosphopeptide from (A) an asparagine synthetase, ASN2, with global protein abundance plots (B) from aquaporin PIP2;7 with global protein abundance plot from dataset I (not detected in dataset II) (C) from the sucrose efflux protein SWEET12 and (D) from VCR. Protein not detected in global proteomics for (C) and (D). JTK_CYCLE p-values are indicated under the graphs.

### Very few proteins are rhythmically phosphorylated in the CCA1-OX

Only 2 phosphopeptides with p<0.05 for the CCA1-OX appear in both datasets (Table 2B): A phosphopeptide of a Leucine rich repeat protein kinase (AT2G33830) and a dormany/auxin associated family protein (AT1G51805). The latter showed a significant increase in its total protein abundance in dataset I (Table 2B), therefore changes may be due to increasing protein expression in constant light.

### ‘Phospho-dawn’: most rhythmic phosphopeptides peak in the subjective morning

Analysis of the number of phosphopeptides that peak at each time point revealed that 45% (dataset I) and 73% (dataset II) of rhythmic phosphopeptides peak around subjective dawn in the WT (Figure 4A, Figure 5A,B). This is in agreement with previous observations in *Ostreococcus* and *Arabidopsis* (9, 43). By contrast, in the global proteomics dataset no tendency for increased abundance at dawn was observed (Figure 4C, Figure 5C,D). These observations hold true when using an alternative ANOVA analysis, detecting change rather than rhythmicity, with a p-value cutoff of p<0.05 (Figure S 1). Therefore, the preponderance of ‘phospho dawn’ patterns is more likely due to (de)phosphorylation events rather than to changes in the abundance of the cognate proteins.

**Figure 4:**
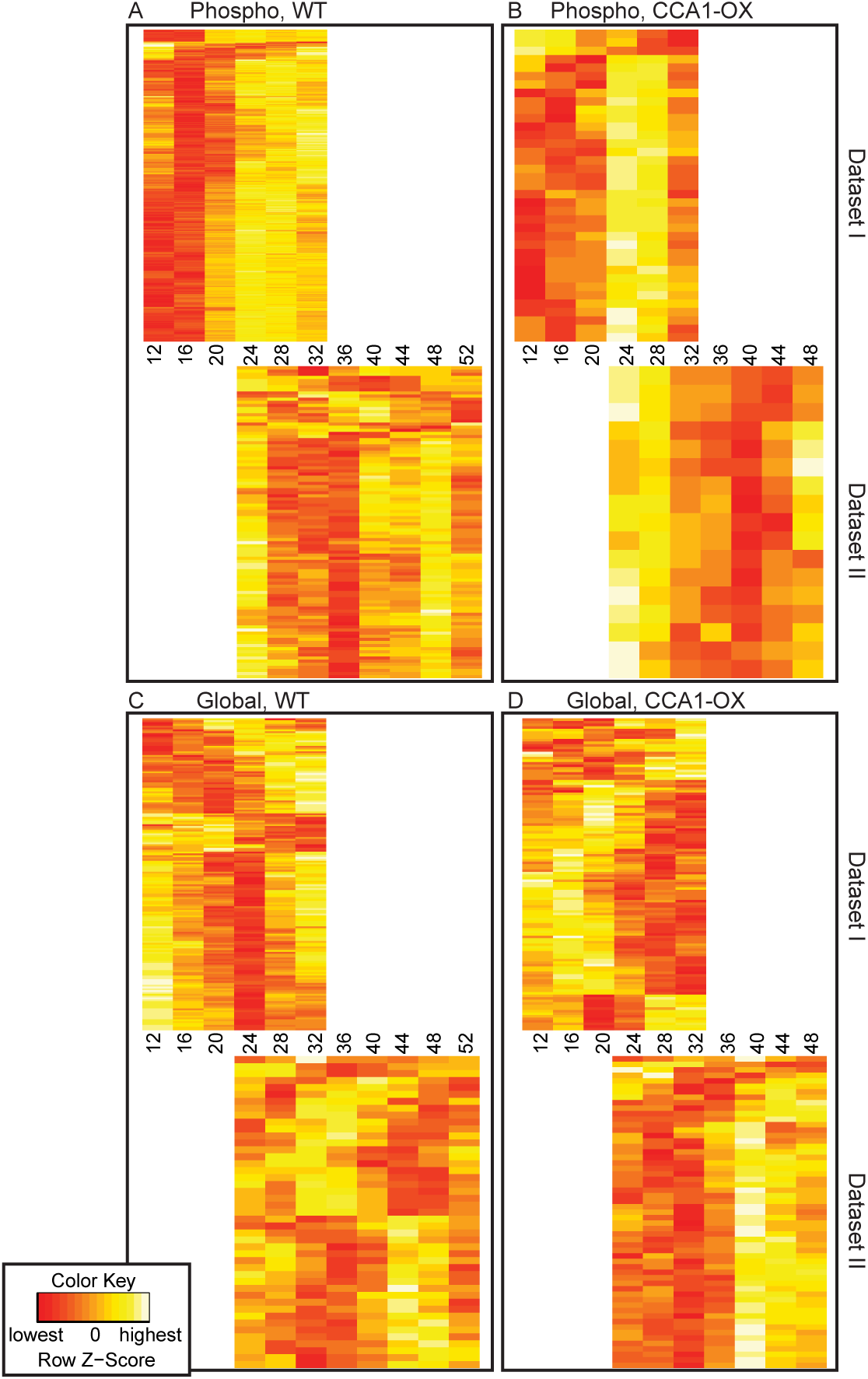
Whole-dataset protein and phosphopeptide dynamics. Heatmaps were generated by hierarchical clustering of phosphopeptide (A, B) or global protein (C, D) abundance time courses in dataset I and II (indicated on the right) for WT (A,C) and CCA1-OX (B,D).

**Figure 5:**
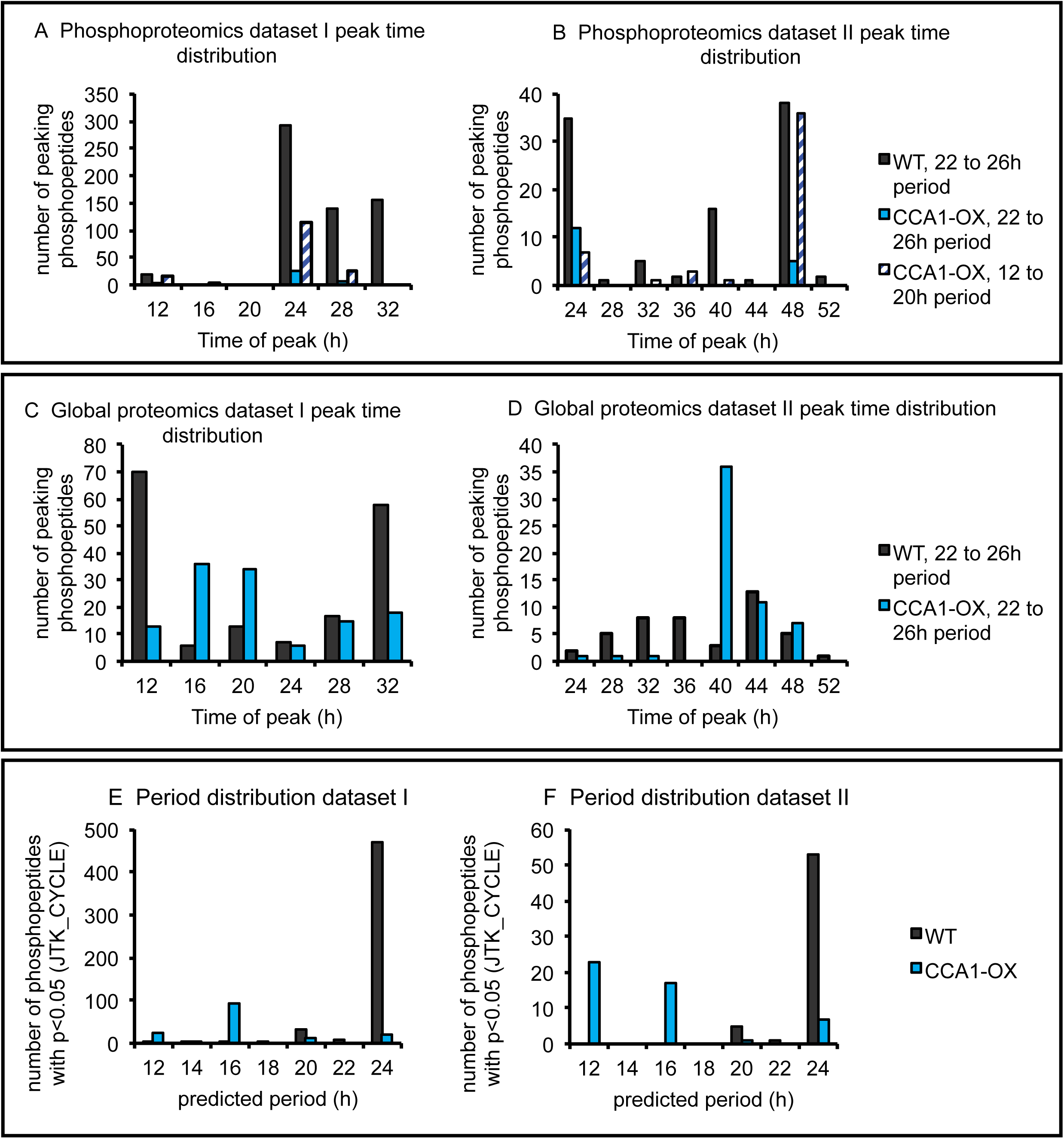
Peak time and period distribution of phosphopeptide and global protein time courses. (A-D): Number of rhythmic (JTK_CYCLE p-value <0.05) phosphopeptides (A,B) or proteins (C,D) peaking at each time point, allowing a period of around 24h (22 to 26) or 12 to 20h. (E-F) Periods according to JTK_CYCLE allowing periods from 12 to 24h for phosphoproteomics dataset I (E) and II (F). Number of rhythmic phosphopeptides (JTK_CYCLE p-value < 0.05) for each predicted period is plotted.

Interestingly, almost all of the few phosphopeptides with JTK_CYCLE p-value < 0.05 also peaked at subjective dawn in the CCA1-OX plants (Figure 4B, Figure 5A,B, Figure S 1), which suggests residual rhythmicity phased similarly to the WT. To expand the search for rhythms in the CCA1-OX, we tested whether there are rhythmic phosphopeptides with shorter periods that would have been excluded from the analysis above. We repeated the JTK_CYCLE analysis, allowing periods down to 12 h. Hardly any phosphopeptides had periods of less than 22 h in the WT, while the majority of phosphopeptides had a predicted period of 12 or 16 h in the CCA1-OX (Figure 5E,F). Interestingly, the majority of those short-period rhythmic phosphopeptides also peaked at 24 h or 48 h (Figure 5A,B). In conclusion, in both of our independent WT phosphoproteomics datasets, the majority of phosphopeptides peak around subjective dawn, and this ‘phospho dawn’ may not be completely abolished by disruption of the TTFL.

### Kinase prediction suggests CDPK/SnRK family members target dawn phased phosphopeptides

We reasoned that there may be a kinase activity that is present predominantly around subjective dawn which is either very robustly dawn-timed by the TTFL even when it is strongly impaired, or an alternative oscillator, such as an NTO, contributes to the dawn phased kinase activity. Characterisation of the phospho-dawn peptides could help to identify either a very robust dawn-phased TTFL output, or potentially consequences of an NTO. For this reason, we focused on the dawn peaking phosphopeptides to identify candidate kinases, using phosphosite motif analysis and kinase prediction. In these analyses we used the ZT24 or ZT48 peaking rhythmic (JTK_CYCLE p-value <0.05) phosphopeptides as foreground and all other identified phosphopeptides as background. No target site motifs were significantly over-represented in a consistent way between datasets (Figure S 2, Figure S 3). For predicting candidate kinases, we searched for enrichment of kinase groups that target phosphopeptides with JTK_CYCLE p-value <0.05 (Table 3) using the GPS3 resource. In the WT, both datasets share enrichment of CMGC kinase groups such as MAPK, and CAMK groups. The latter caught our attention since it is the only consistently enriched group in both genotypes and both datasets. The CAMK group was also consistently enriched among significantly changing phosphopeptides scored using ANOVA p-value < 0.05 (Table S 2).

**Table 3:** Summary of GPS3 kinase prediction followed by Fisher’s Exact test. Fisher’s exact test p-values for enrichment of each kinase group are shown. a) dataset I, b) dataset II. Foreground groups were chosen with JTK_CYCLE p-values <0.05 and peaks or troughs at indicated ZTs.

| time point | WT, 22 to 26h period |  |  |  | CCA1-OX, 22 to 26h period |  |  | CCA1-OX, 12 to 20h period |
| --- | --- | --- | --- | --- | --- | --- | --- | --- |
|  | peak |  | trough | all p<0.05 | peak | trough | all p<0.05 | peak |
|  | 24h | 28h | 12h |  | 24h | 12h |  | 24h |
| AGC |  |  |  |  | 0.043 |  |  |  |
| AGC/NDR |  | 0.0072 |  |  |  |  |  |  |
| Atypical/TAF1 | 6.50E-06 |  | 2.50E-06 |  |  |  |  |  |
| CAMK |  | 0.0012 |  | 0.0040 |  | 0.00063 | 0.0090 | 0.024 |
| CAMK/DAPK |  |  |  |  |  |  |  | 0.029 |
| CMGC/CK2 | 0.0012 |  | 4.70E-06 |  |  |  |  |  |
| CMGC/MAPK |  |  |  | 0.027 |  |  |  |  |
| Other / ULK |  |  | 0.040 |  |  | 0.0030 | 0.0066 |  |
| Other/WNK |  |  |  |  |  |  |  | 0.044 |

B) dataset II
| time point | WT, 22 to 26h period |  |  | CCA1-OX, 22 to 26h period |  | CCA1-OX, 12 to 20h period |
| --- | --- | --- | --- | --- | --- | --- |
|  | peak |  | all p<0.05 | peak | all p<0.05 | peak |
|  | 24h | 48h |  | 24h |  | 24h |
| AGC | 1.4E-06 |  |  |  |  |  |
| AGC/PDK1 | 0.0049 |  |  |  |  |  |
| CAMK | 0.00024 |  | 0.0049 |  |  | 0.048 |
| CAMK-L |  |  |  |  |  | 0.0035 |
| CAMK-Unique | 0.057 |  |  |  |  |  |
| CMGC | 0.0061 |  |  | 0.040 |  |  |
| CMGC/GSK | 0.0078 |  |  |  |  |  |
| CMGC/MAPK |  | 0.040 |  |  |  |  |
| Other/PEK |  |  |  | 0.028 | 0.047 |  |
| STE/STE7 | 0.037 |  |  |  |  |  |

The CAMK group in plants contains the CDPK/SnRK family of kinases with 89 members (44) in the EKPD database (45) that informs GPS3. Interestingly, among the phospho dawn peptides we found phosphosites that may be direct or indirect SnRK1 target proteins according to two proteomics studies (46, 47): NITRATE REDUCTASE (NIA) 1 and 2 and the bifunctional enzyme F2KP (Figure S 4, Table S 3). Interestingly, nitrate reductases have been reported as classical SnRK1 targets in other species (48). In light-dark cycles, NIA1 and NIA2 protein abundances are rhythmic (25), while in our analysis under constant light NIA1 protein was not detected in the global analysis and NIA2 protein abundance changed significantly (Figure S 4B) but not in phase with the phosphosites, suggestive of regulated (de)phosphorylation. Another indication of increased SnRK1 activity at subjective dawn are rhythms in phosphopeptides and abundance of the protein FCS-LIKE ZINC FINGER (FLZ)6 (Figure S 5): transcriptional up-regulation of *FLZ6* by SnRK1 signalling has previously been shown (49, 50). In a datset with WT seedlings in constant light (51), the *FLZ6* transcript peaks 4h before the FLZ6 protein in our dataset.

### Rhythmically phosphorylated kinases and phosphatases

Since kinases and phosphatases themselves can be regulated by phosphorylation, we were interested in rhythmic phosphopeptides of kinases and phosphatases. Identification of rhythmic kinase or phosphatase activities in the WT could help to discover components of clock output pathways that are mediated *via* protein phosphorylation.

For two kinases, CRK8 and AT5G61560, we found rhythmic phosphopeptides in the WT where protein abundance did not oscillate in parallel, indicating that rhythmicity is due to phosphorylation rather than changes in protein abundance (Figure S 6). CRK8 is a member of the CDPK-SnRK1 superfamily (44), To our knowledge no specific functions have been investigated for either of these kinases.

All rhythmically phosphorylated phosphatases in our data are members of the protein phosphatase 2C (PP2C) family (Figure S 7) and were classified as rhythmic only in the WT. PP2C G1 (Figure S 7A) is involved in ABA dependent salt stress response and, in contrast to PP2CAs is a positive ABA signalling regulator (52) but to our knowledge, no reports exist on the functional relevance of its own phosphorylation. AT3G51470 is also a PP2C G family member, was only rhythmic in dataset I and no functional information is available (53). The final PP2C POLTERGEIST (POL, Figure S 7C) is involved in stem cell regulation (54). In the CCA1-OX data, the profiles for the kinases and phosphatases discussed above can show some similarity to the WT pattern but none were classified as rhythmic by JTK_CYCLE. The biochemical mechanisms underlying the relatively robust phosphoprotein rhythms in the WT should prove easier to investigate than any remaining rhythmicity in the CCA1-OX.

### Phospho-null mutation of Ser267 of the enzyme F2KP enhances F6P-2kinase activity in-vitro

As an example of how our datasets can be used to investigate new clock output pathways, we analysed the molecular function of a phosphosite of the bifunctional enzyme F2KP. Several phosphosites were detected in F2KP with only Ser276 showing a circadian rhythm in the WT, in dataset II only but at a very high significance level (Figure S 4C, Supplementary Data S3). One F2KP peptide was detected in the global protein analysis of dataset II. Its changes over the timecourse are not significant and do not parallel the Ser276 phosphopeptide, therefore it is unlikely that the rhythm in Ser276 phosphorylation is caused by changes in F2KP protein abundance.

We tested whether the Ser276 phosphorylation site is relevant for F2KP function, since this site is highly conserved with other plant species (Figure S 8A), and a very specific enzymatic assay has been described (41, 55). Maximum F6P,2K activity was measured of GST-tagged phosphomimetic mutants S276D and S276A, and the unmutated WT control enzyme *in vitro*. Two independent preparations (bacterial expression and purification using a GST tag) were tested to ensure reproducibility. Equivalence of the amounts of expressed F2KP protein in assays was verified by western blotting with two different antibodies (Figure 6C, Figure S 8D). S276A had an approximately 2.5 fold increased activity compared to the unmutated version, while S276D had only slightly increased activity (Figure 6 A,B, Figure S 8B,C,E). The rhythmicity observed at this phosphosite is therefore consistent with a rhythmic input to F2KP function in central carbon metabolism.

**Figure 6:**
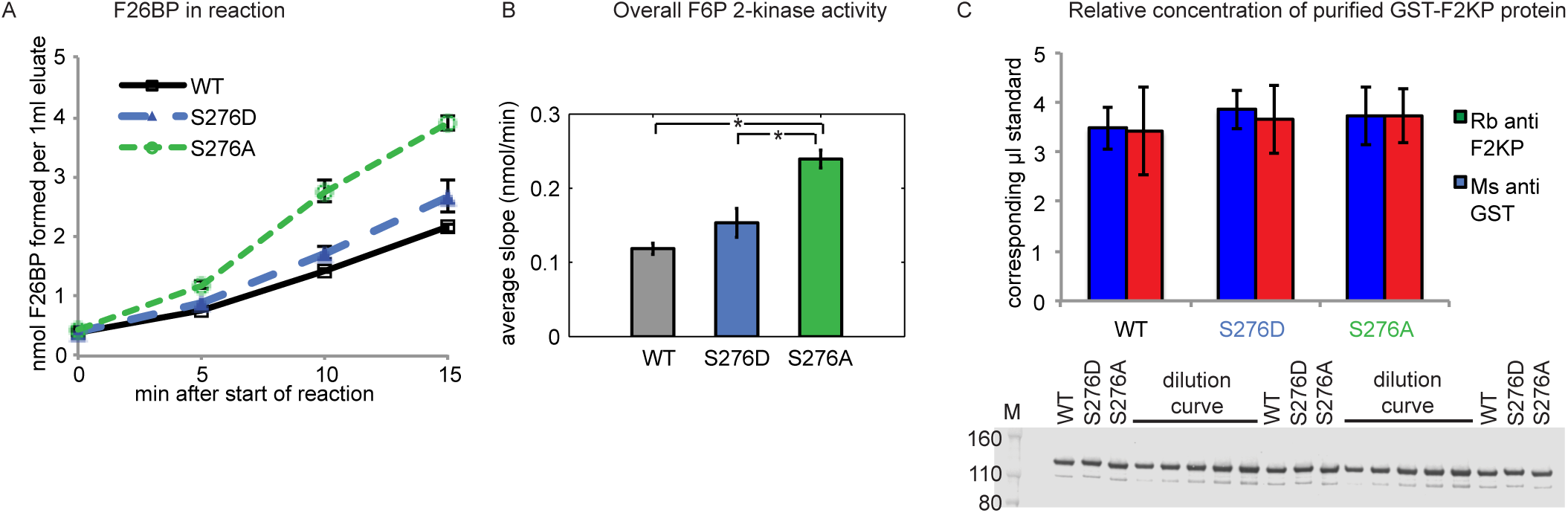
*In vitro* GST-F2KP activity assay with WT and Ser276 point mutations. A) fructose-2,6-bisphosphate (F26BP)) accumulation during the reaction. B) Kinase activity calculated from slopes in (A). C) relative quantification of GST-F2KP in eluates probed with rabbit (Rb) anti F2KP and mouse (Ms) anti GST. Protein blot is shown below quantification for rabbit anti F2KP. A dilution series of a sample mix was used for quantification, ranging from 0.5 to 1.5 loading equivalent of the samples. Averages of both dilution curves were used. Error bars: SEM. * p-value < 0.05 in t-test.

## Discussion

This study focusses on global and phospho-proteomic timeseries in *Arabidopsis* plants harvested under constant light conditions, where rhythmicity is driven by the circadian clock. In WT plants, we tested the prevalence of circadian rhythms in protein phosphorylation and abundance, which has rarely been reported. In the CCA1-OX transgenic line, we tested the importance of the clock gene circuit for that rhythmic regulation, given that markers of a Non-Transcriptional Oscillator (NTO) are rhythmic in Arabidopsis, and that protein phosphorylation drives the best-characterised NTO in cyanobacteria.

### Could an NTO contribute to the phospho-dawn?

CCA1-OX transgenic lines have been widely used as an approximation of a plant without a functional clock. In this line we found very few rhythmic phosphopeptides and proteins, and those with statistical significance typically had less convincing waveforms than rhythms in the WT (Table 1). Therefore, if an NTO exists in *Arabidopsis*, it confers rhythmicity only to very few of the phosphosites we detected. Intriguingly, in both datasets, we still observe the phospho-dawn phenomenon in the CCA1-OX (Figure 4, Figure 5, Figure S 1), whereas a uniform phase distribution would be expected in the noise-driven fluctuations of an arrhythmic plant. It remains possible that an NTO controls this fraction of dawn-phased phosphorylation, albeit too weakly to yield robust rhythms, whereas the majority of phosphoprotein rhythmicity requires a functioning clock gene circuit. We cannot exclude that some rhythmicity of that canonical gene circuit remains even in the CCA1-OX, for example as the CaMV 35S promoter does not confer strong expression in all tissues (56). Additional experiments, such as phosphoproteomics in other arrhythmic mutants, testing other post-translational markers such as redox modifications, and identifying NTO outputs other than PRX over-oxidation (19) and Mg^2+^ rhythms (20, 21) might further define the effects of a potential NTO in *Arabidopsis*.

### Proteins with rhythmic abundance

We found that the proportion of rhythmic phosphopeptides is larger than for protein abundance, but smaller than for rhythmic transcripts (Table 1). The set of rhythmic proteins does not support extensive inference, though we note that the three examples of rhythmic proteins in Figure 2 all follow the rhythmic regulation of their cognate transcripts. MIPS1 (Figure 2A) is transcriptionally induced by light, acting through FAR-RED ELONGATED HYPOCOTYLS (FHY)3 and FAR-RED IMPAIRED RESPONSE (FAR)1 at the transition from darkness to light in light-dark cycles, and enhances myo-inositol abundance, which in turn limits oxidative stress at the onset of photosynthesis (57, 58). Rhythmic *MIPS1* transcript abundance peaks towards the end of the subjective night in constant light (59, 60), before a protein peak shortly after subjective dawn. The rhythmic control of MIPS1 abundance is consistent with anticipation of light-induced oxidative stress, providing a potential physiological function in addition to the subsequent, light-responsive induction of myo-inositol production.

BAM3 (Figure 2B) is the dominant beta-amylase contributing to starch degradation (61). The circadian clock is key for the timing of night-time starch degradation (62). In light-dark cycles, *BAM3* transcript abundance drops at the beginning of the day and increases during the night (63, 64), and this pattern persists in constant light (59, 60). We observe a protein abundance pattern that matches the transcript dynamics, indicating that the transcriptional control may be responsible for the protein rhythm. This is also the case for LHCB2.2 (Figure 2C), a component of the photosystem II light harvesting complex (60).

### Proteins with newly discovered phosphopeptide rhythms

We show four examples of newly discovered phosphopeptide rhythms in Figure 3. ASN2 (Figure 3A) is one out of three described *Arabidopsis* asparagine synthetases which catalyse the transfer of an amino group from glutamine to aspartate, producing asparagine and glutamate, but ASN2 may also directly use ammonia as a substrate (65, 66). The two most expressed asparagine synthetase enzymes in Arabidopsis are ASN1 and ASN2 (65), and in contrast to ASN1, the physiological function of ASN2 is less well understood. We did not find any evidence for ASN2 protein abundance rhythms (Figure 3A), indicating that the phosphosite’s peak near subjective dawn is due to rhythmic kinase and / or phosphatase action. Interestingly, the ammonia transporter AMT1;1 also has a rhythmic phosphosite with a temporal profile that parallels the ASN2 peptide (Data S3, Table 2). The presence of rhythmic phosphosites of proteins involved in nitrate metabolism or transport in our data and (9) supports the notion that nitrogen related processes are under control of the circadian clock at the post-translational level, in part through rhythmic AMT1;1 and ASN2 phosphorylation.

In water transport, our results demonstrate a previously undiscovered phosphosite rhythm on the aquaporin PLASMA MEMBRANE INTRINSIC PROTEIN (PIP)2;7 (Figure 3B). According to (67) this phosphosite is a CPK1 and CPK34 target, and its abundance decreases in response to ABA treatment (68). In response to salt stress, the entire protein is internalised from the plasma membrane, with a concomitant reduction in hydraulic conductivity (69), indicating that decreasing PIP2;7 activity limits water loss. Rhythmic phosphorylation of other aquaporins was previous demonstrated in constant light or darkness (9, 70). Therefore, PIP2;7 may, together with other aquaporins, mediate circadian clock regulation of hydraulic conductivity or high salinity response through its phosphorylation status.

In carbon transport, a newly discovered phosphosite rhythm was found for the sucrose efflux transporter SWEET12 (Figure 3C) (71). To our knowledge, the function of this phosphosite is unknown but one may speculate that this rhythm could reflect circadian control of carbon reallocation.

Finally, we have high confidence in the rhythmicity of a phosphopeptide of the putative RNA decapping protein VARICOSE RELATED (VCR) (Figure 3D). VCR and its close homologue VARICOSE (VCS) interact with and are phosphorylated by SnRK2.6 and SnRK2.10 at several serines (72, 73). While the VCR phosphosite shown in Fig S6D is not one of the phosphosites identified by (73), SnRK2.10 phosphorylates an almost identical site on VCS (TLSYPTPPLNLQpSPR). Therefore, it is very likely that the corresponding site on VRC is also a SnRK2.10 target.

Altogether, these examples demonstrate how our data can be used to generate hypotheses on clock output pathways affecting different aspects of plant physiology through phosphorylation.

### A rhythmic F2KP phosphosite is biochemically relevant

The specific roles of most of the rhythmic phosphorylation sites identified in our study have not been investigated. To exemplify in an experimental approach how circadian phosphorylation of a protein can be linked to its function, we analysed the effect of a phosphosite mutation on the activity of the enzyme F2KP. F2KP is one of the regulators of carbon partitioning into starch and sucrose (74) and is necessary to maintain normal growth in fluctuating light conditions (75). With its kinase domain it can synthesize F-2,6-BP from F-6-P, and with its phosphatase domain it catalyzes the reverse reaction (74).

The phosphosite of interest, Ser276, is within the plant-specific regulatory N-terminal domain (42, 76) but is not among the phosphosites in the known 14-3-3 binding site (77). Ser276 is regulated by SnRK1 (46) and is conserved in many plant species (Figure S 8A).

Our *in-vitro* F-6-P,2 kinase activity measurement experiments showed that substitution of Ser276 by Ala increases F2KP’s kinase activity. It is unknown whether *in vitro* expressed F2KP is phosphorylated at Ser276 or not. However, comparison of Ser276 mutation to Ala with the WT and with mutation to Asp, suggests that a lack of negative charge at position 276 leads to increased kinase activity. In dataset II, pSer276 decreased gradually during the subjective day (Figure S4C). Assuming that the phospho mimic / WT and null mutations reflect the behaviour of the phosphoryated and non-phosphorylated site, respectively, we extrapolate that towards the end of the day, more F-2,6-BP is produced and therefore starch synthesis is favoured over UDP-glucose and sucrose synthesis. Indeed, F-2,6-BP levels in the plant increase slowly across the day in short day conditions (77). Testing the function of these mutations *in planta* will be interesting to determine whether this phosphosite has physiological relevance, in addition to biochemical effectiveness.

### Phospho-dawn is likely mediated by several different kinases

We aimed to characterise the phospho-dawn phenomenon as it may point to novel dawn-specific circadian clock output through post-translational mechanisms. Although the striking abundance of dawn-phased phosphopeptides could partly be biased towards the easily detectable or abundant phosphopeptides of our dataset, it is consistent with highest transcript expression of kinases and phosphatases at the end of the night in diel time courses (26, 78). Our kinase prediction revealed enrichment of some CMGC subgroups, such as MAPK, CK2, GSK, DYRK, CDK or DAPK. CK2 is involved in the circadian clock function in Arabidopsis by phosphorylating CCA1 (4, 79). A previous study reported enrichment of predominantly CK2 predictions among significantly changing phosphopeptides (9). Roles for MAPK and GSK have been reported for the circadian clock function in other eukaryotes (80–83). However, the most consistently enriched group of kinases at subjective dawn in our datasets is the CAMK group (Table 3, Table S 2), which comprises the 89 members of the CDPK-SnRK superfamily of kinases.

All of these 89 CDPK-SnRK member are potential candidates for causing the observed phospho-dawn. Not all of these kinases have been studied in much detail, and for the majority of rhythmic phosphopeptides no experimental evidence for kinase specificities exists.

However, making use of literature on existing kinase – target pairs can help to narrow down candidates. For example, as mentioned above, CPK1, CPK34 and likely SnRK2.10 phosphorylate dawn-peaking phosphosites shown in Figure 3. In addition, SnRK and CPK / CDPK kinases can themselves be regulated by phosphorylation (44). CRK8, of which we found a very prominently dawn-peaking rhythmic phosphopeptide (Figure S 6A), is therefore another candidate phospho-dawn kinase.

We also show that several previously reported SnRK1 regulated sites are rhythmic with peaks around subjective dawn including phosphopeptides of F2KP and nitrate reductases NIA1 and NIA2 (Figure S 4). Additional indication comes from the protein FLZ6 which is transcriptionally induced by and interacts with SnRK1 and may serve as a platform for SnRK1 signalling (84). FLZ6 protein abundance and two phosphopeptides were rhythmic in the WT with a peak around subjective dawn (Figure S 5). SnRK1 may be a particularly relevant candidate as its involvement in circadian timing has previously been reported (85–87) and as it is an important metabolic hub. In normal light-dark conditions the morning is associated with profound metabolic changes in plants, such as the transition from using starch to direct photoassimilates, or to the alternative, a starvation response if light intensities remain low while starch is almost depleted.

SnRK1 signaling is regarded as antagonistic to TOR signaling (47). Nevertheless, RPS6A and RPS6B phosphosites that are targets of the TOR signalling kinase S6K, are rhythmically phosphorylated in the WT, and in dataset I also in the CCA1-OX (Table 2, supplementary data S3 and (9)), with peaks at subjective dawn. This adds to growing evidence that the interplay between SnRK1 and TOR may be more complex than simply antagonistic (88). In fact, the above-mentioned SnRK1 induced FLZ6 negatively feeds back to SnRK1, and this has been suggested as a mechanism to allow sufficient TOR activity in spite of high SnRK1 activity (50), which may allow RPS6 phosphorylation to peak at approximately the same time as SnRK1 activity.

Altogether, the identities of the phospho dawn peptides in our study, along with their known and predicted kinases suggest that phospho-dawn is caused not by a single kinase but several members of the SnRK-CDPK family and also potentially kinases outside this family such as S6K. Further experimentation is required to give evidence for involvement of any such kinases in phospho-dawn, such as time courses of kinase activity and time-resolved phosphoproteomics in mutants of specific candidate kinases. Finding mechanisms that connect the canonical oscillator to prominent post-translational changes at dawn could reveal major clock output pathways that may control a wide range of physiological functions and expand our understanding of how the circadian oscillator increases plant fitness.

## Abbreviations

TTFL: Transcriptional translational feedback loop
NTO: non-transcriptional oscillator
F2KP: fructose-6-phosphate-2-kinase / phosphatase
CCA1: Circadian clock associated 1
CCA1: OX CCA1 overexpressor
CK: casein kinase
SNF-1: sucrose non-fermenting
SnRK: SNF-1 related kinase
TOC1: timing of CAB expression 1
TOC1: Ox TOC1 overexpressor
PRX: peroxiredoxin
PCA: principal component analysis
Col-0: Columbia 0
ZT: Zeitgeber time
H: hour
WT: wildtype
GO: gene ontology
SEM: standard error of the mean
BH: Benjamini - Hochberg

## Acknowledgements

We thank Lisa Imrie, Katalin Kis and Helle K Mogensen for expert technical support.

## Data availability

The data are publicly available in the pep2pro database (31) at http://fgcz-pep2pro.uzh.ch (Assembly names ‘ed.ac.uk Global I’, ‘ed.ac.uk Global II’, ‘ed.ac.uk Phospho I’, ‘ed.ac.uk Phospho II’) and have been deposited to the ProteomeXchange Consortium (http://proteomecentral.proteomexchange.org) via the PRIDE partner repository (32) with the dataset identifier PXD009230. Exported .csv files from Progenesis with all peptide and protein quantifications can be found in the online supplementary material (Data S1 and S2).

## Footnotes

This work was supported by the Wellcome Trust [096995/Z/11/Z] and BBSRC, and EPSRC awards [BB/D019621] and [BB/J009423].

