## Supplementary Data S4 for "The circadian clock gene circuit controls protein and phosphoprotein rhythms in *Arabidopsis thaliana*": outlier.R

```
# script for PCA and correlation analysis to determine outliers.
# this is for one the Phospho I dataset for the WT as an example
# the zip file includes this script, and input file for the above dataset, and the two output pdfs.
# user needs to set path.


peptidedata = read.table('outlier_input.csv', skip=4, header=1, sep=",", quote="\"")

peptideStats = peptidedata[grep("(^X3702_PhI_wt).+[^\\.][0-9]$", names(peptidedata))] 

Hours <- factor(c(12,  12,  12,  12,  12,  16,  16,  16,  16,  16,  20,  20,  20,  20,  20,  24,  24,  24,  24,  24,  28,  28,  28,  28,  28,  32,  32,  32,  32,  32))
replicates <-   c("A", "B", "C", "D", "E", "A", "B", "C", "D", "E", "A", "B", "C", "D", "E", "A", "B", "C", "D", "E", "A", "B", "C", "D", "E", "A", "B", "C", "D", "E")

labels = sapply(1:length(Hours), function(x) {paste(as.character(Hours[x]), replicates[x], sep="")})
names(peptideStats) <- labels
row.names(peptideStats) <- peptidedata[,"X."]
peptideStats[peptideStats==0] <- NA
peptideStatsArchsinh <- asinh(peptideStats)
meanReading <- as.data.frame(t(apply(peptideStatsArchsinh, 1, function(x) c(by(x, Hours, function(x) mean(t(na.omit(x))))))))

####PCA on replicates###

datasetrep <- t(na.omit(peptideStatsArchsinh))


fitReplicates <- prcomp(datasetrep, cor=TRUE, scale.=FALSE, center=TRUE)

pdf("outlier_PCA.pdf", width=5, height=5)
plot(fitReplicates$x[,c(1,2)], lwd = 0, cex = 0)
text(fitReplicates$x[,c(1,2)], row.names(datasetrep), cex = 0.6)
plot(fitReplicates$x[,c(2,3)], lwd = 0, cex = 0)
text(fitReplicates$x[,c(2,3)], row.names(datasetrep), cex = 0.6)
plot(fitReplicates$x[,c(3,4)], lwd = 0, cex = 0)
text(fitReplicates$x[,c(3,4)], row.names(datasetrep), cex = 0.6)
plot(fitReplicates$x[,c(4,5)], lwd = 0, cex = 0)
text(fitReplicates$x[,c(4,5)], row.names(datasetrep), cex = 0.6)

summary(fitReplicates)

screeplot(fitReplicates, type = "lines")

dev.off()

###outliers with Pearson correlation and box plots####

medianReading <- as.data.frame(t(apply(peptideStatsArchsinh, 1, function(x) c(by(x, Hours, function(x) median(t(na.omit(x))))))))
corr2median = sapply(1:length(Hours), 
                     function(x){cor(peptideStatsArchsinh[,x], medianReading[,as.character(Hours[x])], 
                                     method = "pearson", 
                                     use = "pairwise.complete.obs")})
names(corr2median) <- names(peptideStatsArchsinh)

pdf("outlier_boxplot.pdf", width=30, height=10)
ratio2median = na.omit(data.frame(sapply(1:length(Hours), 
                                         function(x){peptideStatsArchsinh[,x]-medianReading[,as.character(Hours[x])]})))
names(ratio2median) <- names(peptideStatsArchsinh)
boxplot(ratio2median, xlab="sample", ylab="archsinh(normalised abundance)-sample median for zt", notch=TRUE, main="Ratio to the sample median")
mp <- barplot(corr2median,
              main="Pearsons correlation of protein abundance to median sample abundance with 1 standard deviation", 
              xlab="Sample", ylab="pearsons correlation",ylim=c(0.5,1), xpd = FALSE)
segments(mp, corr2median - sd(corr2median), mp, corr2median + sd(corr2median), lwd=2)

dev.off()
```
