## Supplementary Data S4 for "The circadian clock gene circuit controls protein and phosphoprotein rhythms in *Arabidopsis thaliana*": OutlierFigures.docx

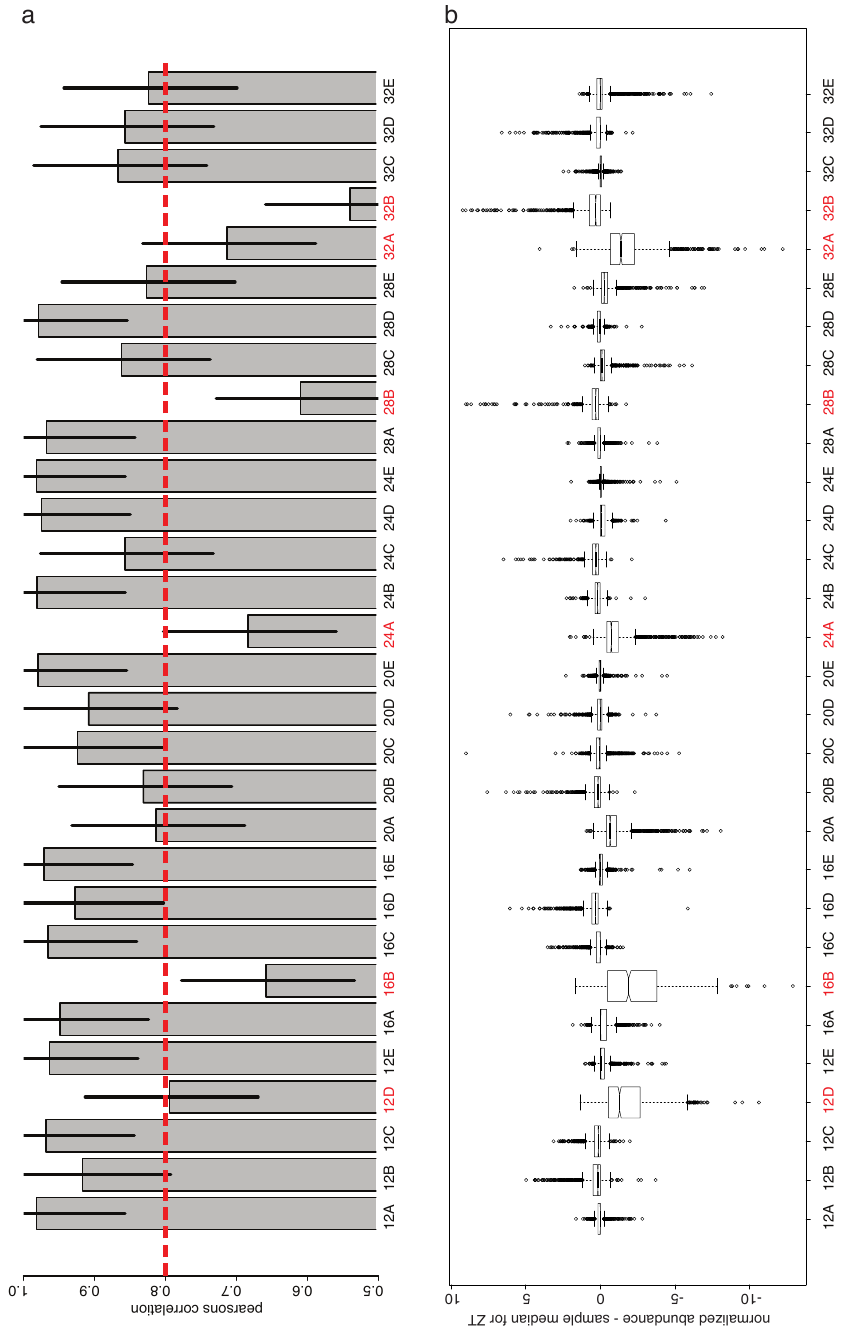

Correlation analysis of the merged WT phosphoproteomics dataset I (arcsinh transformed data). a) Pearsons correlation of protein abundance to median sample abundance, errorbars: standard deviation. The dashed red line indicates cutoff chosen for removal of outliers. Removed samples are indicated in red. b) Boxplot diagram of the differences between the replicate values and the time point median of each phosphopeptide.

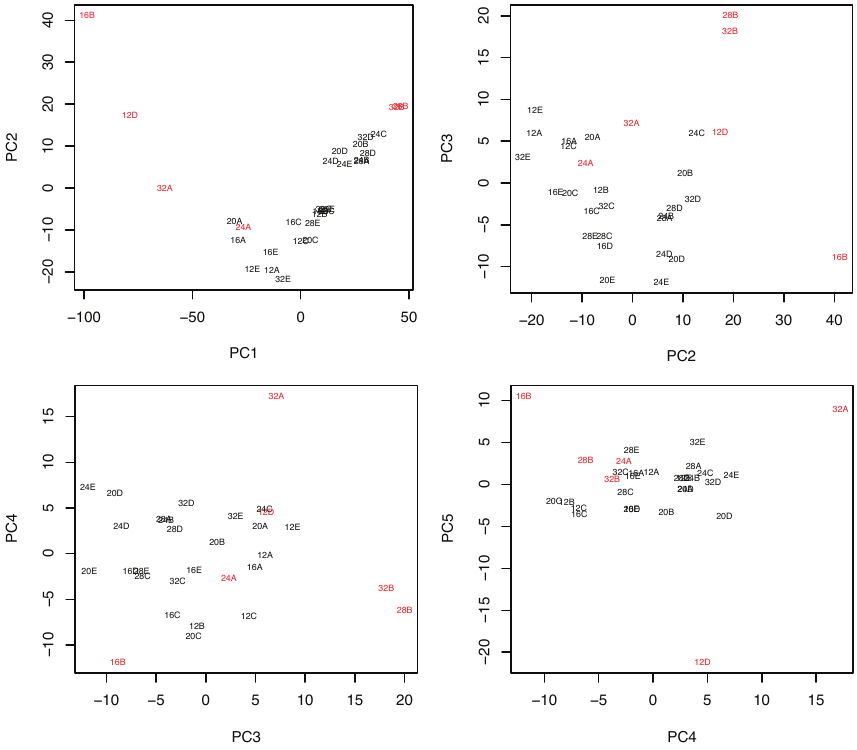

Figure S 2: PCA analysis of the merged WT phosphoproteomics I data. Outliers are indicated in red.

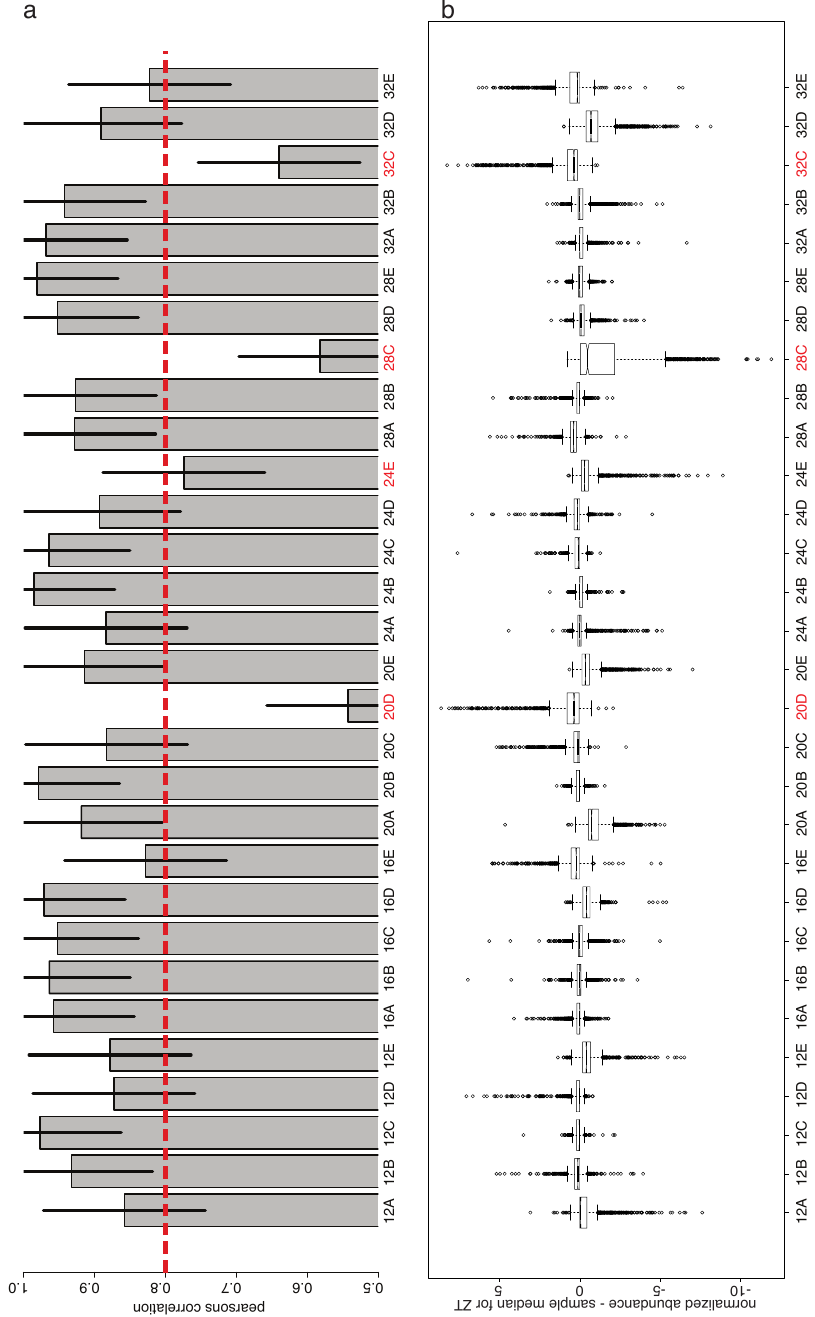

Correlation analysis of the merged CCA1-Ox phosphoproteomics dataset I (arcsinh transformed data). a) Pearsons correlation of protein abundance to median sample abundance, errorbars: standard deviation. The dashed red line indicates cutoff chosen for removal of outliers. Removed samples are indicated in red. b) Boxplot diagram of the differences between the replicate values and the time point median of each phosphopeptide.

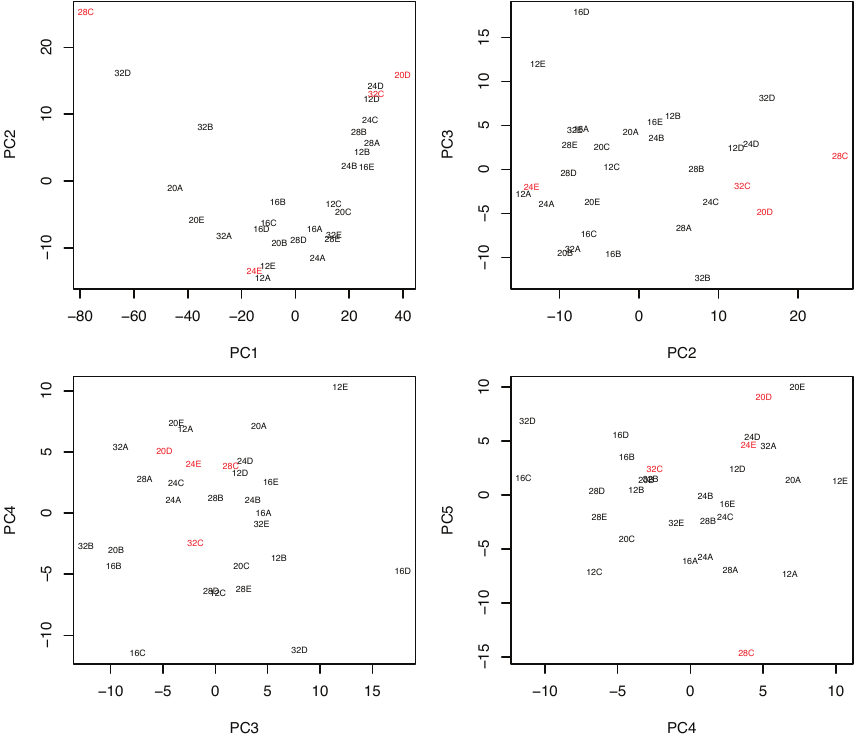

PCA analysis of the merged CCA1-Ox phosphoproteomics I data. Outliers are indicated in red.

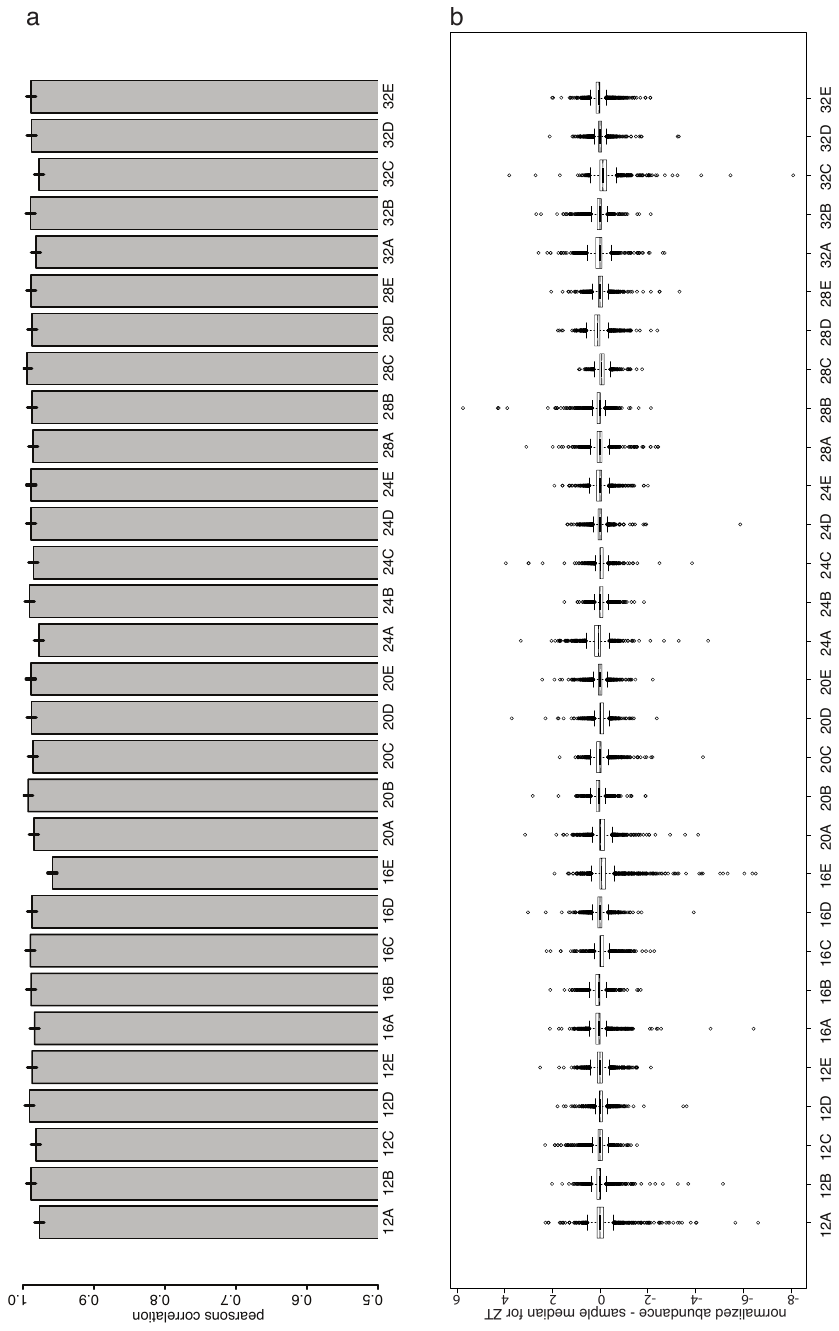

Correlation analysis of the merged WT global proteomics dataset I (arcsinh transformed data). a) Pearsons correlation of protein abundance to median sample abundance, errorbars: standard deviation. b) Boxplot diagram of the differences between the replicate values and the time point median of each phosphopeptide.

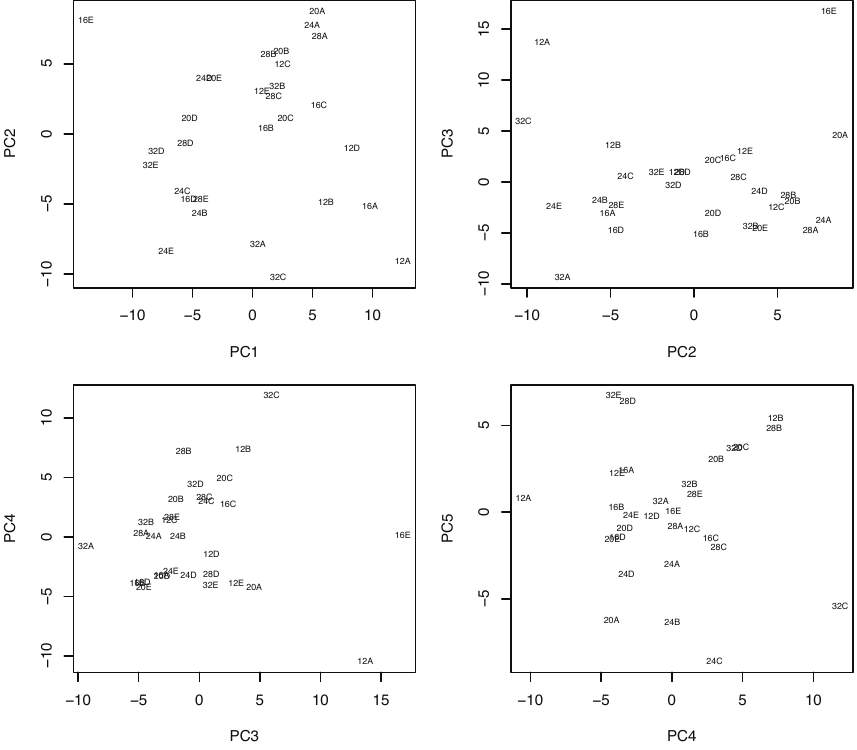

PCA analysis of the merged WT global proteomics I data.

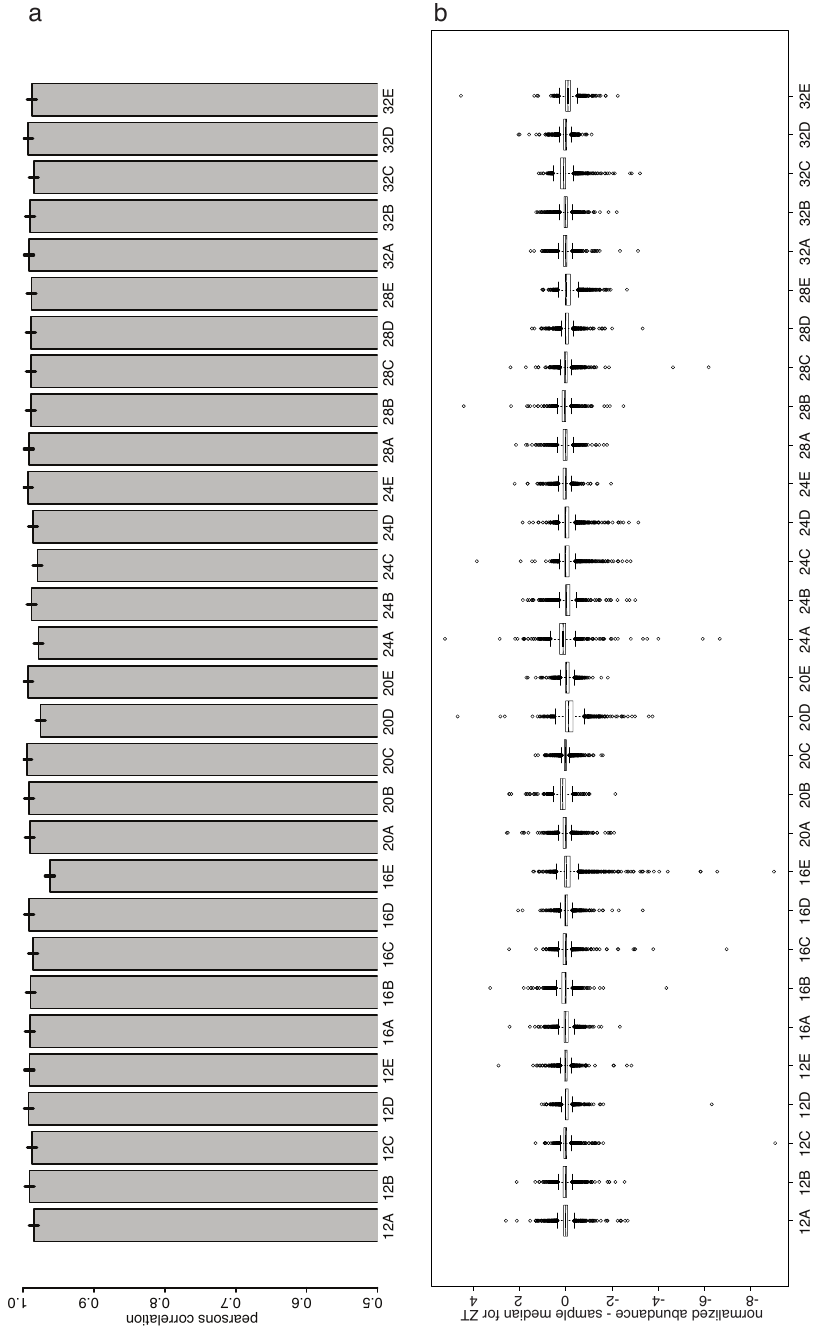

Correlation analysis of the merged CCA1-Ox global proteomics dataset I (arcsinh transformed data). a) Pearsons correlation of protein abundance to median sample abundance, errorbars: standard deviation. b) Boxplot diagram of the differences between the replicate values and the time point median of each phosphopeptide.

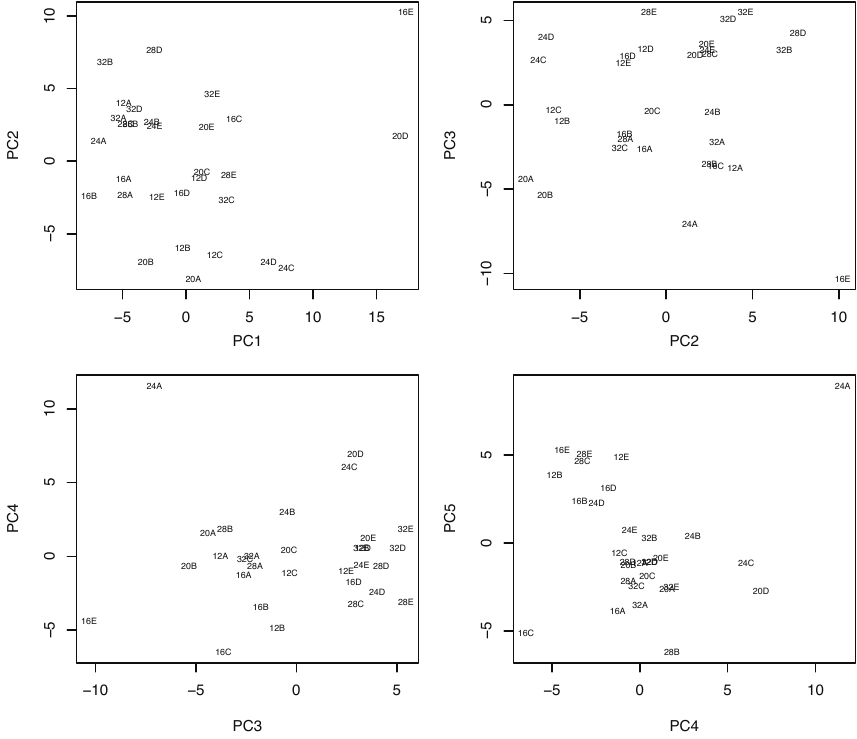

PCA analysis of the merged CCA1-Ox global proteomics I data. Outliers are indicated in red.

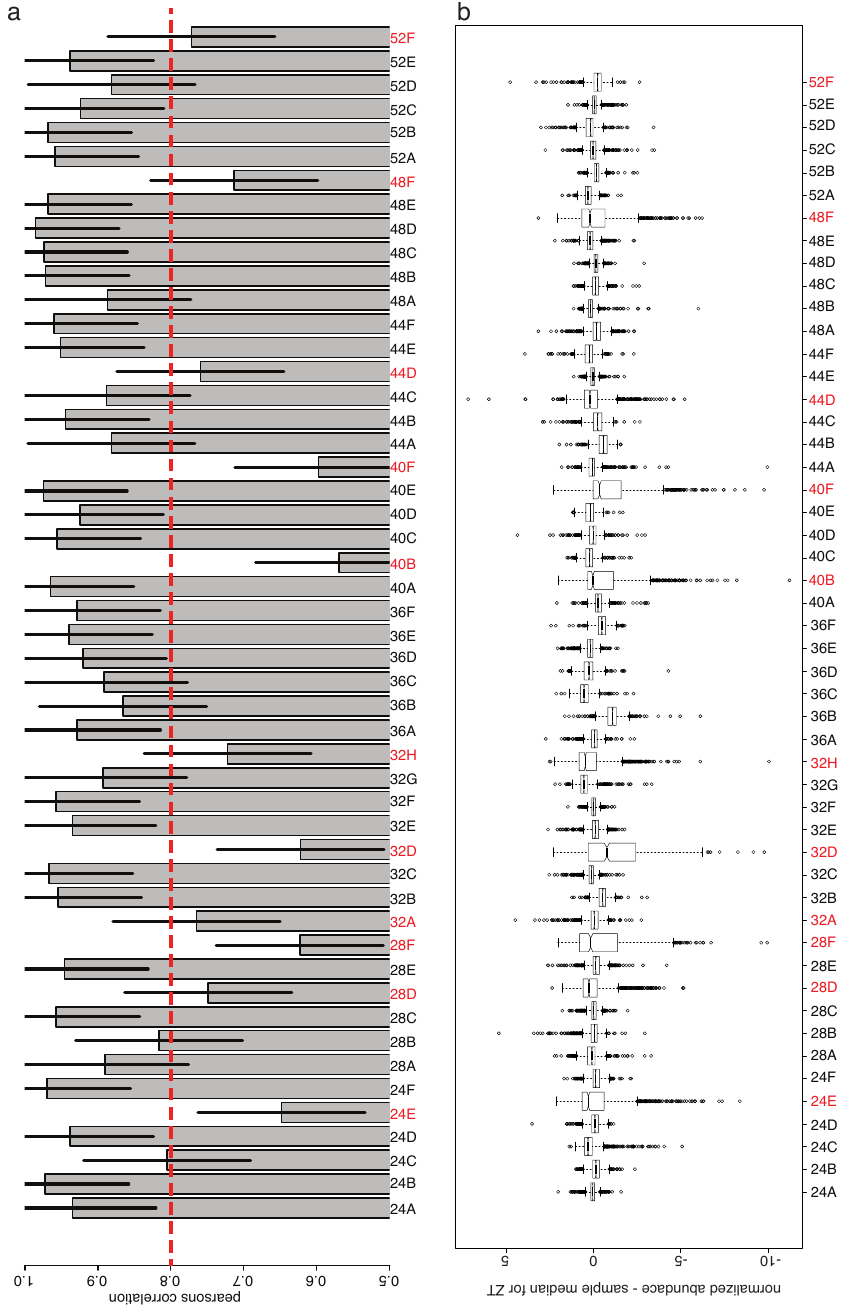

Correlation analysis of the merged WT phosphoproteomics dataset II (arcsinh transformed data). a) Pearsons correlation of protein abundance to median sample abundance, errorbars: standard deviation. The dashed red line indicates cutoff chosen for removal of outliers. Removed samples are indicated in red. b) Boxplot diagram of the differences between the replicate values and the time point median of each phosphopeptide.

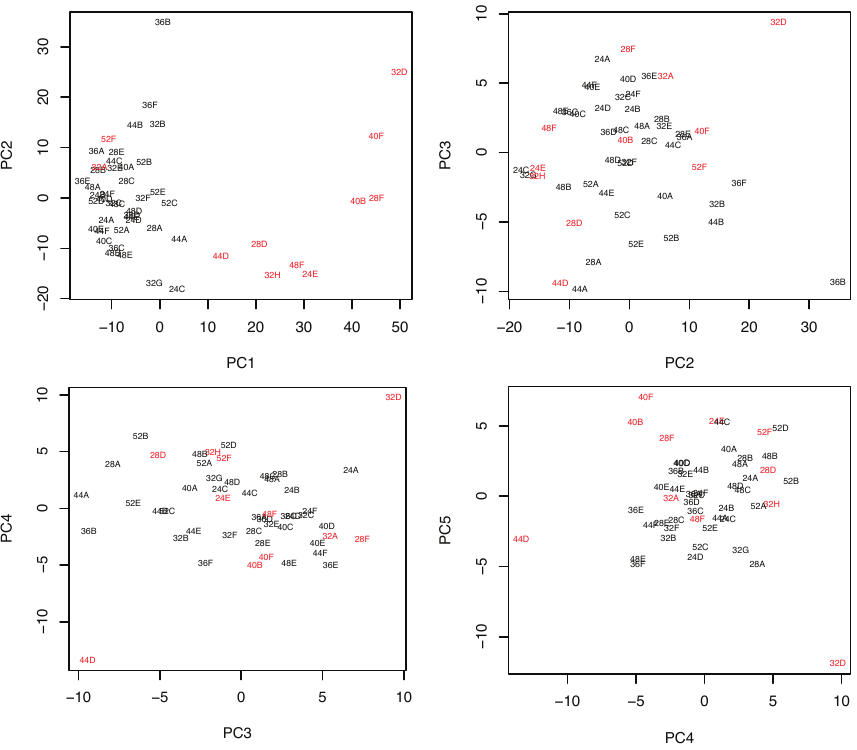

PCA analysis of the merged WT phosphoproteomics dataset II. Outliers are indicated in red.

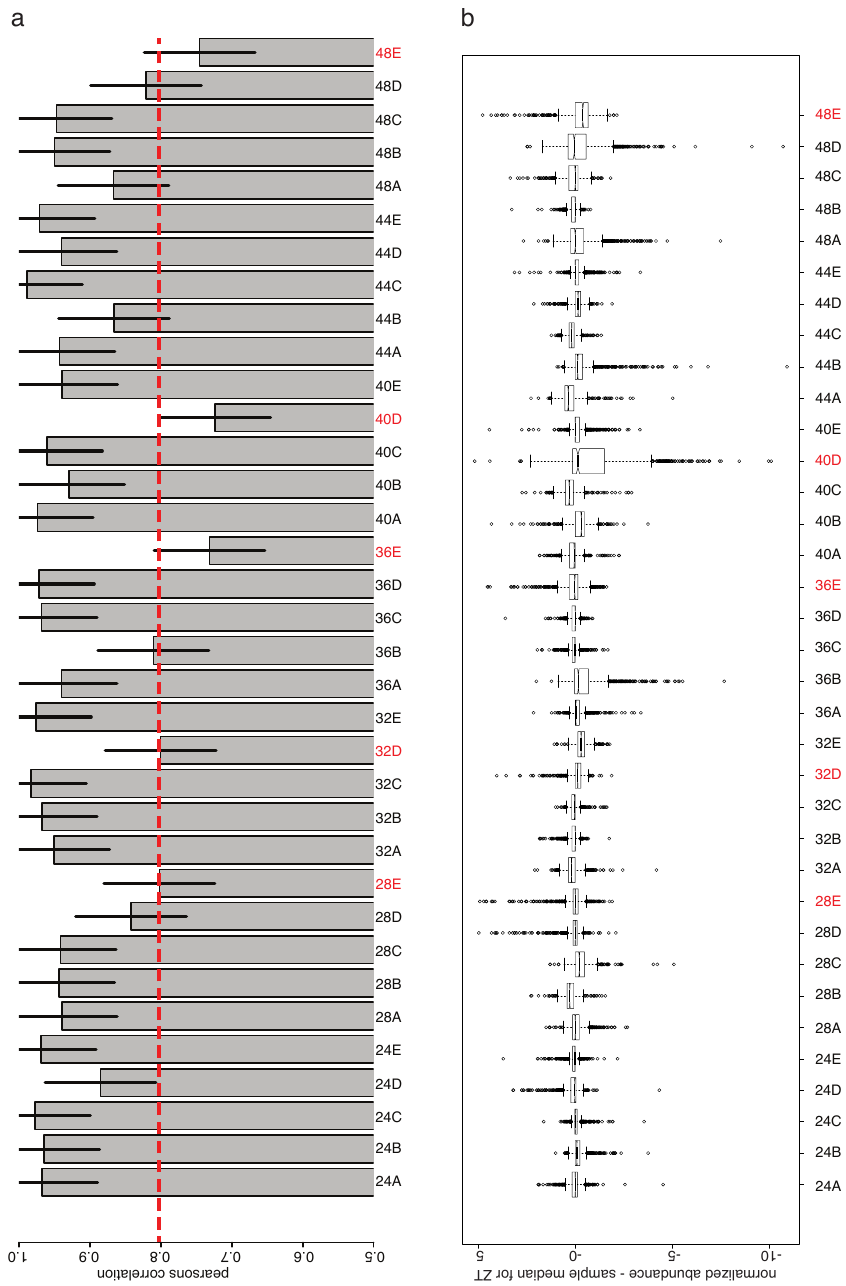

Correlation analysis of the merged CCA1-Ox phosphoproteomics dataset II (arcsinh transformed data). a) Pearsons correlation of protein abundance to median sample abundance, errorbars: standard deviation. The dashed red line indicates cutoff chosen for removal of outliers. Removed samples are indicated in red. b) Boxplot diagram of the differences between the replicate values and the time point median of each phosphopeptide.

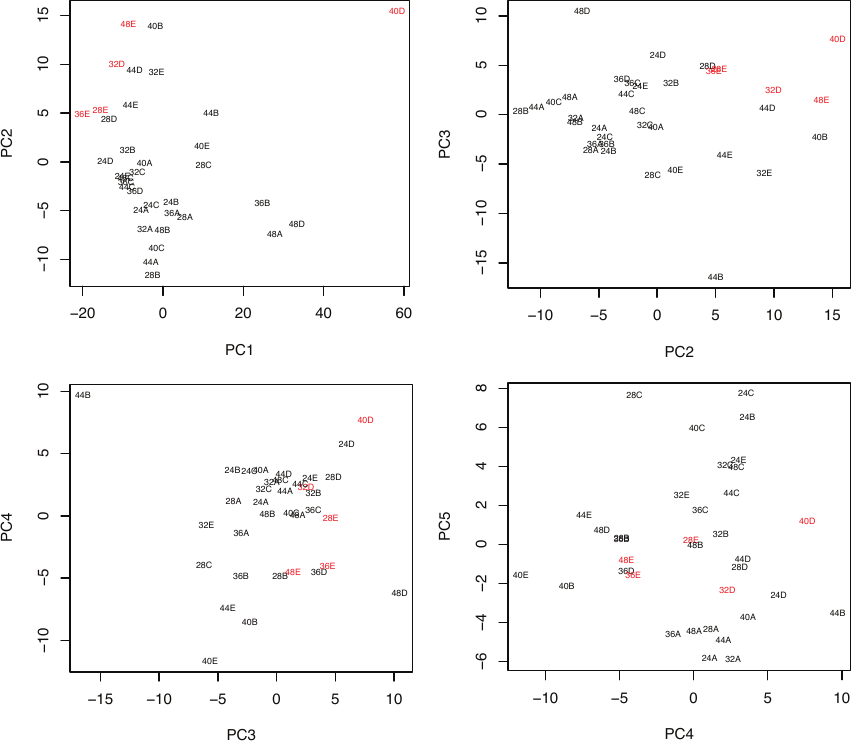

PCA analysis of the merged CCA1-Ox phosphoproteomics dataset II. Outliers are indicated in red.

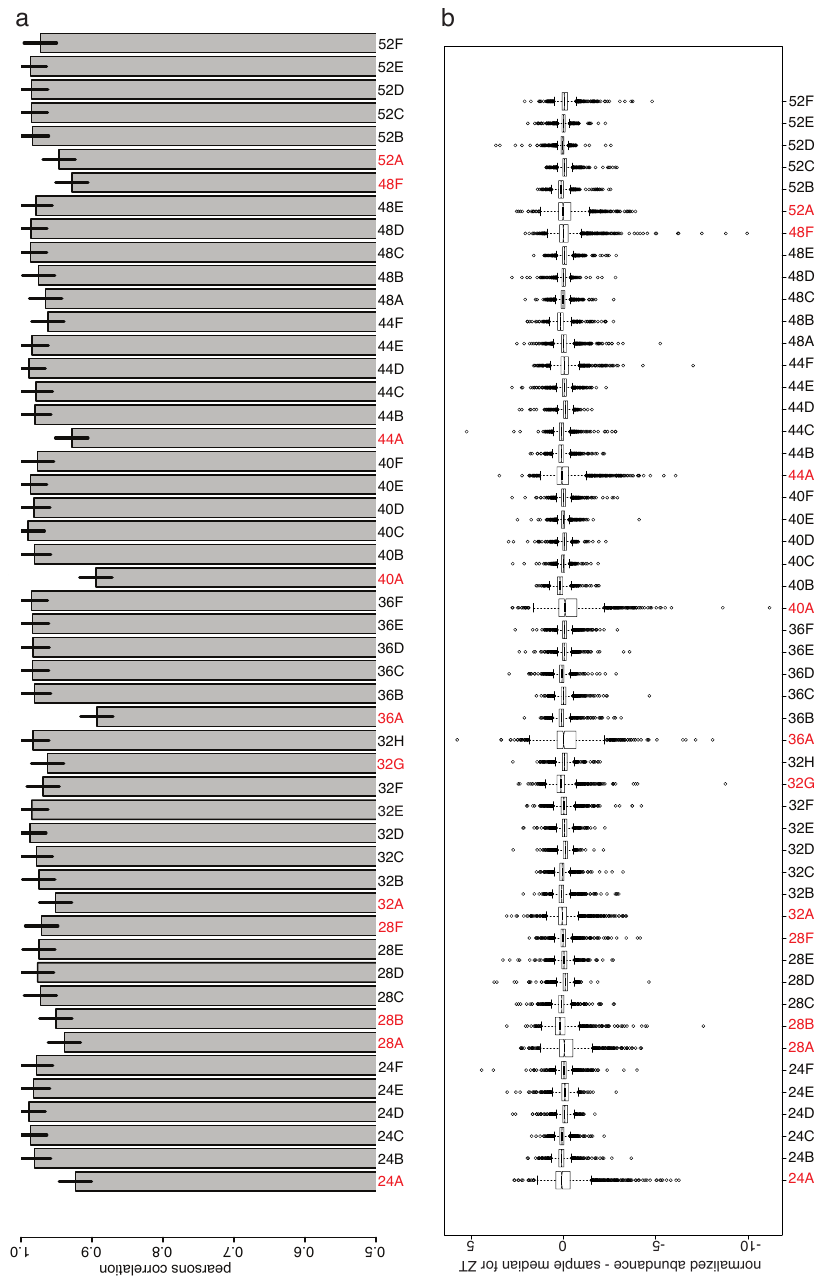

Correlation analysis of the merged WT global proteomics dataset II (arcsinh transformed data). a) Pearsons correlation of protein abundance to median sample abundance, errorbars: standard deviation. Removed samples are indicated in red. b) Boxplot diagram of the differences between the replicate values and the time point median of each phosphopeptide.

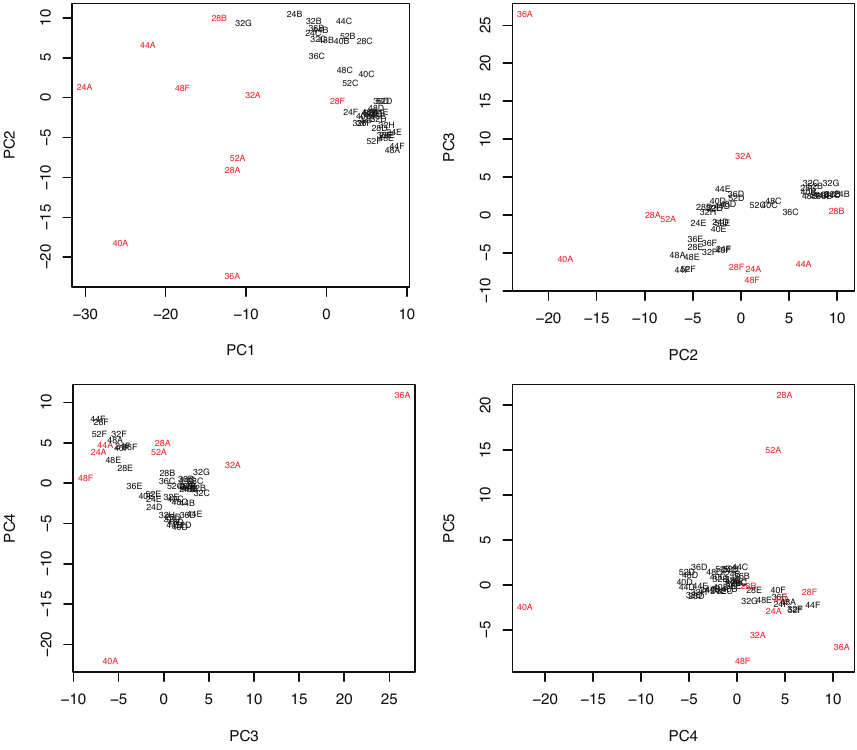

PCA analysis of the WT global proteomics dataset II. Outliers are indicated in red.

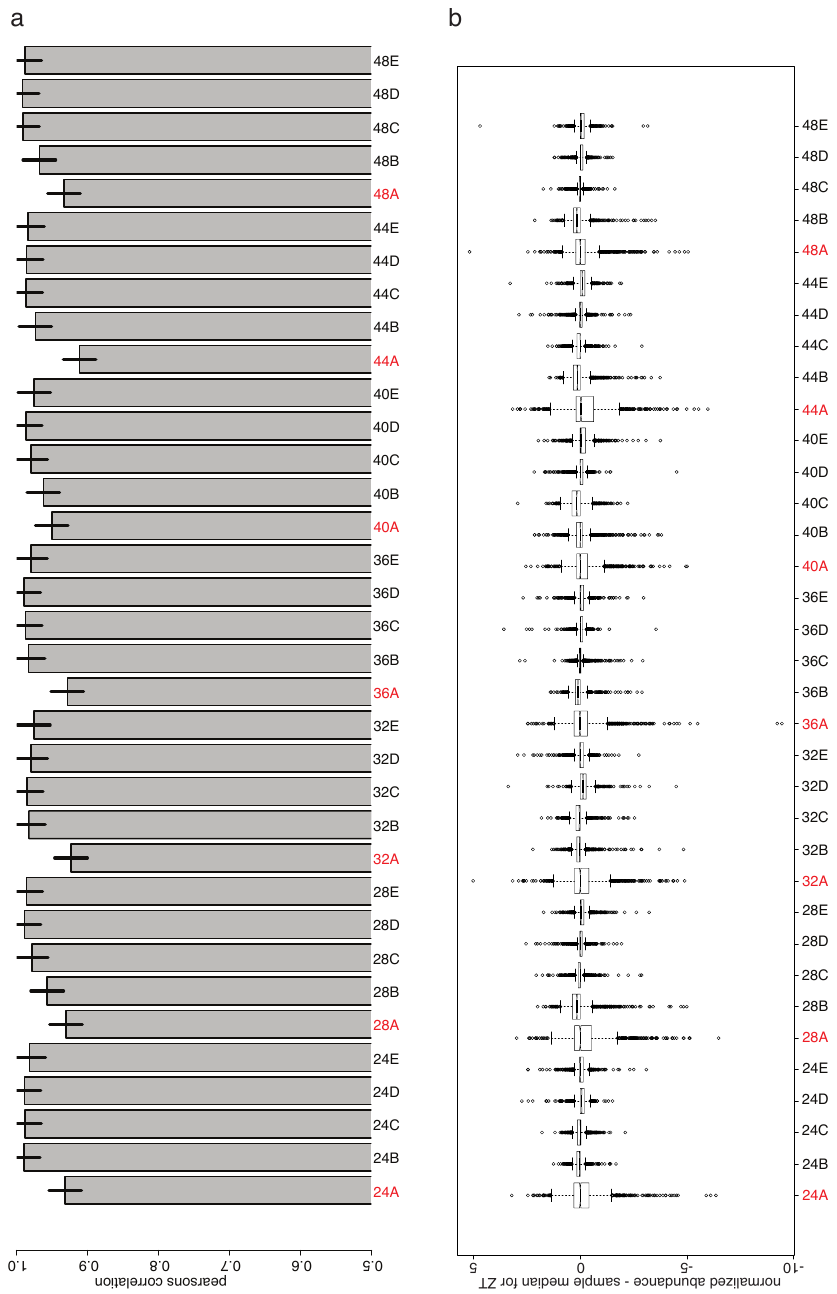

Correlation analysis of the merged CCA1-Ox global proteomics dataset II (arcsinh transformed data). a) Pearsons correlation of protein abundance to median sample abundance, errorbars: standard deviation. Removed samples are indicated in red. b) Boxplot diagram of the differences between the replicate values and the time point median of each phosphopeptide.

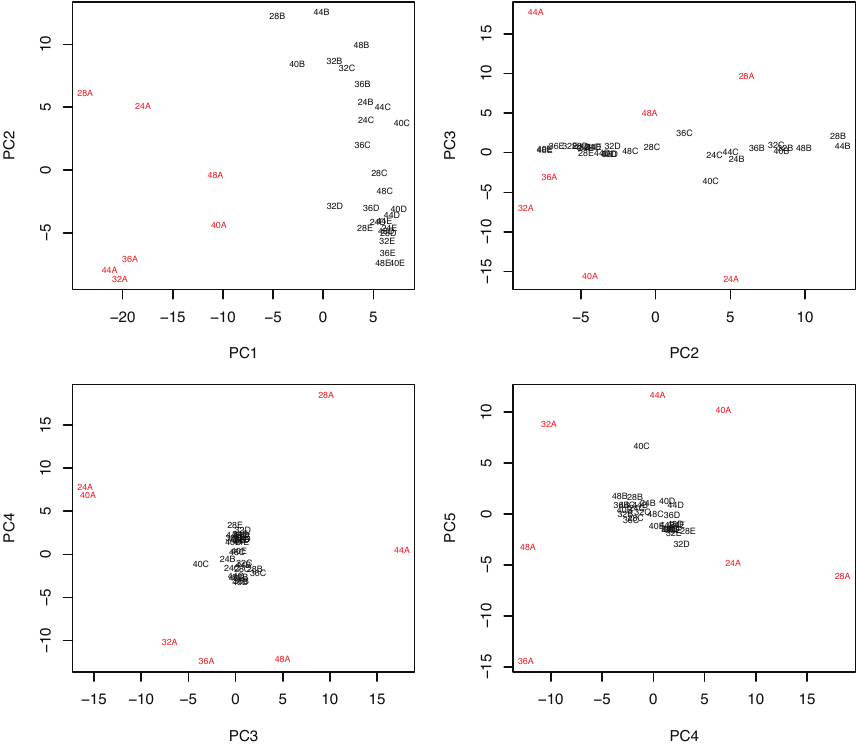

PCA analysis of the CCA1-Ox global proteomics dataset II. Outliers are indicated in red.
