## Supplementary figures and images for "The circadian clock gene circuit controls protein and phosphoprotein rhythms in *Arabidopsis thaliana*"

### outlier_boxplot.pdf

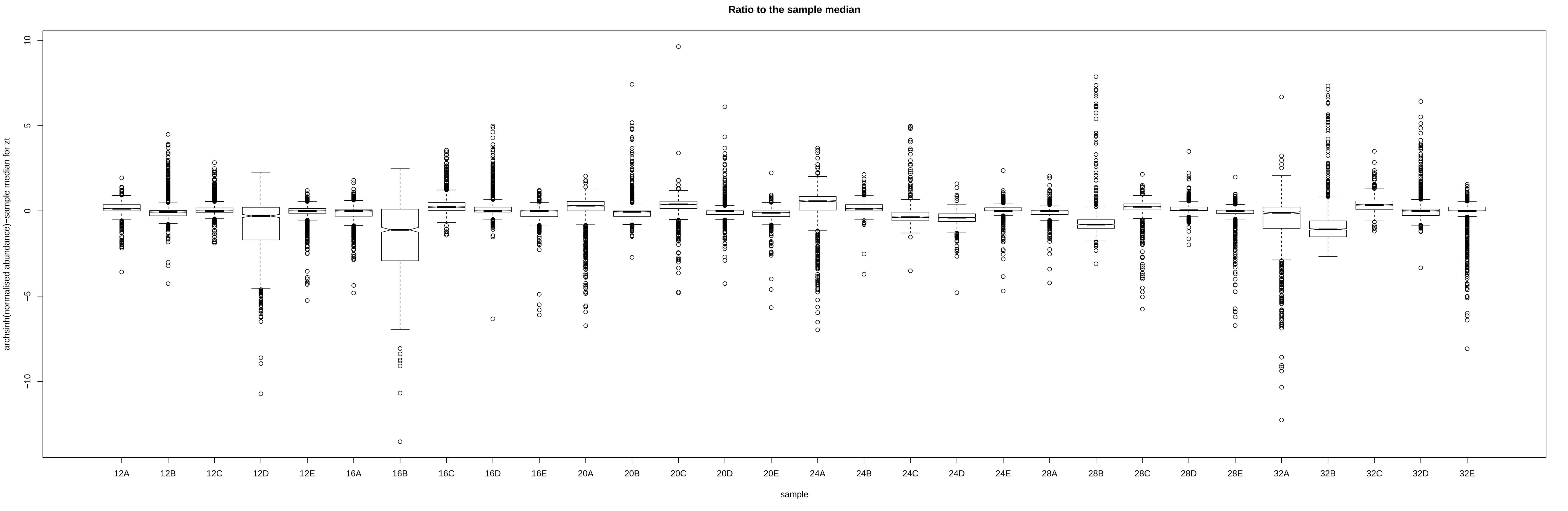

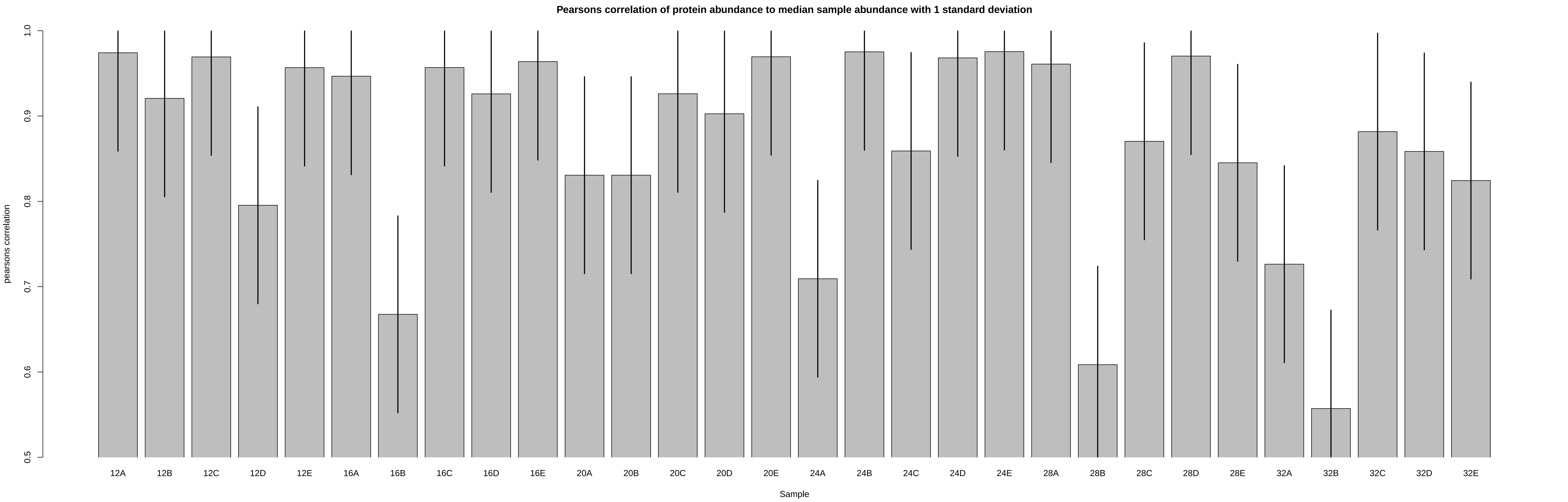

### outlier_PCA.pdf

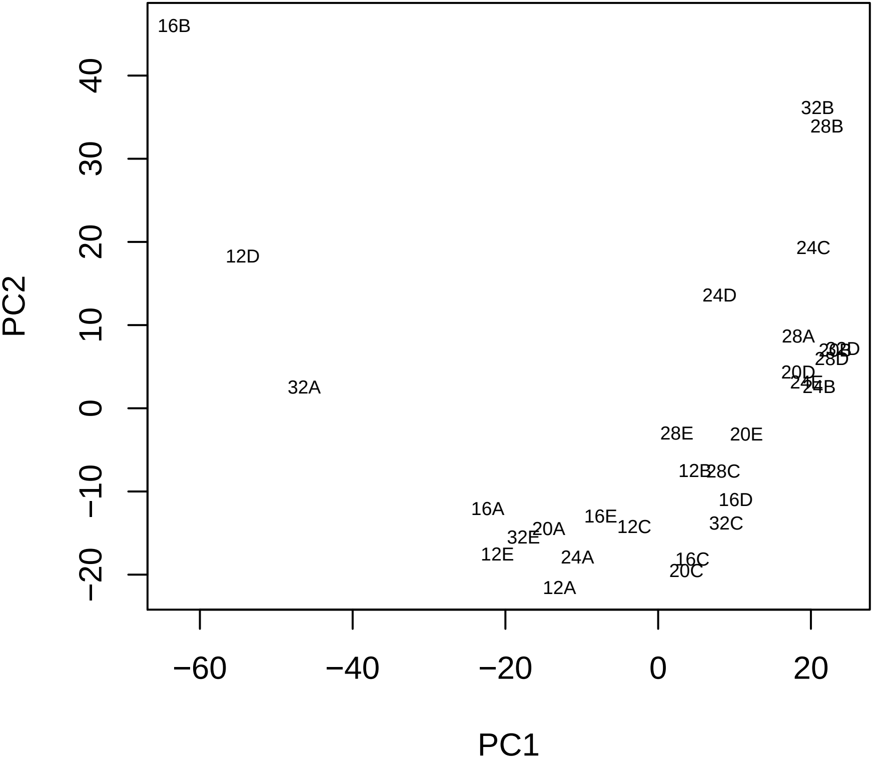

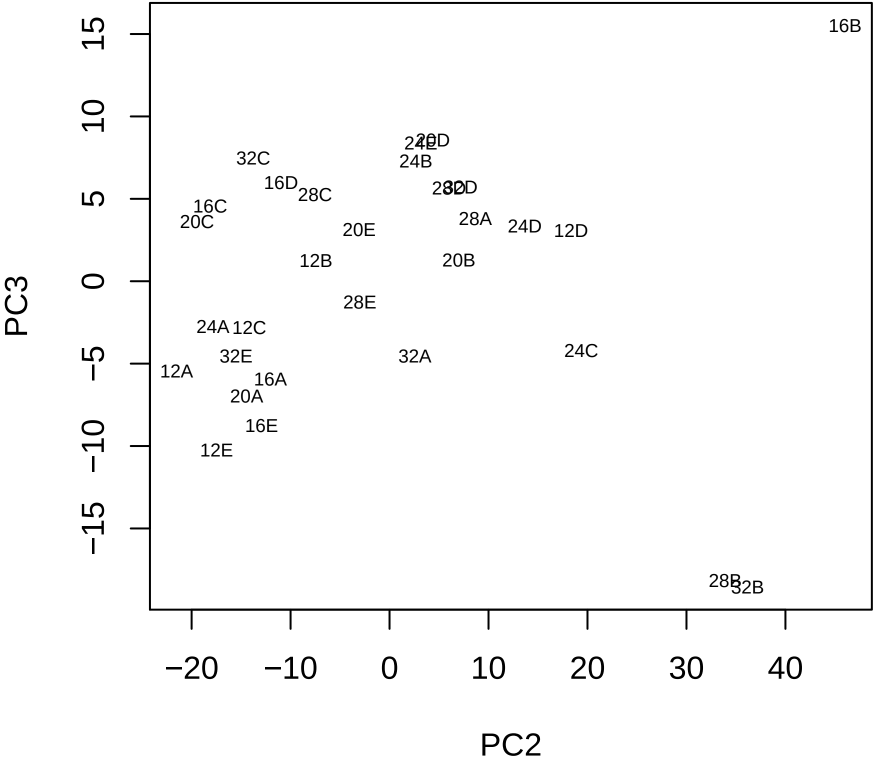

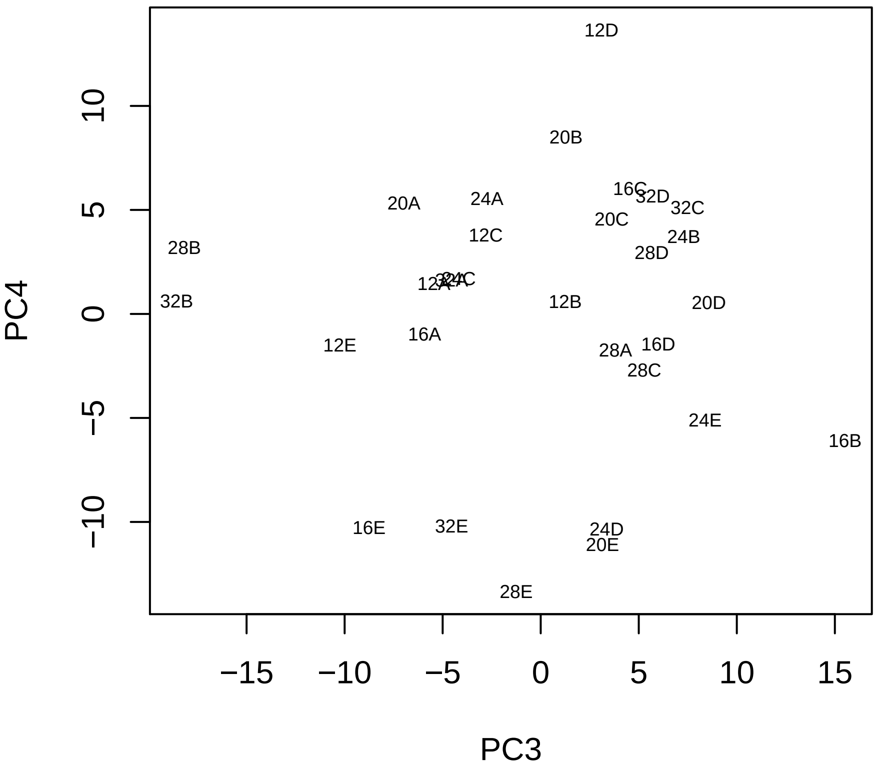

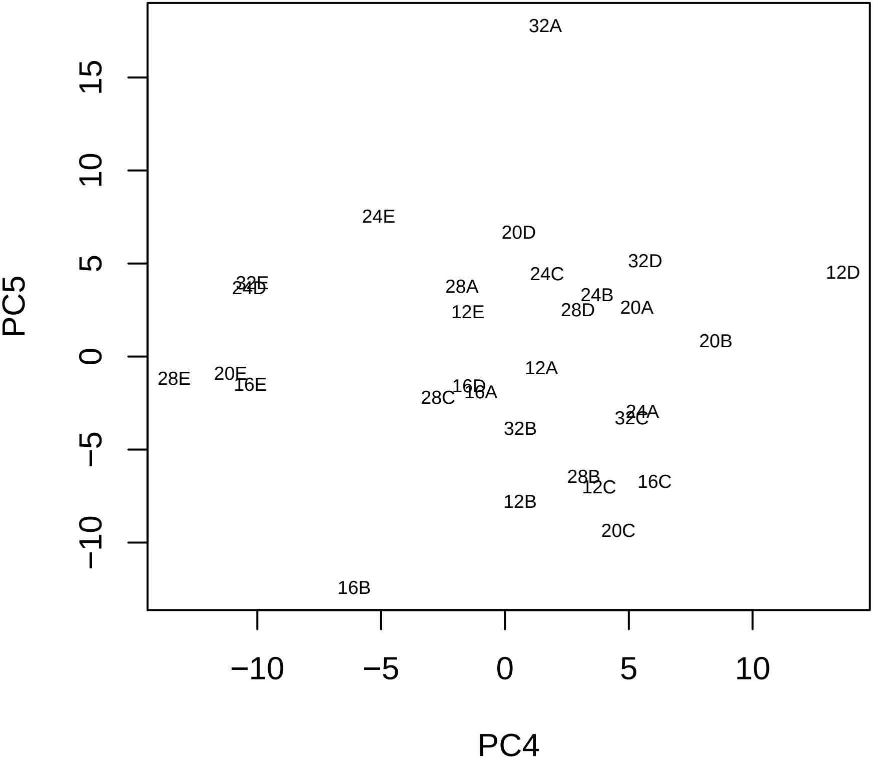

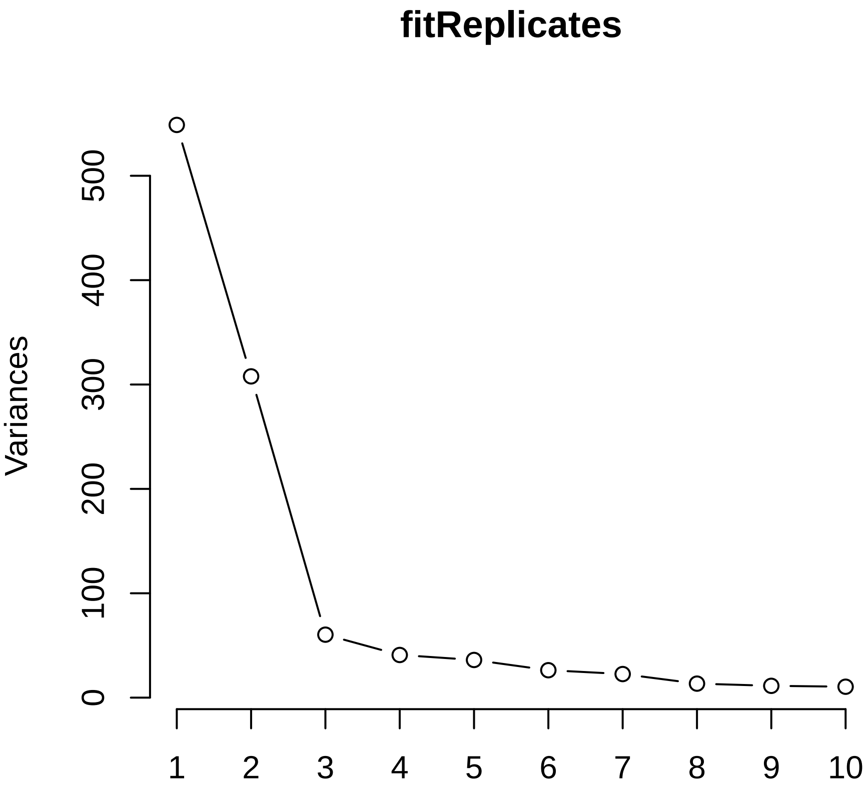
