## Supplementary Data S5 for "The circadian clock gene circuit controls protein and phosphoprotein rhythms in *Arabidopsis thaliana*": JTK_CYCLE.R

```
# Shewchuk algorithms for adaptive precision summation used in jtkdist
# http://www.cs.cmu.edu/afs/cs/project/quake/public/papers/robust-arithmetic.ps

fast.two.sum <- function(a,b) {                   # known abs(a) >= abs(b)
  x <- a+b
  bv <- x-a
  y <- b-bv
  if(y==0) return(x)
  c(x,y)
}

two.sum <- function(a,b) {                        # unknown order
  x <- a+b
  bv <- x-a
  av <- x-bv
  br <- b-bv
  ar <- a-av
  y <- ar+br
  if(y==0) return(x)
  c(x,y)
}

expansion.sum <- function(g) {
  g <- g[order(abs(g))]
  z <- fast.two.sum(g[2],g[1])
  q <- z[1]
  h <- NULL
  if(length(z)!=1) h <- z[2]
  n <- length(g)
  if(n==2) return(c(h,q))
  for(i in 3:n) {
    z <- two.sum(q,g[i])
    q <- z[1]
    if(length(z)!=1) h <- c(h,z[2])
  }
  c(h,q)                                          # strongly non-overlapping values
}

JTK.AMPFACTOR <- sqrt(2)                          # 1/median(cosine) used to calculate amplitudes
JTK.PIHAT <- round(pi,4)                          # replacement for pi to ensure unique cos values

# jtkdist: calculate the exact null distribution using the Harding algorithm
# http://www.jstor.org/pss/2347656

jtkdist <- function(timepoints,reps=1,normal=FALSE) {
  
  if(length(reps)==timepoints) {
    tim <- reps                                   # support for unbalanced replication
  } else {
    tim <- rep(reps[1],timepoints)                # balanced replication
  }
  JTK.GRP.SIZE <<- tim                            # sizes of each replicate group
  JTK.NUM.GRPS <<- length(tim)                    # timepoints = number of groups
  JTK.NUM.VALS <<- sum(tim)                       # number of data values (independent of period and lag)
  JTK.MAX <<- (sum(tim)^2-sum(tim^2))/2           # maximum possible jtk statistic
  JTK.DIMS <<- c(sum(tim)*(sum(tim)-1)/2,1)

  nn <- JTK.NUM.VALS
  ns <- JTK.GRP.SIZE
  maxnlp <- lfactorial(nn)-sum(lfactorial(ns))    # maximum possible negative log p-value
  limit <- log(.Machine$double.xmax)              # largest representable nlp
  normal <- normal | (maxnlp>limit-1)             # switch to normal approximation if maxnlp is too large
  if(normal) {
    JTK.VAR <<- (nn^2*(2*nn+3) - 
                  sum(ns^2*(2*ns+3)))/72          # variance of jtk
    JTK.SDV <<- sqrt(JTK.VAR)                     # standard deviation of jtk
    JTK.EXV <<- JTK.MAX/2                         # expected value of jtk
    JTK.EXACT <<- FALSE
    return(0)                                     # omit calculation of exact distribution
  }
  MM <- floor(JTK.MAX/2)                          # mode of the jtk distribution
  cf <- as.list(rep(1,MM+1))                      # initial lower half cumulative frequency distribution
    
  size <- JTK.GRP.SIZE                            # sizes of each group
  size <- size[order(size)]                       # ascending order for fastest calculation
  k <- JTK.NUM.GRPS                               # number of groups  
  
  N <- size[k]                  
  if(k>2) for(i in (k-1):2) {
    N <- c(size[i]+N[1],N)
  }
  for(i in 1:(k-1)) {                             # count permutations using the Harding algorithm
    m <- size[i]
    n <- N[i]
    
    if(n < MM) {
      P <- min(m+n,MM)
      for(t in (n+1):P) {                         # zero-based offset t
        for(u in 1+MM:t) {                        # one-based descending index u
          cf[[u]] <- expansion.sum(               # Shewchuck algorithm
            c(cf[[u]],-cf[[u-t]]))
        }
      }
    }
    Q <- min(m,MM)
    for(s in 1:Q) {                               # zero-based offset s
      for(u in 1+s:MM) {                          # one-based ascending index u
        cf[[u]] <- expansion.sum(                 # Shewchuck algorithm
          c(cf[[u]],cf[[u-s]]))
      }
    }
  }
  cf <- sapply(cf,sum)
  
  # cf now contains the lower-half cumulative frequency distribution;
  # append the symmetric upper-half cumulative distribution to cf
  
  if(JTK.MAX %% 2) {
    cf <- c(cf,2*cf[MM+1]-c(cf[MM:1],0))          # if JTK.MAX odd (mode is duplicated)
  } else {
    cf <- c(cf,cf[MM+1]+cf[MM]-c(cf[MM:2-1],0))   # if JTK.MAX even (unique mode is in lower half)
  }
  jtkcf <- rev(cf)                                # upper-tail cumulative frequencies for all integer jtk
  ajtkcf <- (jtkcf[-length(cf)]+jtkcf[-1])/2      # interpolated cumulative frequency values for all half-integer jtk  
  
  id <- 1+0:(2*JTK.MAX)                           # one-based indices for all jtk values
  cf <- id                                        # container for the jtk frequency distribution
  cf[!!id%%2] <- jtkcf                            # odd indices for integer jtk
  cf[!id%%2] <- ajtkcf                            # even indices for half-integer jtk
  cp <- cf/jtkcf[1]                               # all upper-tail p-values
  
  JTK.CP <<- cp                                   # same for all periods and lags
  JTK.EXACT <<- TRUE
}

# jtk.init: initialize the JTK environment for all periods
jtk.init <- function(periods, interval=1) {
      
  JTK.INTERVAL <<- interval
  JTK.PERIODS <<- periods
  JTK.PERFACTOR <<- rep(1:length(periods),ti=periods)
  
  tim <- JTK.GRP.SIZE
  timepoints <- JTK.NUM.GRPS
  timerange <- 1:timepoints-1                       # zero-based time indices
  JTK.CGOOSV <<- list()      
  JTK.SIGNCOS <<- list()

  for(i in 1:length(periods)) {
    period <- periods[i]
    time2angle <- 2*JTK.PIHAT/period                # convert time to angle using an approximate pi value
    theta <- timerange*time2angle                   # zero-based angular values across time indices
    cos.v <- cos(theta)                             # unique cosine values at each timepoint
    cos.r <- rank(cos.v)                            # ranks of unique cosine values
    cos.r <- rep(cos.r,ti=tim)                      # replicated ranks
  
    cgoos <- sign(outer(cos.r,cos.r,"-"))
    cgoos <- cgoos[lower.tri(cgoos)]
    cgoosv <- array(cgoos,dim=JTK.DIMS)
    JTK.CGOOSV[[i]] <<- matrix(
      ncol=period,nrow=nrow(cgoosv)
    )
    JTK.CGOOSV[[i]][,1] <<- cgoosv
  
    range <- 1:period
    cos.s <- sign(cos.v)[range]                     # signs over initial full cycle                
    cos.s <- rep(cos.s,ti=tim[range])
    JTK.SIGNCOS[[i]] <<- matrix(
      ncol=period,nrow=length(cos.s)
    )
    JTK.SIGNCOS[[i]][,1] <<- cos.s
    
    for(j in 2:period) {                            # one-based half-integer lag index j
      delta.theta <- (j-1)*time2angle/2             # angles of half-integer lags
      cos.v <- cos(theta+delta.theta)               # cycle left
      cos.r <- rank(cos.v)                          # ranks of unique phase-shifted cosine values
      cos.r <- rep(cos.r,ti=tim)                    # phase-shifted replicated ranks
    
      cgoos <- sign(outer(cos.r,cos.r,"-"))
      cgoos <- cgoos[lower.tri(cgoos)]
      cgoosv <- array(cgoos,dim=JTK.DIMS)
      JTK.CGOOSV[[i]][,j] <<- cgoosv
    
      cos.s <- sign(cos.v)[range]
      cos.s <- rep(cos.s,ti=tim[range])
      JTK.SIGNCOS[[i]][,j] <<- cos.s    
    }
  }
}

# jtkstat: calculate the p-values for all (period,phase) combos
jtkstat <- function(z) {
  
  stopifnot(all(is.finite(z)))                    # missing values are not supported
  foosv <- sign(outer(z,z,"-"))
  foosv <- foosv[lower.tri(foosv)]
  dim(foosv) <- JTK.DIMS
  
  JTK.CJTK <<- list()
  for(i in 1:length(JTK.PERIODS)) {
    JTK.CJTK[[i]] <<- apply(JTK.CGOOSV[[i]],2,
      function(cgoosv) {
        S <- sum(foosv*cgoosv)                    # Kendall's S score for this lag
        if(!S) return(c(1,0))                     # discreteness-corrected two-tailed p-value and S
        M <- JTK.MAX                              # maximum possible JTK statistic for this lag
        jtk <- (abs(S)+M)/2                       # two-tailed JTK statistic for this lag
        if(JTK.EXACT) {
          jtki <- 1+2*jtk                         # index into the exact upper-tail distribution
          p <- 2*JTK.CP[jtki]                     # exact two-tailed p-value for this lag
        } else {
          p <- 2*pnorm(-(jtk-1/2),
                -JTK.EXV,JTK.SDV)                 # two-tailed normal approximation with continuity correction
        }
        c(p,S)                                    # need S for phase and tau 
    })
  }
}

# jtkx: integration of jtkstat and jtkdist for repeated use
jtkx <- function(z) {
  
  jtkstat(z)                                      # calculate p and S for all (period,phase) combos
  
  pvals <- sapply(JTK.CJTK,function(cjtk) {
    cjtk[1,]
  })                                              # exact two-tailed p-values for all (period,phase) combos
  padj <- p.adjust(unlist(pvals),"bonf")          # Bonferroni adjusted two-tailed p-values
  JTK.ADJP <<- min(padj)                          # global minimum adjusted p-value
  
  padj <- split(padj,JTK.PERFACTOR)
  minpadj <- sapply(padj,min)                     # minimum adjusted p-value for each period
  
  peris <- which(JTK.ADJP==minpadj)     ##        # indices of all optimal periods
  pers <- JTK.PERIODS[peris]                      # all optimal periods
  
  lagis <- lapply(padj[peris],function(z) {
    which(JTK.ADJP==z)									##				# indices of all optimal lags
  })                                              # list of optimal lag indices for each optimal period
  count <- sum(sapply(lagis,length))              # total number of optimal lags for all optimal period
  
  sumper <- 0
  sumlag <- 0
  sumamp <- 0
  
  for(i in 1:length(pers)) {
    per <- pers[i]
    peri <- peris[i]
    cjtk <- JTK.CJTK[[peri]]
    sc <- JTK.SIGNCOS[[peri]]
		w <- z[1:nrow(sc)]									##
		w <- (w-median(w))*JTK.AMPFACTOR		##
    
    for(lagi in lagis[[i]]) {  
      sumper <- sumper+per
      
      S <- cjtk[2,lagi]                           # optimal Kendall's S
      s <- sign(S)
      if(!s) s <- 1
  
      lag <- (per +(1-s)*per/4 -(lagi-1)/2)%%per
      sumlag <- sumlag+lag
  
      signcos <- sc[,lagi]
      amp <- s*w*signcos									##
			amp <- median(amp[!!amp])
      sumamp <- sumamp+amp
    }
  }
  JTK.PERIOD <<- JTK.INTERVAL*sumper/count        # mean optimal period (hours)
  JTK.LAG <<- JTK.INTERVAL*sumlag/count           # mean lag to peaks of optimal cosine waves (hours)
  JTK.AMP <<- max(0,sumamp)/count                 # mean amplitude of optimal cosine waves
  JTK.TAU <<- abs(S)/JTK.MAX                      # all optimal abs(S) are the same
}
```
