## Supplementary Data S5 for "The circadian clock gene circuit controls protein and phosphoprotein rhythms in *Arabidopsis thaliana*": run_JTK.R

```
# Script written by Johanna Krahmer, to analyse large sample timecourse data for 1 cycle
#   or longer for rhythmicity and period using JTK_CYCLE but
#   with variable replicate numbers for each rhythm to be tested.

# What this script does:
# - reads in file with circadian phosphoproteomics data (easily adaptable for global proteomics)
# - substitutes all 0 values of measurements for NA
# - ignores IDs (rows) where one or more time points do not have any measurements, will results in NA,
#   i.e. if a time point has 5 replicates and all 5 replicates are NA, the whole ID is not considered for JTK_CYCLE analysis
# - runs JTK_CYCLE script on each ID. Adjusts number of replicates if there are NA values
#   since it runs JTK_CYCLE for each row rather than entire file.
# - correction for multiple testing: BH option
# - determines some additional values that will be useful: minimum, maximum, time of peaks and troughs
# - gives counts of significant (with and without adjustment for multiple testing) IDs for each timepoint
# - note: considers periods 22 to 26h. If a different range is required, change line "periods <- 5.5:6.5"
#   to different sampling frequency
# - assumes sampling every 4h. Change line "jtk.init(periods,4)" (the '4' means sampling every 4h)
# - for traces where averages go either just up or just down over the entire timecourse, the value 1 is added to the pvalue as well as the qvalue

# output of JTK_CYCLE: (IDs are in same order as input file apart but note that some lines have been removed)
#   ADJ.P: anadjusted p values
#   BH.Q: BH corrected p values
#   PER: predicted period
#   LAG: some phase information (haven't used this, don't know what exactly it does)
#   AMP: some amplitude information (haven't used this, don't know what exactly it does)

# How to use this script
# Prepare input file: either from after qpMerge (phosphopeptides) or Progenesis output (global proteins, but need to adjust script slightly for global)
#   needed: peptide ID, "Sequence", "Accession", "Description", "Progenesis column" (modifications) normalized abundance values (if using normalized abundance)
#   other columns do not bother the script (unless same column headers on normalized and raw data)
#   Headers of data columns in the input file need specific structure:
#      [some experiment name that is the same for each sample]_[time point in hours]_[replicate number]
#      example: X3702_PhII_wt_24_2, where time point is 24h, replicate 2
#  Save input file as comma delimited csv file and quote all text

# Prepare metadata file using metdata example / template file and save:
# (1) write hour numbers of time point in column "time point"
# (2) write number of replicates for each time point in column "number of replicates"
# (3) write column header of phosphopeptide or protein accessions in data input file into column
#    "column name of peptide (phospho) or accession (global)"; is e.g. X. for peptides, accession for global protein data
# (4) In cell under "Data column name before timepoint", write the non-changing part of the sample names (e.g. X3702_PhII_wt), without undercore at the end
# (5) In column "replicates", write the number of replicates for each time point written in column "at timepoints" (see example template)
# note: all cells that are in bold in the example template need to be updated for each input file

# What to change in this script: (all at the very start of this script; double backward slashes on PC???)
#   name project (2nd line of code)
#   paths and file names
#   for input of metadata and data files
#   for saving output data file, an .xlsx file (to create object JTK_output_filename)
#   for saving a temporary annotation file, a .txt file (annot_filename)
#   for saving a temporary data file, a .txt file (data_filename)
#   for finding the JTK_CYCLE.R resource


# set path!

library(WriteXLS)
project <- "PhI_wt" # change to name of project

####### put in all file paths:
# metadata file:
metadata=read.table("metadata.csv", sep=",", quote="\"", header=1)
# data file:
PData = read.table('input_data.csv', skip=0, header=1, sep=",", quote="\"")
# final output file:
JTK_output_filename="JTK_output.xlsx"
# temporary file for annotations:
annot_filename="annot_temp.txt"
# temporary file for data:
data_filename="data_temp.txt"

# JTK_CYCLE source file:
source("JTK_CYCLE.R")

# get parameters from metadata:
timepoints=metadata$"time.point"[is.na(metadata$"time.point")==FALSE]
number_of_replicates = metadata$"number.of.replicates"

# make the Hours factor from metadata
factor=c()
for (i in 1:length(timepoints)){
  timepoint_factor=matrix(timepoints[i],nrow=1,ncol=number_of_replicates[i])
  factor=c(factor, timepoint_factor)
}

factor=as.factor(factor)

Hours = factor
constname=toString(metadata$"Data.column.name.before.timepoint"[1])


replicate_numbers=metadata$"replicates"
replicates_factor=replicate_numbers

sample_names=matrix(0, nrow=1, ncol=length(factor))
for (i in 1:length(sample_names)){
  sample_names[i] = paste0(constname,'_',factor[i],'_',replicates_factor[i])
}

# find the columns that have these sample names, create data frame measurements with all the relevant measurements

samplecolumn = matrix(NA, nrow=1, ncol=length(sample_names))
PData_colnames = names(PData)
for (i in 1:length(sample_names)){
  samplecolumn[i]=which(PData_colnames == as.character(sample_names[i]))
}
# exchange all zero values for NA
PData[PData==0]=NA
measurements = PData[,min(samplecolumn):max(samplecolumn)]

Hours <- factor

measurements_rownames = metadata$"column.name.of.peptide..phospho..or.accession..global."[1]
row.names(measurements) <- PData[,toString(measurements_rownames)]

dump_row=rep(0,nrow(measurements))

for (i in 1:nrow(measurements)) {
  NAPositions=rep(0, length(sample_names))
  ithrow=measurements[i,][1:length(sample_names)]
  for (j in 1:length(measurements)){
    if (is.na(ithrow[[j]]) == TRUE){
      NAPositions[j]=1
    }
  }
  
  # from NAPositions vector get number of NA values for each time point:
  NAPositions_startstop=matrix(NA, nrow=length(timepoints), ncol=2)
  names(NAPositions_startstop)=c("start timepoint","end timepoint")
  NA_counts=matrix(NA, nrow=length(timepoints), ncol=1)
  endcounter=0
  NAtimepointcounter=0
  for (j in 1:length(timepoints)){
    startcounter=endcounter+1
    endcounter=endcounter+number_of_replicates[j]
    NAPositions_startstop[j,1]=startcounter
    NAPositions_startstop[j,2]=endcounter
    NA_counts=sum(NAPositions[startcounter:endcounter])
    if (NA_counts == number_of_replicates[j]){
      dump_row[i]=1      
    }
  }
}
  
measurements$dump_row=dump_row

# Now, do a JTK_CYCLE analysis for each row (i).
# Firstly, make data and annot matrices, but need to make sure it is in the right format to be read
# Saves a file for each data and annot object which is very unelegant but couldn't find any other
#    way that works
# Then generate the group.sizes vector depending on NAs.
# Then produce st object
# write data from st object into another dataframe called 'allresults'

# initialize some data frames:
allresults=data.frame(matrix(data=0, nrow=nrow(measurements),ncol=6))
norm_mean=data.frame(matrix(data=NA, nrow=nrow(measurements),ncol=length(timepoints)))
names_norm_mean = c()

for (i in 1:length(timepoints)){
  names_norm_mean=rbind(names_norm_mean,paste0("mean abundance at ", toString(timepoints[i]), "h"))
}

for (i in 1:nrow(measurements)) {
  # make data and annot vectors; this needs to be adjusted to data
  if (measurements$dump_row[i]!=1){
  ithrow=(measurements[i,1:length(sample_names)])
  split_row=matrix(NA, nrow=length(timepoints), ncol=max(na.omit(number_of_replicates)))
  reps=matrix(NA, nrow=length(timepoints), ncol=1)
  
  for (j in 1:length(timepoints)){
    split_row[j, 1:number_of_replicates[j]] = as.numeric(ithrow[NAPositions_startstop[j,1]:NAPositions_startstop[j,2]])
    reps[j]=length(split_row[j,is.na(split_row[j,])==FALSE])
    norm_mean[i,j] = mean(na.omit(split_row[j,]))
  }
  
  group.sizes=t(reps)
  
  # now run JTK_CYCLE
  
  options(stringsAsFactors=FALSE)
  data=measurements[i,1:length(sample_names)]
  
  data=data[,is.na(data)==FALSE]
  data=as.data.frame(data)
  annot=as.data.frame(row.names(measurements)[i])
  names(annot)="x"
  annot=rbind(annot, annot, annot)
  annot[2,1]='dummy1'
  annot[3,1]='dummy2'
  data=rbind(data, data, data)
  row.names(data)[2]='dummy1'
  row.names(data)[3]='dummy2'
  
  write.table(data, file=data_filename, row.names=TRUE, sep="\t", quote = FALSE)
  data <- read.delim(data_filename)
  
  write.table(annot, file=annot_filename, row.names=FALSE, sep="\t", quote = FALSE)
  annot <- read.delim(annot_filename)
  
  jtkdist(length(group.sizes),group.sizes) # changed this from originally provided file to accomodate different numbers of replicates
  
  periods <- 5.5:6.5 # i.e. looks for periods between 22 and 26h
  jtk.init(periods,4)  # changed this to 4 because every 4h
  
  cat("JTK analysis started on",date(),"\n")
  flush.console()
  
  st <- system.time({
    res <- apply(data,1,function(z) {
      jtkx(z)
      c(JTK.ADJP,JTK.PERIOD,JTK.LAG,JTK.AMP)
    })
    res <- as.data.frame(t(res))
    bhq <- p.adjust(unlist(res[,1]),"BH")
    res <- cbind(bhq,res)
    colnames(res) <- c("BH.Q","ADJ.P","PER","LAG","AMP")
    results <- cbind(annot,res,data)
    results <- results[order(res$ADJ.P,-res$AMP),]
  })
  print(st)
  
  
  # store results in separate dataframe called allresults
  
  allresults[i,]=results[1,1:6]
  }
}

names(allresults)=c('Peptide ID','BH.Q','ADJ.P','PER','LAG','AMP')
# where dump_row=1, add NA to pvalue and qvalue
for (i in 1:nrow(measurements)){
  if (measurements$dump_row[i]==1){
    is.na(allresults$"ADJ.P"[i]) = TRUE    
  }
}

allresults$"BH.Q" = p.adjust(allresults$"ADJ.P", method="BH")

# if timecourse goes either up or down in a straight line over the entire timecourse, add 
# value 1 to o and q values

straightlines = 0 # straightline counts in how many cases a timecourse goes just up or just down

for (i in 1:nrow(allresults)){
  if (measurements$dump_row[i] != 1){
    down = 0
    up = 0
    for (j in 2:length(timepoints)){
      if (norm_mean[i,j] < norm_mean[i,j-1]){down = down + 1}
      if (norm_mean[i,j] > norm_mean[i,j-1]){up = up + 1}
    }
    if (down == length(timepoints)-1 | up == length(timepoints)-1){
      allresults$"ADJ.P"[i] = allresults$"ADJ.P"[i] + 1
      allresults$"BH.Q"[i] = allresults$"BH.Q"[i] + 1
      straightlines=straightlines + 1
    }
  }
}


# Determine maximum and minimum time points and their values

minmax=as.data.frame(matrix(NA, nrow=nrow(measurements), ncol=5))
names(minmax)=c("Peak timepoint","Values at peak","Trough timepoint","Values at trough","Fold change")

for (i in 1:nrow(measurements)){
  if (measurements$"dump_row"[i] != 1){
    minmax$"Values at peak"[i]=max(na.omit(norm_mean[i,]))
    minmax$"Values at trough"[i]=min(na.omit(norm_mean[i,]))
    minmax$"Peak timepoint"[i]=timepoints[which(norm_mean[i,]==max(na.omit(norm_mean[i,])))]
    minmax$"Trough timepoint"[i]=timepoints[which(norm_mean[i,]==min(na.omit(norm_mean[i,])))]
    if (is.na(minmax$"Values at peak"[i]) == FALSE & is.na(minmax$"Values at trough"[i]) == FALSE ){
      minmax$"Fold change"[i]=minmax$"Values at peak"[i]/minmax$"Values at trough"[i]
    }
  }
}

names(norm_mean)=names_norm_mean

measurements_extended=cbind(PData$Sequence, PData$Accession, PData$Description, PData$Progenesis, allresults, minmax, norm_mean, measurements)
names(measurements_extended)[1:4]=c("Sequence", "Accession", "Description", "Modifications")

# make table to count peak times
peaktime_counts_table=as.data.frame(matrix(NA, nrow=length(timepoints), ncol=7))
names(peaktime_counts_table) = c("Timepoint","total number of peaking IDs","total number of IDs at trough","number of peaking significant (AD.P) IDs","number of significant (ADJ.P) IDs at trough", "number of peaking significant (BH.Q) IDs","number of significant (BH.Q) IDs at trough")
peaktime_counts_table[,1] = timepoints
# make table with p and q values and peak and trough timepoints

temp_peak_table=as.data.frame(cbind(allresults$"BH.Q", allresults$"ADJ.P",minmax$"Peak timepoint", minmax$"Trough timepoint"))
names(temp_peak_table)=c("BH.Q", "ADJ.P", "Peak timepoint", "Trough timepoint")
row.names(temp_peak_table)=row.names(measurements)

for (i in 1:length(timepoints)){
  peaktime_counts_table$"total number of peaking IDs"[i] = length(temp_peak_table$"Peak timepoint"[temp_peak_table$"Peak timepoint"==timepoints[i] & is.na(temp_peak_table$"Peak timepoint")==FALSE])
  peaktime_counts_table$"total number of IDs at trough"[i] = length(temp_peak_table$"Trough timepoint"[temp_peak_table$"Trough timepoint"==timepoints[i] & is.na(temp_peak_table$"Trough timepoint")==FALSE])
  peaktime_counts_table$"number of peaking significant (AD.P) IDs"[i] = nrow(temp_peak_table[temp_peak_table$"ADJ.P" < 0.05 & is.na(temp_peak_table$"ADJ.P")==FALSE & temp_peak_table$"Peak timepoint"==timepoints[i],])
  peaktime_counts_table$"number of significant (ADJ.P) IDs at trough"[i] = nrow(temp_peak_table[temp_peak_table$"ADJ.P" < 0.05 & is.na(temp_peak_table$"ADJ.P")==FALSE & as.numeric(temp_peak_table$"Trough timepoint")==as.numeric(timepoints[i]),])
  peaktime_counts_table$"number of peaking significant (BH.Q) IDs"[i] = nrow(temp_peak_table[temp_peak_table$"BH.Q" < 0.05 & is.na(temp_peak_table$"BH.Q")==FALSE & temp_peak_table$"Peak timepoint"==timepoints[i],])
  peaktime_counts_table$"number of significant (BH.Q) IDs at trough"[i] = nrow(temp_peak_table[temp_peak_table$"BH.Q" < 0.05 & is.na(temp_peak_table$"BH.Q")==FALSE & temp_peak_table$"Trough timepoint"==timepoints[i],])
}

# making sheet for counts:
counting_info=data.frame(matrix(nrow=6 + length(timepoints), ncol=10))
counting_info[1,1] = "Number of identifications"
counting_info[1,2] = nrow(measurements)
counting_info[2,1] = "Number of identifications with JTK pvalue determined"
counting_info[2,2] = nrow(measurements)-sum(dump_row)
counting_info[3,1] = "Number JTK significant (p<0.05), unadjusted"
counting_info[3,2] = length(allresults$"ADJ.P"[allresults$"ADJ.P"<0.05 & is.na(allresults$"ADJ.P")==FALSE])
counting_info[4,1] = "Number JTK significant (p<0.05), BH adjusted"
counting_info[4,2] = length(allresults$"BH.Q"[allresults$"BH.Q"<0.05 & is.na(allresults$"ADJ.P")==FALSE])
counting_info[5,1] = "Number JTK and BH adjusted q<0.15"
counting_info[5,2] = length(allresults$"BH.Q"[allresults$"BH.Q"<0.15 & is.na(allresults$"BH.Q")==FALSE])

counting_info[6,1] = "Number JTK significant (p<0.05), unadjusted but straight line"
counting_info[6,2] = length(allresults$"ADJ.P"[allresults$"ADJ.P" > 1 & allresults$"ADJ.P" < 1.05 & is.na(allresults$"ADJ.P")==FALSE])
counting_info[7,1] = "Number JTK significant (q<0.05), but straight line"
counting_info[7,2] = length(allresults$"BH.Q"[allresults$"BH.Q" > 1 & allresults$"BH.Q" < 1.05 & is.na(allresults$"BH.Q")==FALSE])
counting_info[8,1] = "Number JTK q<0.15, but straight line"
counting_info[8,2] = length(allresults$"BH.Q"[allresults$"BH.Q" > 1 & allresults$"BH.Q" < 1.15 & is.na(allresults$"BH.Q")==FALSE])


counting_info[1:length(timepoints),4:10] = peaktime_counts_table

names(counting_info) = c("Description of ID groups","number of IDs","Counts of peaks and troughs per timepoint:","Timepoint","total number of peaking IDs","total number of IDs at trough","number of peaking significant (AD.P) IDs","number of significant (ADJ.P) IDs at trough","number of peaking significant (BH.Q) IDs","number of significant (BH.Q) IDs at trough")

for (i in 1:nrow(counting_info)){
  for (j in 1:ncol(counting_info)){
    if (is.na(counting_info[i,j])==TRUE){
      counting_info[i,j]=""
    } 
  }
}

sheets=c("PData","measurements_extended","counting_info")
sheetNames=c("Original input data","JTK_CYCLE and other results","Summary of numbers")
WriteXLS(sheets, ExcelFileName=JTK_output_filename,  SheetNames = sheetNames, row.names=TRUE)
```
