## Supplementary Data S6 for "The circadian clock gene circuit controls protein and phosphoprotein rhythms in *Arabidopsis thaliana*": topGO_analysis.R

```
# reference to topGO:
# Alexa, A., Rahnenf�hrer, J. & Lengauer, T. Improved scoring of functional groups from gene expression data by decorrelating GO graph structure. Bioinformatics 22, 1600-1607 (2006).


# on phospho I wt data, using pvalues from JTK_CYCLE analysis and by peak time point groups

# allGenes: needs to be a list of indices of background accessions referring to GlobalProteindata. Here: background_indices.
# then allGenes also needs for each index the corresponding accession. That is what names(background_indices does)

# genSel is a function processed for each value in allGenes (i.e. each accession that is referred to by allGenes)
     # and then does one of more operations on that row in GlobalProteindata that alltogether return either
     # true or false for each accession

##prior to running this script install topGO
#source("http://bioconductor.org/biocLite.R")
#biocLite("topGO")
library("topGO")
library(WriteXLS)

# user needs to set path

GlobalProteindata = read.table('topGO_input.csv', skip=0, header=1, sep=",", quote="\"")

row.names(GlobalProteindata)=GlobalProteindata$Peptide.ID
GlobalProteindata$pvalues = GlobalProteindata$ADJ.P


#############################
# remove rows with NA in WT_JTK_pvalue
#############################

GlobalProteindata = GlobalProteindata[GlobalProteindata$pvalues != "#N/A",]
GlobalProteindata = GlobalProteindata[is.na(GlobalProteindata$pvalues) == FALSE,]


#############################
# remove entries where another phosphopeptides has same accession and same pvalue
#############################

# make unique list of accessions

AllAccessions=as.vector(unique(GlobalProteindata$Accession))
GlobalProteindata$"Delete_if_1"=rep(0,nrow(GlobalProteindata))

for (i in 1:length(AllAccessions)){
 ithAccession=AllAccessions[i]
  # find all rows with this accession
 # initialize dataframe that is going to contain the peptide IDs of the same accession and their pvalue
 InfoIthAccession=as.data.frame(matrix(data=NA,nrow=62,ncol=4)) # number of rows set to 10 which should be plenty (change if necessary)
 names(InfoIthAccession)=c("PeptideID","Accession","pvalue","Delete_if_1") 
 InfoIthAccession$"Delete_if_1"=rep(0,62)
 rowcounter=1
   for (j in 1:nrow(GlobalProteindata)){
     
     # make data frame with all PeptideIDs and their pvalues for the current accession
     if (as.vector(GlobalProteindata$Accession[[j]])==ithAccession){
       InfoIthAccession[rowcounter,"PeptideID"]=as.vector(GlobalProteindata$"Peptide.ID"[j])
       InfoIthAccession[rowcounter,"Accession"]=as.vector(GlobalProteindata$"Accession"[j])
       InfoIthAccession[rowcounter,"pvalue"]=as.vector(GlobalProteindata$"pvalues"[j])
       rowcounter=rowcounter+1
     }
   }
   # remove unnecessary rows in InfoIthAccession
   InfoIthAccession=InfoIthAccession[is.na(InfoIthAccession$PeptideID)==FALSE,]
   # find out if there are duplicate pvalues
      # find how many duplicates there must be
    NumberOfDuplicates=length(InfoIthAccession$pvalue) - length(unique(InfoIthAccession$pvalue))
     if(NumberOfDuplicates > 0){
       t=nrow(InfoIthAccession)-1
       for (k in 1:t){
         kthPeptideID=InfoIthAccession[k,"PeptideID"]
         kthPvalue=InfoIthAccession[k,"pvalue"]
         a=k+1
         b=nrow(InfoIthAccession)
         for (l in a:b){
           if(InfoIthAccession[l,"pvalue"]==kthPvalue & InfoIthAccession[l,"Delete_if_1"] == 0){
             remove_PeptideID=InfoIthAccession[l,"PeptideID"]
             InfoIthAccession[l,"Delete_if_1"]=1
             GlobalProteindata[remove_PeptideID,"Delete_if_1"]=1 # need to activate when finished testing
            }
           
         }

       }
       
     }
    
    
  }
  

GlobalProteindata = GlobalProteindata[GlobalProteindata$"Delete_if_1" == 0,]
GlobalProteindata$"pvalues"[GlobalProteindata$"pvalues"== '#N/A']  = NA

pvaluesvector = as.vector(GlobalProteindata$"pvalues")

GlobalProteindata$"pvalues" = pvaluesvector

#############################
## Remove all rows where accession has several phosphopeptides and peptide of the row is not the lowest bpvalue
#############################


bestPValue <- function(gene, pvalue="pvalues")
{  
  return( min(GlobalProteindata[GlobalProteindata[["Accession"]]==gene,pvalue]))
}


background_indices_temp <- sapply(unique(GlobalProteindata$Accession), function(x) {bestPValue(x)})
names(background_indices_temp) = unique(GlobalProteindata$Accession)

GlobalProteindata$ZeroThenRemove=rep(0, (length(GlobalProteindata$Peptide.ID)))

# for all peptide IDs in GlobalProteinsdata, set the ZerThenRemove field to 1 if the peptide's
# pvalue is the one that was selected as the best pvalue for this protein:

for(Peptide in GlobalProteindata$Peptide.ID)
{
  i=which(GlobalProteindata$Peptide.ID == Peptide)
  protein = as.character(GlobalProteindata$Accession[i])
  if (GlobalProteindata[[Peptide,"pvalues"]] == background_indices_temp[[protein]])
  {
    GlobalProteindata[Peptide,"ZeroThenRemove"] = 1
  }
}


######### Get actual indices that have to be removed ($IndicesToRemove is not a good name, is just a 1 and 0 column, 1 has to be kept, 0 to be removed)

KeepIndex=which(GlobalProteindata$ZeroThenRemove==1)

######### Make new, unique dataframe from GlobalProteindata that only has the unique entries with lowest p values

GlobalProteindata_unique = GlobalProteindata[KeepIndex,]
GlobalProteindata_unique$OldIndex = KeepIndex


###############################################################################
# GO analysis on peak / trough and time point
################################################################################

measurements = GlobalProteindata_unique[grep("(^X3702_PhI_wt).+[^\\.][0-9]$", names(GlobalProteindata_unique))] 
row.names(measurements) <- GlobalProteindata_unique[,"Peptide.ID"]
measurementsUn = measurements

Hours_wt <- factor(c(12,  12,  12,  12,  16,  16,  16,  16,  20,  20,  20,  20,  20,  24,  24,  24,  24,  28,  28,  28,  28,  32,  32,  32))
Hours = Hours_wt

measurements[measurements==0] <- NA
measurementsUn[measurementsUn==0] = NA
measurementsArchsinh <- asinh(measurements)
meanMeasurement <- as.data.frame(t(apply(measurementsArchsinh, 1, function(x) c(by(x, Hours, function(x) mean(t(na.omit(x))))))))
meanMeasurementUn <- as.data.frame(t(apply(measurementsUn, 1, function(x) c(by(x, Hours, function(x) mean(t(na.omit(x))))))))

# determine for each row, which time point best peptide peaks
GlobalProteindata_unique$peak = sapply(apply(meanMeasurement, 1, function(x) { as.numeric(names(meanMeasurement)[which(x==max(na.omit(x)))])}), function(x) {x[1]})
GlobalProteindata_unique$trough = sapply(apply(meanMeasurement, 1, function(x) { as.numeric(names(meanMeasurement)[which(x==min(na.omit(x)))])}), function(x) {x[1]})

row.names(GlobalProteindata_unique)=GlobalProteindata_unique$Accession

# calculate maximum and minimum

GlobalProteindata_unique$Maxi = apply(meanMeasurementUn, 1, function(x) {max(na.omit(x))})

GlobalProteindata_unique$Mini = apply(meanMeasurementUn, 1, function(x) {min(na.omit(x))})

# matrix of extrema to pass into applying fold change

c=matrix(c(GlobalProteindata_unique$Maxi, GlobalProteindata_unique$Mini), nrow=length(GlobalProteindata_unique$Maxi), ncol=2)

# calculate fold change; note: there is a foldchange column in the file already which was calculated in excel
# but this does not deal with zeros and missing values properly

GlobalProteindata_unique$foldchange_n = apply(c, 1, function(x) {x[1]/x[2]})


# calculate BH adjusted pvalues
GlobalProteindata_unique$pvaluesadjust_n=p.adjust(GlobalProteindata_unique$pvalues, method="BH")

# initialize list of sheet names for excel output file
sheetNames = list()


# read in annotations file
annotations = read.table("gene_association.csv", skip=24,header=0, sep=",", quote="\"")
names(annotations) <- c("DB","DB Object ID","DB Object Symbol","Qualifier","GO ID","DB:Reference","Evidence Code","With (or) From","Aspect","DB Object Name","DB Object Synonym","DB Object Type","Taxon(|taxon)","Date","Assigned By","Annotation Extension","Gene Product Form ID")

#ditch anything that's not a protein annotation
annotations = annotations[annotations$"DB Object Type" == "protein",]

# makes tables of DB Object Name (which is the ATG accession) and GO ID for each of the 3 categories
functions = annotations[annotations$"Aspect" == "F",][,c("DB Object Name", "GO ID")]
processes = annotations[annotations$"Aspect" == "P",][,c("DB Object Name", "GO ID")]
components = annotations[annotations$"Aspect" == "C",][,c("DB Object Name", "GO ID")]

#function to collect go terms for a gene

grabGOTerms <- function(x, dataframe, fieldGene="DB Object Name", fieldGO="GO ID") {
  index = apply(dataframe, 1, function(y) {grepl(y[fieldGene], x, fixed = TRUE)})
  return(unique(as.character(dataframe[index,fieldGO])))
}

#define our annotation universe which we enrich against

background_indices = which(GlobalProteindata_unique$ZeroThenRemove==1)
names(background_indices) = GlobalProteindata_unique[GlobalProteindata_unique$ZeroThenRemove==1,"Accession"]

# all (unique) observed accessions in dataset:
observedProteins <- unique(names(background_indices))

# for each of the 3 GO categories, make list of GO terms for each protein and name entries by the accessions
gene2GOMF <- lapply(observedProteins, function(x) grabGOTerms(x, functions))
names(gene2GOMF) = observedProteins

gene2GOMF_df = as.data.frame(matrix(nrow = length(gene2GOMF), ncol = 1000))

# following code puts the GO term list into a data frame for putting into file later
maxlength = 0
for (i in 1:length(gene2GOMF)){
  gene2GOMF_df[i,1] = names(gene2GOMF)[i]
  GOs = as.data.frame(gene2GOMF[i])
  GONumber = nrow(GOs)
  if (GONumber > maxlength){
    maxlength = GONumber
  }
  for (j in 1:GONumber){
    gene2GOMF_df[i,j+1] = as.character(GOs[j,1])
  }
}

gene2GOMF_df = gene2GOMF_df[,1:(maxlength + 1)]

gene2GOBP <- lapply(observedProteins, function(x) grabGOTerms(x, processes))
names(gene2GOBP) = observedProteins

gene2GOBP_df = as.data.frame(matrix(nrow = length(gene2GOBP), ncol = 1000))

maxlength = 0
for (i in 1:length(gene2GOBP)){
  gene2GOBP_df[i,1] = names(gene2GOBP)[i]
  GOs = as.data.frame(gene2GOBP[i])
  GONumber = nrow(GOs)
  if (GONumber > maxlength){
    maxlength = GONumber
  }
  for (j in 1:GONumber){
    gene2GOBP_df[i,j+1] = as.character(GOs[j,1])
  }
}

gene2GOBP_df = gene2GOBP_df[,1:(maxlength + 1)]

gene2GOCC <- lapply(observedProteins, function(x) grabGOTerms(x, components))
names(gene2GOCC) = observedProteins

gene2GOCC_df = as.data.frame(matrix(nrow = length(gene2GOCC), ncol = 1000))

maxlength = 0
for (i in 1:length(gene2GOCC)){
  gene2GOCC_df[i,1] = names(gene2GOCC)[i]
  GOs = as.data.frame(gene2GOCC[i])
  GONumber = nrow(GOs)
  if (GONumber > maxlength){
    maxlength = GONumber
  }
  for (j in 1:GONumber){
    gene2GOCC_df[i,j+1] = as.character(GOs[j,1])
  }
}

gene2GOCC_df = gene2GOCC_df[,1:(maxlength + 1)]


for (hour in c(12, 16, 20, 24, 28, 32)) {
geneSel = function(x) {return(!is.na(GlobalProteindata_unique[x,"pvalues"])& GlobalProteindata_unique[x,"pvalues"] <= 0.05
                              & !is.na(GlobalProteindata_unique[x, "peak"])& GlobalProteindata_unique[x, "peak"] == hour)}


typeName = "_peak_at_"
typeOf = "peak"
pvalueType = "pvaluesadjustBH"

###############
#Molecular Function
###############
  name = paste("MF", typeName, hour, sep="")
  sheetNames=c(sheetNames,name)
  go_data <- new("topGOdata",
                 description = "Enrichment test", "MF",
                 allGenes = na.omit(background_indices), geneSel = geneSel,
                nodeSize = 3, annot = annFUN.gene2GO, gene2GO = gene2GOMF)
                  
  resultFisher.weight01 <- runTest(go_data, algorithm = "weight01", statistic = "fisher")
  
  resultsMF <- GenTable(go_data, resultFisher= resultFisher.weight01, orderBy = "resultFisher", topNodes=geneData(resultFisher.weight01)["SigTerms"])
names(resultsMF)[6] = 'resultFisher'  
eval(parse(text=paste(name, "<-", "resultsMF", sep="")))
  

###############
#Biological Process
###############

name = paste("BP", typeName, hour, sep="")
sheetNames=c(sheetNames,name)
go_data <- new("topGOdata",
               description = "Enrichment test", "BP",
               allGenes = na.omit(background_indices), geneSel = geneSel,
               nodeSize = 3, annot = annFUN.gene2GO, gene2GO = gene2GOBP)

resultFisher.weight01 <- runTest(go_data, algorithm = "weight01", statistic = "fisher")

resultsBP <- GenTable(go_data, resultFisher= resultFisher.weight01, orderBy = "resultFisher", topNodes=geneData(resultFisher.weight01)["SigTerms"])
names(resultsBP)[6] = 'resultFisher'
eval(parse(text=paste(name, "<-", "resultsBP", sep="")))


###############
#Cellular Component
###############
name = paste("CC", typeName, hour, sep="")
sheetNames=c(sheetNames,name)
go_data <- new("topGOdata",
               description = "Enrichment test", "CC",
               allGenes = na.omit(background_indices), geneSel = geneSel,
               nodeSize = 3, annot = annFUN.gene2GO, gene2GO = gene2GOCC)

resultFisher.weight01 <- runTest(go_data, algorithm = "weight01", statistic = "fisher")

resultsCC <- GenTable(go_data, resultFisher= resultFisher.weight01, orderBy = "resultFisher", topNodes=geneData(resultFisher.weight01)["SigTerms"])
names(resultsCC)[6] = 'resultFisher'
eval(parse(text=paste(name, "<-", "resultsCC", sep="")))

}


new_sheetnames = matrix(nrow = length(sheetNames), ncol = 1)
for (i in 1:length(sheetNames)){
  new_sheetnames[i] = sheetNames[[i]]  
}


WriteXLS(new_sheetnames, ExcelFileName = "GO_output_enrichment.xlsx",
         SheetNames = sheetNames)

# also save the GO terms for each observed protein:

sheet_names_gene2GO = c('gene2GOMF_df','gene2GOBP_df','gene2GOCC_df')
WriteXLS(sheet_names_gene2GO, ExcelFileName = "GO_output_terms.xlsx",
         SheetNames = sheet_names_gene2GO)
```
