## Supplementary Figures, Supplementary Tables, Supplementary Methods for "The circadian clock gene circuit controls protein and phosphoprotein rhythms in *Arabidopsis thaliana*"

**Supplementary material included inthis document:**

- Supplementary Tables S1 to S4
- Supplementary Figures S1 to 8
- Supplementary Methods

**Supplementary material included as separate files:**

- Supplementary Data S1 to S6

**Supplementary tables**

Table S 1: Number of rhythmic WT time courses (JTK_CYCLE p-value $<0.05$) where the absolute difference in phosphopeptide or protein abundance between CCA1-OX and WT is larger either at dusk or at dawn.

<table>
<tr><th>Dataset and time points</th><th>|CCA1-OX - WT| larger at dusk (ZT12, ZT36)</th><th>|CCA1-OX - WT| larger at dawn (ZT24, ZT48)</th></tr>
<tr><td>Phospho I (ZT12, ZT24)</td><td>424</td><td>182</td></tr>
<tr><td>Phospho II (ZT36, ZT24)</td><td>73</td><td>27</td></tr>
<tr><td>Phospho II (ZT36, ZT48)</td><td>64</td><td>36</td></tr>
<tr><td>Global I (ZT12, ZT24)</td><td>110</td><td>61</td></tr>
<tr><td>Global II (ZT36, ZT24)</td><td>21</td><td>24</td></tr>
<tr><td>Global II (ZT36, ZT48)</td><td>25</td><td>20</td></tr>
</table>

Table S 2: GPS3 kinase prediction, using ANOVA p-values. ANOVA analysis on each phosphosite time course was carried out on the data after outlier exclusion, treating 0 values as missing values. GPS3 predictions from Phosphopeptides with ANOVA p-values < 0.05 were used as foreground for testing enrichment of kinase predictions in all significantly changing phosphopeptides, while phosphopeptides with p<0.05 and a given peak time were used to analyse for enrichment at specific times of the day (24 or 48h). In the case of several predicted kinases within a family of kinases for the same phosphosite, only the one with the highest difference of score and cutoff was considered. Predictions for each kinase were counted in the foreground and the background and a Fisher's exact test was used to test for significant enrichment and significant p-values of over-represented groups are displayed in the table.

| | Phosphoproteomics dataset I | | | | | | | | Phosphoproteomics dataset II | | | | |
|---|---|---|---|---|---|---|---|---|---|---|---|---|---|
| | WT | | | | CCA1-Ox | | | | WT | | CCA1-Ox | | |
| | | peaks | | | | peaks | | | | peaks | | peaks | |
| EKPD kinase group | all p<0.05 | 12h | 24h | 28h | all p<0.05 | 23h | 24h | 28h | all p<0.05 | 24h | all p<0.05 | 24h | 48h |
| AGC | 0.0015 | | | | | | | | | | | | |
| AGC/PDK1 | 0.0000011 | | | | | | | 0.026 | | | | | |
| Atypical/TAF1 | | | | | | | | 0.0038 | 0.022 | | 0.0044 | | |
| CAMK | 0.00000072 | | | 0.022 | | | 0.034 | | 0.0013 | 0.001 | | 0.022 | |
| CAMK/CAMK-Unique | 0.000075 | | | | | | | | 0.014 | 0.0071 | | | |
| CAMK/CAMK1 | | | | | | | | | 0.036 | | | | |
| CAMK/CAMKL | 0.0036 | | | | | | | | | | | | |
| CMGC | 0.000026 | | | | | | | 0.048 | | 0.035 | | | |
| CMGC/CK2 | | | | | | | | 0.0048 | 0.011 | | 0.033 | | |
| CMGC/DYRK | 0.00619 | | | | | | | | | | | | |
| CMGC/GSK | 0.00024 | | | | | | | | | | | | |
| CMGC/MAPK | 0.00010 | 0.010 | | | | | | | | | | | |
| CMGC/CDK | 0.000026 | | | | | | | | | | | | |
| CK1/CK1 | 0.012 | | | | | | | | | | | | |
| STE/STE11 | 0.015 | | 0.020 | | | | | | | | 0.029 | | 0.039 |
| STE/STE7 | 0.003 | | | | | | | | | | | | |
| TK | | | | | 0.045 | | | | | | | | |
| TKL/IRAK | | | | | | | | | | | 0.0037 | | 0.0083 |
| Other/Haspin | | | | | | 0.00027 | | | | | | | |
| Other/WNK | | | | | | | 0.040 | | | | | | |
| Other/TTK | | | | | 0.045 | | | 0.026 | | | | | |

Table S 3: A selection of previously described SnRK1 regulated phosphosites that were also detected in our dataset. For (1), the listed peptides were were significantly lower in a line with downregulated SnRK1 activity, or significantly up-regulated in a SnRK1 over-expression line.

| Accession and name | reference | published peptide | peptide in this analysis | JTK_CYCLE p-values |
|---|---|---|---|---|
| AT1G77760, nitrate reductase 1 | (1) | SV**pS**SPFMN TASK | SV**pS**SPFMNTASK | dataset I:<br>WT: **3.6*10$^{-7}$**<br>CCA1-OX: **0.0045**<br>datasetII, JTK:<br>WT: 0.11<br>CCA1-OX: 1 |
| AT1G37130, nitrate reductase 2 | (1)<br>peptide ID 174 | VHDDDEDV (p)S(p)SEDE NETHNSNA VYYK | VHDDDEDV(p)S(p)SED ENETHNSNAVYYK | dataset I:<br>WT: **0.0009**<br>CCA1-OX: 0.23<br>datasetII, JTK:<br>WT: 0.0088<br>CCA1-OX: 1 |
| AT1G37130, nitrate reductase 2 | (1)<br>peptide ID 173 | SV**pS**TPFMN TTAK | SV**pS**TPFMNTTAK | dataset I:<br>WT: **2x10$^{-6}$**<br>CCA1-OX: 0.079<br>dataset II:<br>WT: 0.60<br>CCA1-OX: 1.0 |
| AT1G07110 F2KP | (2) | SVETL**pS**PF QQK | SVETL**pS**PFQQKDGQK | dataset 1:<br>WT: 1.0<br>CCA1-OX: 0.16<br>dataset2:<br>WT: **4.3x10$^{-8}$**<br>CCA1-OX: **0.0032** |

Table S 4: List of primers for point mutagenesis and subcloning of F2KP

| Primer name | Mutation | Sequence |
|---|---|---|
| AspF | S276D | ACCATCTTTCTGCTGAAACGGATCGAGTGTCTCCACAGACTTTGA |
| AspR | S276D | TCAAAGTCTGTGGAGACACTCGATCCGTTTCAGCAGAAAGATGGT |
| AlaF | S276A | CATCTTTCTGCTGAAACGGAGCGAGTGTCTCCACAGACTTT |
| AlaR | S276A | AAAGTCTGTGGAGACACTCGCTCCGTTTCAGCAGAAAGATG |
| F2KP-F | none | GAGCTTAAGATGGGGTCAGGTGCATCGAAGAATAC |
| F2KP-R | none | AGCTCTAGATCAGTCCATGAGTTTGTAGCGTTTCTCTTGC |

**Supplementary figures**

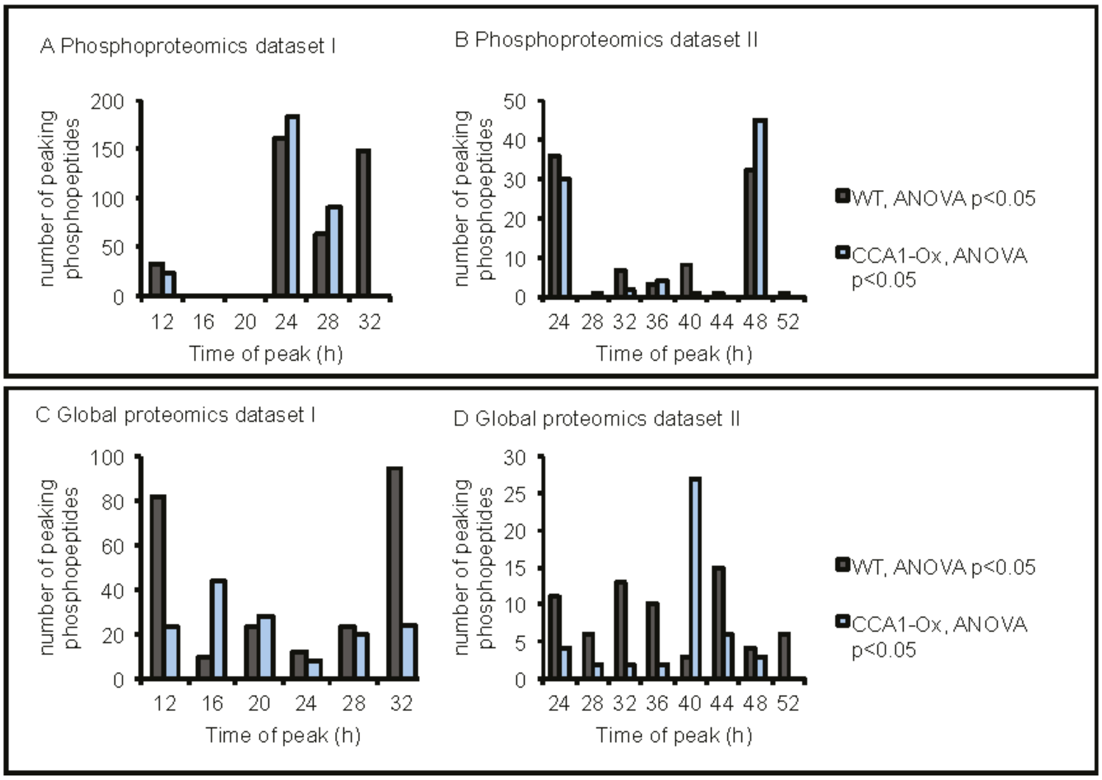


Figure S 1: Peak time distribution of changing (ANOVA p<0.05) phosphopeptides (A.B) and proteins (C,D) in datasets I (A,C) and II (B,D).

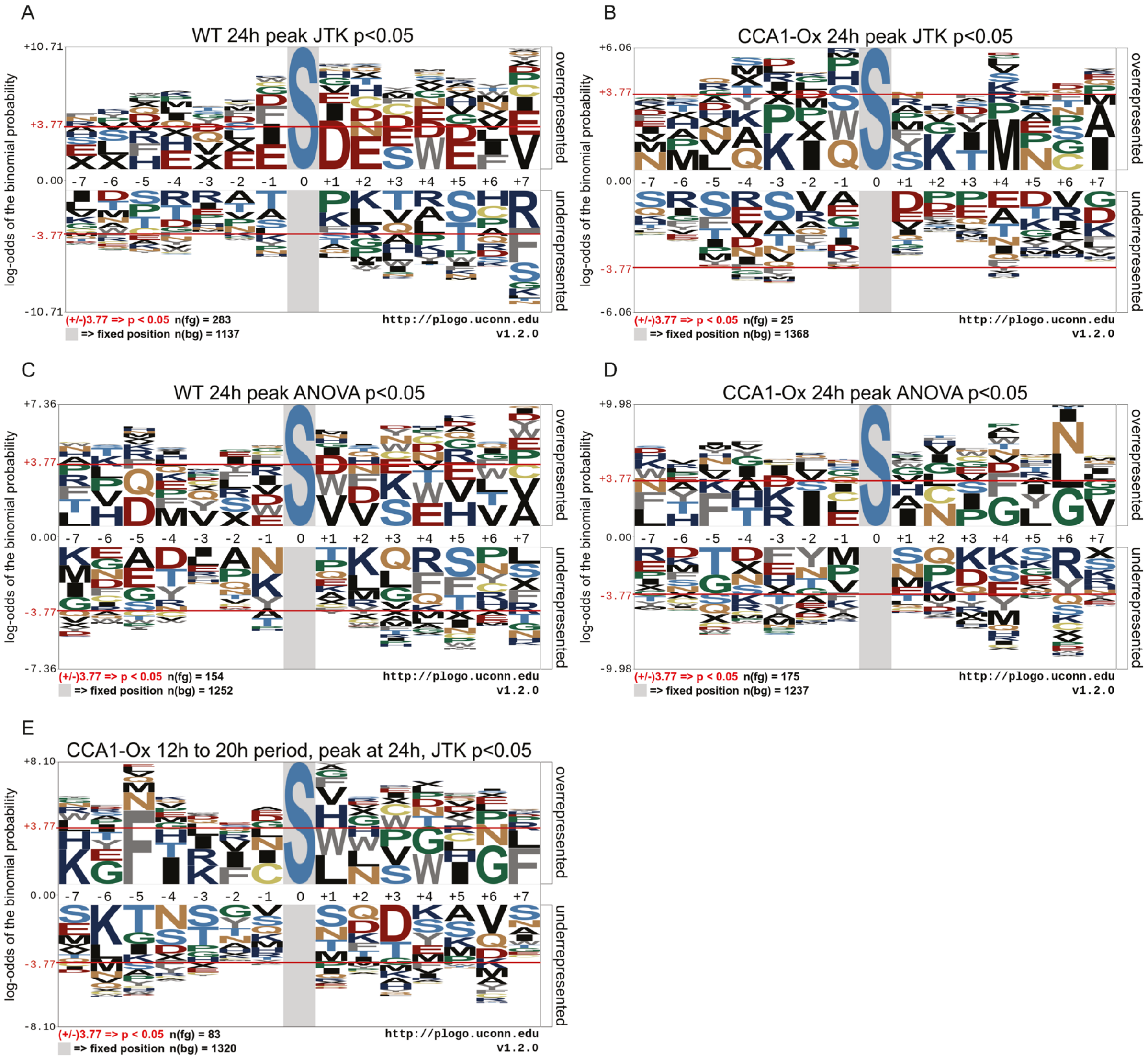


Figure S 2: pLogos generated for phosphoproteomics dataset I. (A,B) using peptides as foreground with JTK_CYCLE p<0.05 (allowed periods 22h to 26h) and peaking at 24h for (A) WT and (B) CCA1-OX, (C,D) using peptides as foreground with ANOVA p<0.05 and peaking at 24h for WT (C) and CCA1-OX (D). (E) CCA1-OX JTK_CYCLE p<0.05 when allowing periods 12h to 20h, and peaking at 24h.

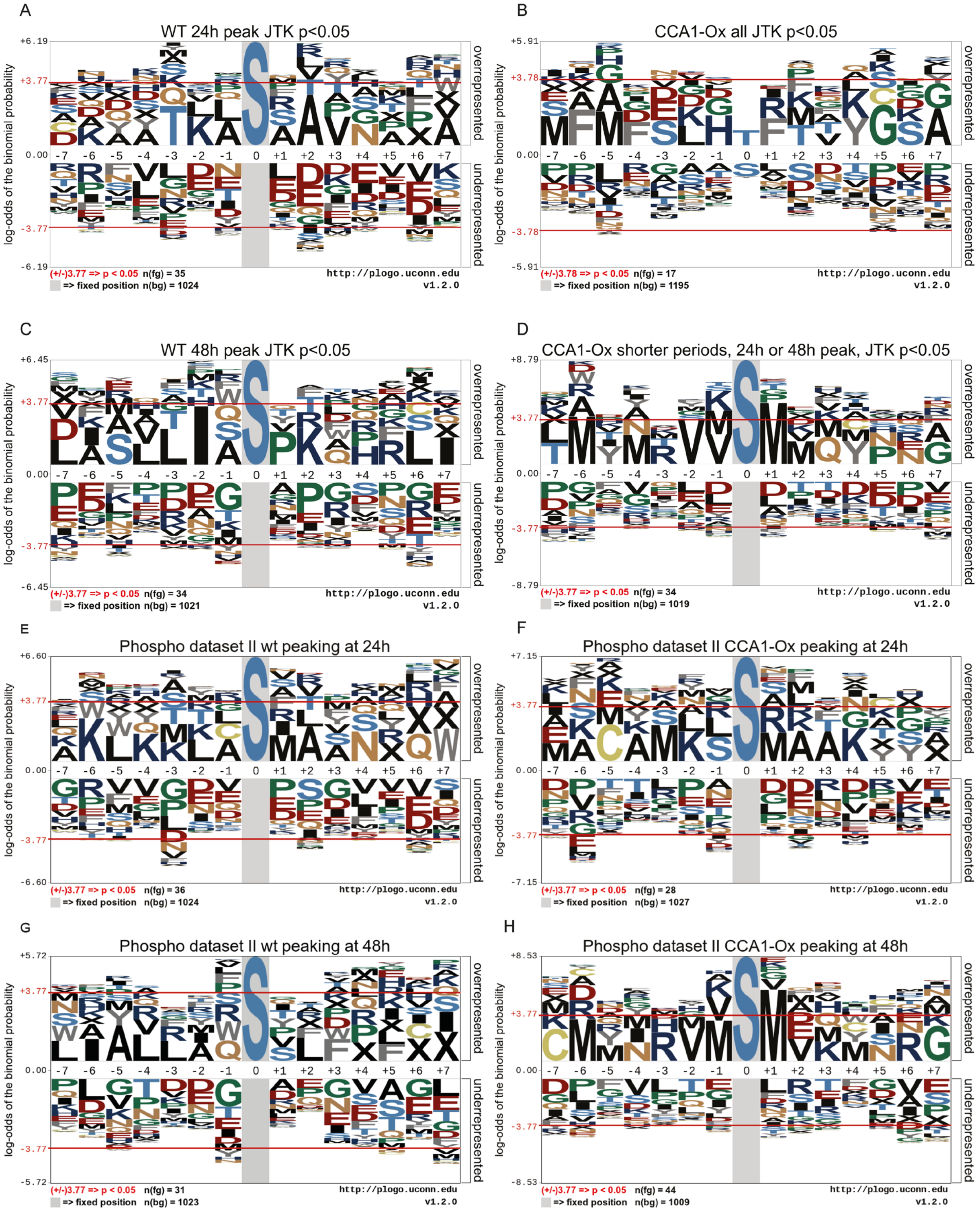


Figure S 3: pLogos generated for phosphoproteomics dataset II. (A,C) For the WT, phosphopeptides with JTK_CYCLE p-value <0.05 were chosen that peak at either 24h (A) or 48h (C). (B,D) For CCA1-OX phosphopeptides with JTK_CYCLE p-value <0.05 and peaking at 24h or 48h (identical to all significant phosphopeptides for (B)) and a period of 22 to 26h (B) or a period of 12h to 16h (D). (E,G) Like (A,C) but using ANOVA p-values <0.05 instead of JTK_CYCLE. (F,H) same as (E,G) for CCA1-OX.

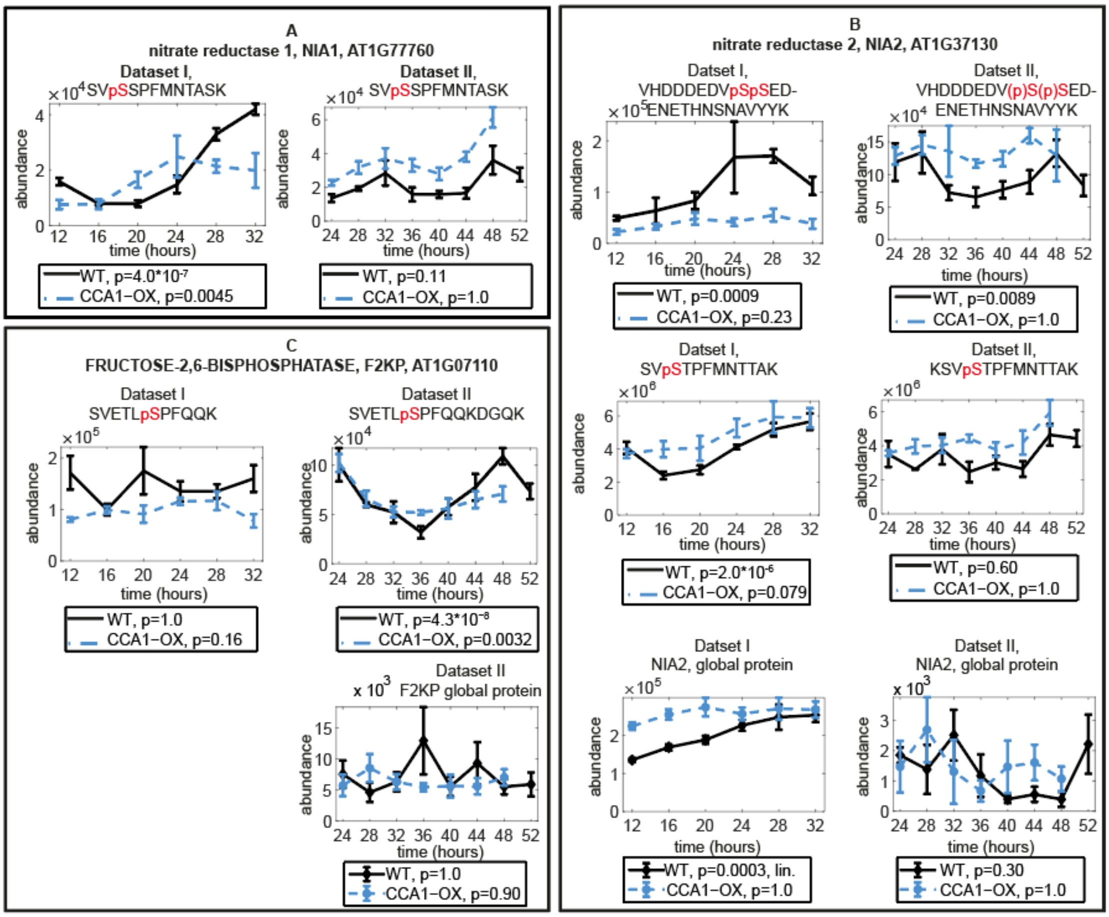


Figure S 4: Time courses of phosphopeptides that were previously described as SnRK1 regulated. (A) nitrate reductase NIA1, (B) NIA2, (C) the bifunctional enzyme fructose-6-phosphate 2-kinase/fructose-2,6-bisphosphatase, F2KP. Global protein data are shown where detected. p-values of JTK_CYCLE analysis for rhythmicity are shown under the plots. In the case of NIA2, one out of two serines (p in parentheses) is phosphorylated. NIA2 protein abundance was linear (’lin.’). See Table S3 for previous reports of these sites.

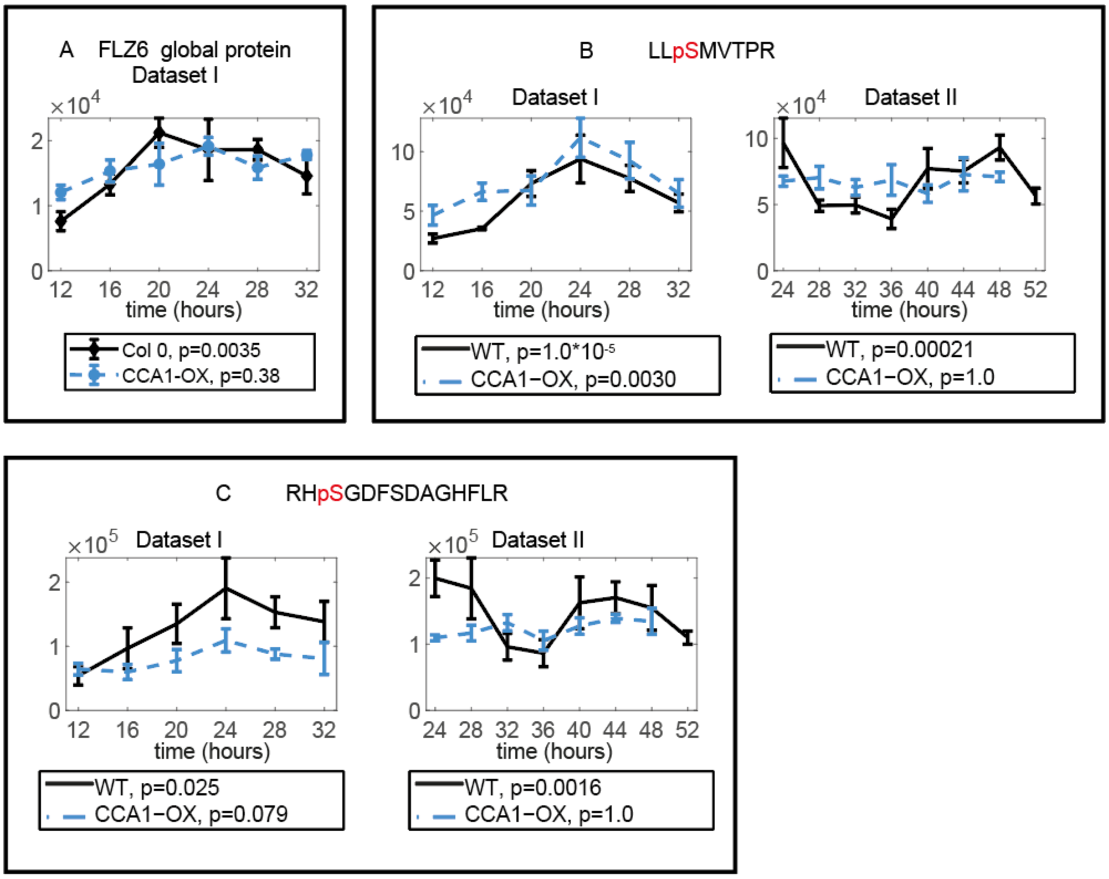


Figure S 5:Time courses of FCS LIKE ZINC FINGER (FLZ6, AT1G78020) identifications; (A) protein, (B,C) two phosphopeptides (B,C). p-values are shown under the plots.

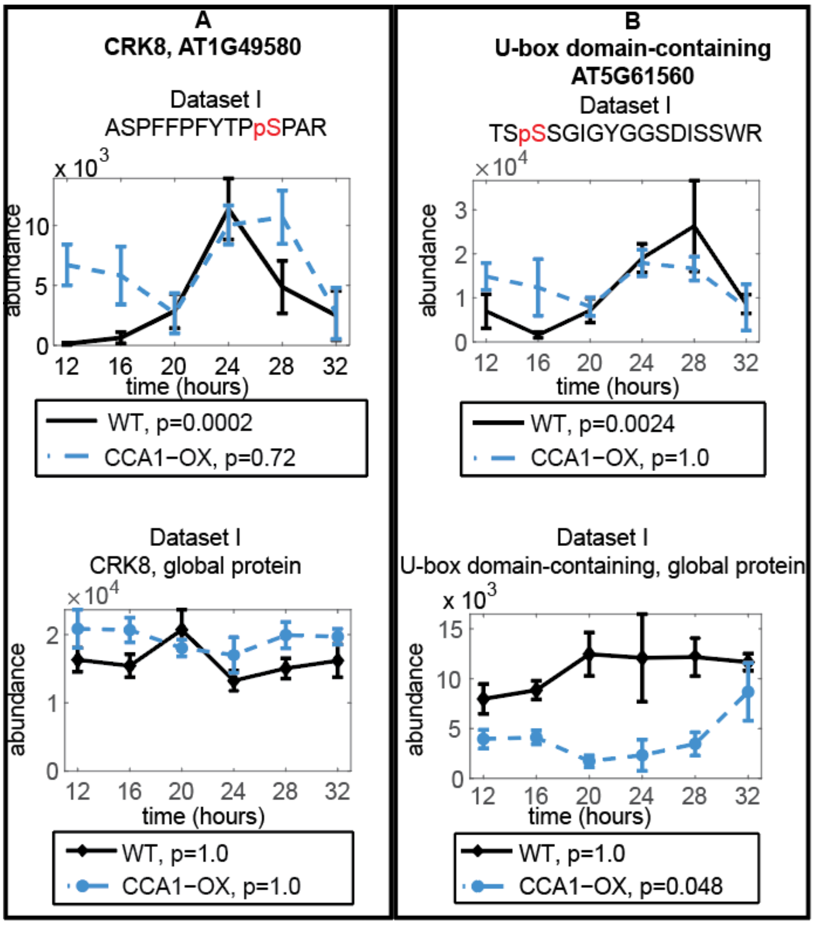


Figure S 6: Rhythmic kinases in dataset I: (A) CRK8, (B) U-box domain containing, profiles of phosphopeptides (top) and global protein (bottom). JTK_CYCLE p-values are given in the boxes under the graph. Phosphosites were not detected in datset II.

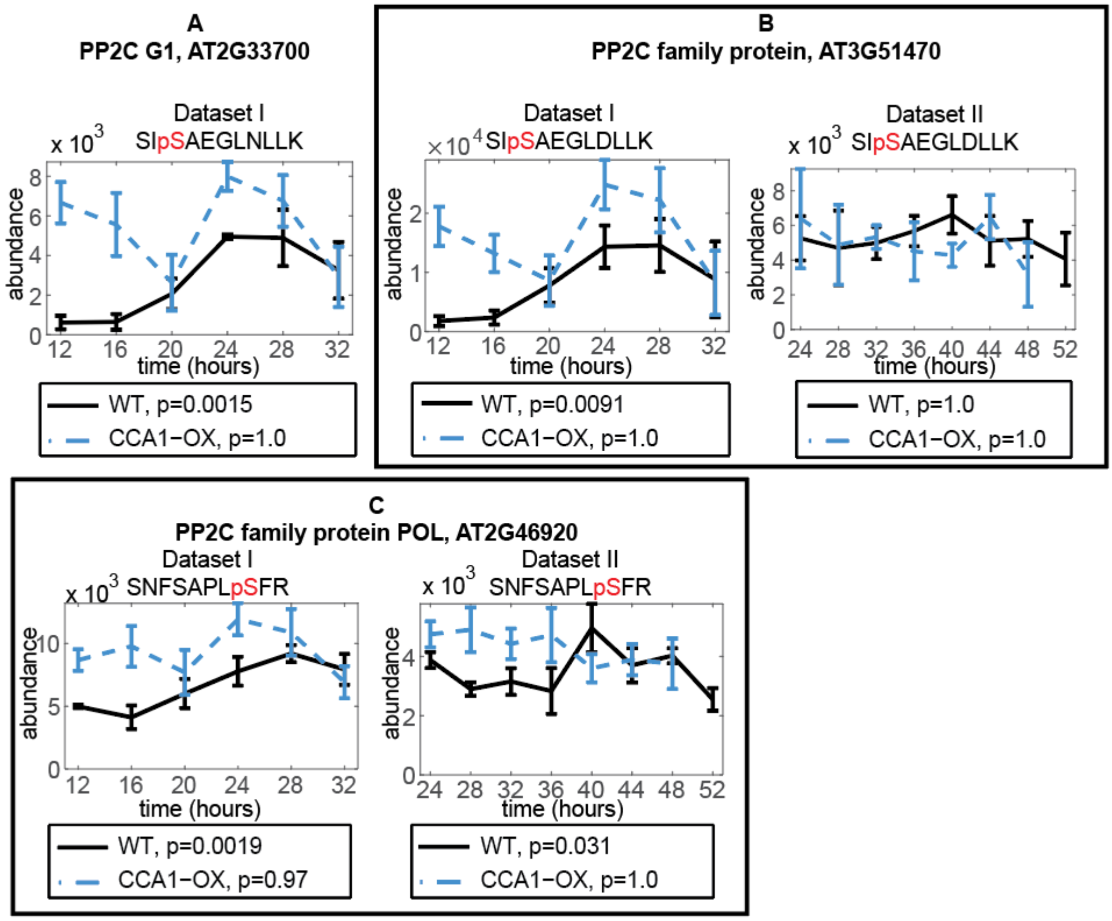


Figure S 7: Phosphosites of PP2C family phosphatases that are rhythmic in dataset I. (A) PP2C G1 / AT2G33700, (B) AT3G51470, (C) AT2G46920 / POL. JTK_CYCLE p-values are given in the boxes under the graph. No corresponding peptides detected in global protein analysis.

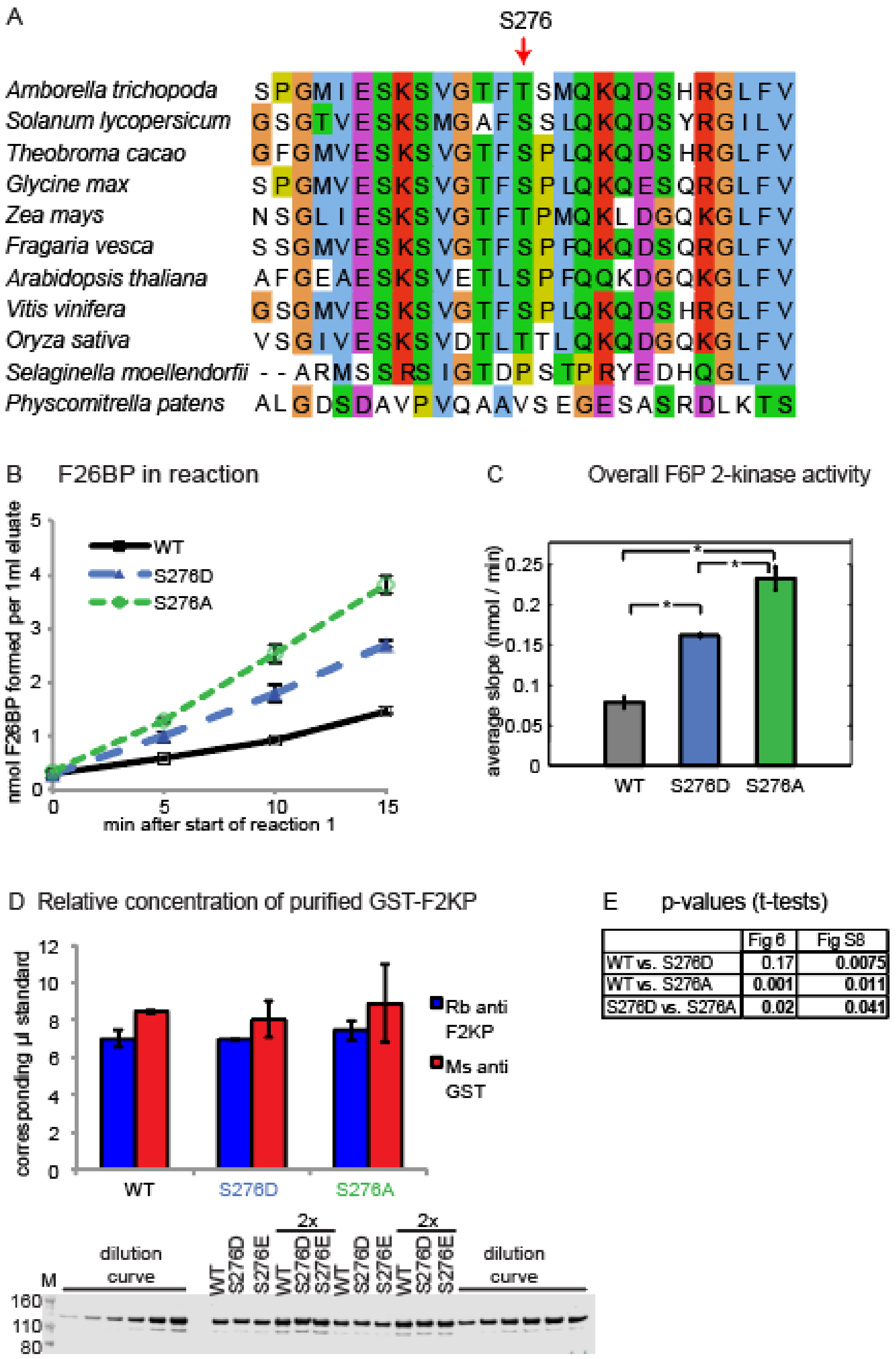


Figure S 8: Alignment of sequence surrounding F2KP Ser276 and *in vitro* GST-F2KP activity assay with WT and Ser276 point mutations. (A) Alignment of sequence area surrounding S276 in a selection of plant species. S276 is highlighted with a red arrow. Amino acid sequences of F2KP were aligned and viewed using the ClustalX2 and Jalview software (3,4). The *Arabidopsis thaliana* sequence was obtained from tair (www.arabidopsis.org, accession AT1G07110), the other sequences from ncbi (http://www.ncbi.nlm.nih.gov): XP_006843654.1 (*Amborella trichopoda*), XP_004242511.1 (*Solanum lycopersicum*), XP_007016919.1, (*Theobroma cacao*), XP_003552945.1 (*Glycine max*), ACF78476.1 (*Zea mays*), XP_004294306.1 (*Fragaria vesca*), XP_002276394.1 (*Vitis vinifera*),

NP_001054740.1 (*Oryza sativa* Japonica group), XP_002991023.1 (*Selaginella moellendorfii*), XP_001777124.1 (*Physcomitrella patens*). (B-D) Repeat experiment of F2KP in vitro activity measurements. B) F2KP activity at different time points of the reaction. (C) Activity calculated from slopes in (B). (D) relative quantification of GST-F2KP in eluates probed with rabbit (Rb) anti F2KP and mouse (Ms) anti GST. Original blot is shown below quantification for rabbit anti F2K. A dilution series of a sample mix was used for quantification, ranging from 0.5 to 1.5 loading equivalent of the samples. Averages of two dilution curves were used. (E) p-values of F2KP activity in Figure 6 and this figure. Error bars: SEM. * $p$-value $< 0.05$ in two-sided t-test.

**Supplementary methods**

*outlier analysis*

Two measures were used to identify statistical outliers on arcsinh transformed data: Pearson correlation analysis and PCA. Both were calculated in an R script included in Data S4. PCA was calculated within each time course and genotye. The Pearson correlation coefficient of peptide or protein abundance values of each replicate with the time point median was determined. For phosphopeptide data, all replicates in which the Pearson correlation coefficient was < 0.8 were regarded as outliers. The PCA analysis and plots of the difference to the sample median served to validate the exclusion of replicates. In the global data analysis of datase II, these measures revealed a machine drift and we regarded it as necessary to remove each first-run replicate of each time point.

These criteria resulted in removal of the following replicates as outliers:

Phospho I dataset – WT: 12D, 16B, 24A, 28B, 32A, 32B; CCA1.OX: 20D, 24E, 28C, 32C

Global I dataset: no outliers

Phospho II dataset – WT: 24E, 28D, 28F, 32A, 32D, 32H, 40B, 40F, 44D, 48F, 52F; CCA1-OX: 28E, 32D, 36E, 40D, 48E

Global II dataset – WT: 24A; 28A, 28B, 28F, 32A, 36A, 40A, 44A, 48F, 52A; CCA1-OX: 24A, 28A, 32A, 36A, 40A, 44A, 48A, 52A

All plots of PCA, Pearson correlation coefficient and boxplots of median of replicate minus time point mean are included in Supplementary Data S4.

*pLogo* generation

The online tool pLogo was used to visualize enrichment of amino acids at the 14 positions surrounding phosphorylated serine or threonine residues (5). A python script was used to generate lists of phosphosites for foreground and background with 7 amino acids on either side of the phosphorylated position.

As foreground, rhythmic (JTK_CYCLE $p<0.05$) or significantly changing (ANOVA $p<0.05$) were used, either the group or all significant phosphopeptides or the subgroup phosphopeptides that peak at 24h or 48h or both. The background consisted of all phosphopeptides detected in the respective experiment (dataset I or II) without the foreground.
